# Evaluation of Methods for Protein Representation Learning: A Quantitative Analysis

**DOI:** 10.1101/2020.10.28.359828

**Authors:** Serbulent Unsal, Heval Ataş, Muammer Albayrak, Kemal Turhan, Aybar C. Acar, Tunca Doğan

## Abstract

Data-centric approaches have been utilized to develop predictive methods for elucidating uncharacterized aspects of proteins such as their functions, biophysical properties, subcellular locations and interactions. However, studies indicate that the performance of these methods should be further improved to effectively solve complex problems in biomedicine and biotechnology. A data representation method can be defined as an algorithm that calculates numerical feature vectors for samples in a dataset, to be later used in quantitative modelling tasks. Data representation learning methods do this by training and using a model that employs statistical and machine/deep learning algorithms. These novel methods mostly take inspiration from the data-driven language models that have yielded ground-breaking improvements in the field of natural language processing. Lately, these learned data representations have been applied to the field of protein informatics and have displayed highly promising results in terms of extracting complex traits of proteins regarding sequence-structure-function relations. In this study, we conducted a detailed investigation over protein representation learning methods, by first categorizing and explaining each approach, and then conducting benchmark analyses on; *(i)* inferring semantic similarities between proteins, *(ii)* predicting ontology-based protein functions, and *(iii)* classifying drug target protein families. We examine the advantages and disadvantages of each representation approach over the benchmark results. Finally, we discuss current challenges and suggest future directions. We believe the conclusions of this study will help researchers in applying machine/deep learning-based representation techniques on protein data for various types of predictive tasks. Furthermore, we hope it will demonstrate the potential of machine learning-based data representations for protein science and inspire the development of novel methods/tools to be utilized in the fields of biomedicine and biotechnology.

## Introduction

Protein informatics, which can be defined as the use of molecular modelling and data-driven computational methods (e.g., machine learning, statistical modelling) on the proteome to create efficient and scalable solutions, is increasingly becoming an active field of research. The computational methods to come out of such research have the potential to impact daily life through the fields of biomedicine and biotechnology, which have a current market size of 417 Billion USD and is expected to reach 729 Billion USD in 2025^†^. Functional annotation of proteins is critical for protein informatics, as they are the primary inputs to said computational methods. As of October 2020, there are around 189 million protein entries in the UniProt knowledgebase; however, only 0.56 million (around 0.3%) of them are manually reviewed and annotated by expert curators, indicating a large gap between the current sequencing and annotation capacities. This gap is mainly due to the cost and time intensive nature of *in vitro* and *in vivo* experiments and the manual curation of their results. To supplement experimental and curation-based annotation, automated i*n silico* approaches are being used. In this context, many research groups have been working on developing new computational methods to predict proteins’ enzymatic activities^1,2,3^, biophysical properties^4,5,6^, interactions^7^, 3-D structures^8,9,10^, and ultimately, their functions^11,12,13^.

Protein function prediction (PFP) is the assignment of semantic meaning (i.e., functional definitions) to proteins, automatically or semi-automatically. The primary terminology for the functions of biomolecules are codified in the Gene Ontology (GO), a hierarchical network of concepts that annotate molecular functions of genes and proteins, as well as their subcellular localizations and the biological processes in which they are involved^14^. The most comprehensive benchmark project for PFP is the Critical Assessment of Functional Annotation (CAFA) challenge^15^, in which participants predict GO-based functional associations for target proteins. CAFA challenges so far indicate that PFP is still an open problem.

It has been shown in literature that complex computational problems, where features are high dimensional and have complex/non-linear relationships, are amenable to deep learning-based techniques^16^. These techniques can efficiently learn task-related representations from noisy and high dimensional input data. Thus, deep learning has been successfully applied to various domains such as computer vision, natural language processing, and the life sciences^17,18,19,20^.

Deep learning is also a promising avenue of attack for protein informatics. Features of proteins should be extracted and encoded as quantitative/numerical vectors to be used in machine/deep learning-based predictive modelling. A protein representation model, given the raw input features of a protein, calculates a feature vector that is a succinct and an orthogonal representation of the protein. An optimally trained predictive system can efficiently learn features of samples and perform the prediction task using these representations as input. Protein representation construction approaches can be grouped under two categories; *(i)* classical protein representations (i.e., the model-driven approach), which are generated using predefined rules about properties such as the evolutionary relationships between genes/proteins or the physicochemical properties of amino acids, and *(ii)* learned protein representations (i.e., the data-driven approach), which are constructed using statistical and machine learning algorithms (e.g. artificial neural networks) that are optimized on predefined tasks, such as the prediction of the next amino acid on the sequence. Here, the ultimate aim of these representation models is not the prediction of the next amino acid for some sequence, this task is only used in the objective function during the training of the representation model and to measure its success (i.e., its ability to represent the inherent properties of proteins). Later, the output of the trained model, which is the representation feature vector, can be used for other protein informatics-related tasks such as the prediction of function. In this sense, representation learning models leverage the *transfer of knowledge* from one task to another. The generalized form of this process is known as transfer learning^21^ and it is reported to be a highly efficient data-analysis approach in terms of time and cost^22^. Due to this ability, protein representation learning models minimize the need for data labeling^23^.

Protein representation learning methods collect data from one or more resources (e.g., sequences, interactions, etc.) and employ either supervised or unsupervised learning to train a model, which outputs the representation vector to be used in various protein informatics related applications. Supervised and unsupervised training are the two main approaches of system training in artificial learning. Supervised methods require labelled data (e.g., gene/protein entries that are annotated with biomolecular functional definitions such as GO terms), which is mostly produced via experimental procedures and manual curation in protein science. Since the annotation procedure has a high cost, only a small percentage of biomolecular data is labelled. On the other hand, unsupervised models do not need labelling, which makes it easily applicable to any type of biomedical data. However, unsupervised models generally require larger training datasets and additional computational power, especially when deep learning-based methods are used (e.g. GPT-3 which is a state-of-the-art language model trained with 300 billion tokens which costs 3.14E+23 flops^24^). In the framework of data representation learning approaches, unsupervised methods can further be divided into local and global models^25^. Methods in the former group construct representations based on the local context (e.g., in a language model, words surrounding the word of interest in a text), whereas in the latter, the sample is evaluated in terms of a larger, global context (e.g., the whole paragraph or document, the word of interest belongs to).

Protein representation learning method development is a new but highly active area of research, and it mostly gets its technical inspiration from approaches proposed for natural language processing (NLP). It is shown in the literature that various protein representation learning methods, especially the ones that incorporate deep learning, have been successful at extracting relevant inherent features of proteins (Table 1). However, there is no comparative study to systematically evaluate the performance of these methods via quantitative benchmarks, in the context of artificially learning the functional aspects/properties of proteins. Nevertheless, gaining knowledge on complex relationships between sequences, structures, interactions and functions of proteins is especially critical for proposing novel solutions to current biomedical and biotechnological problems.

**Table 1.**
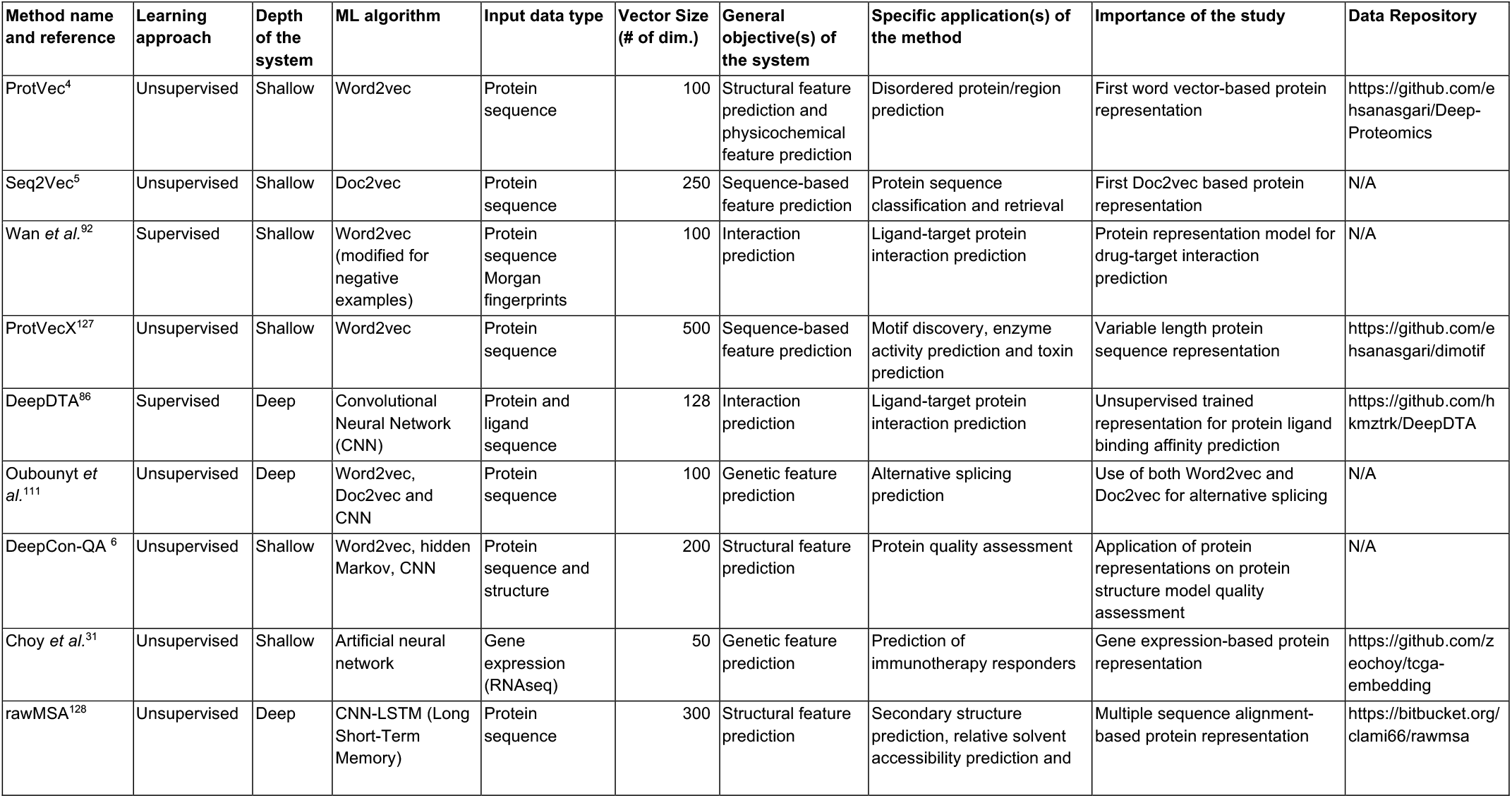
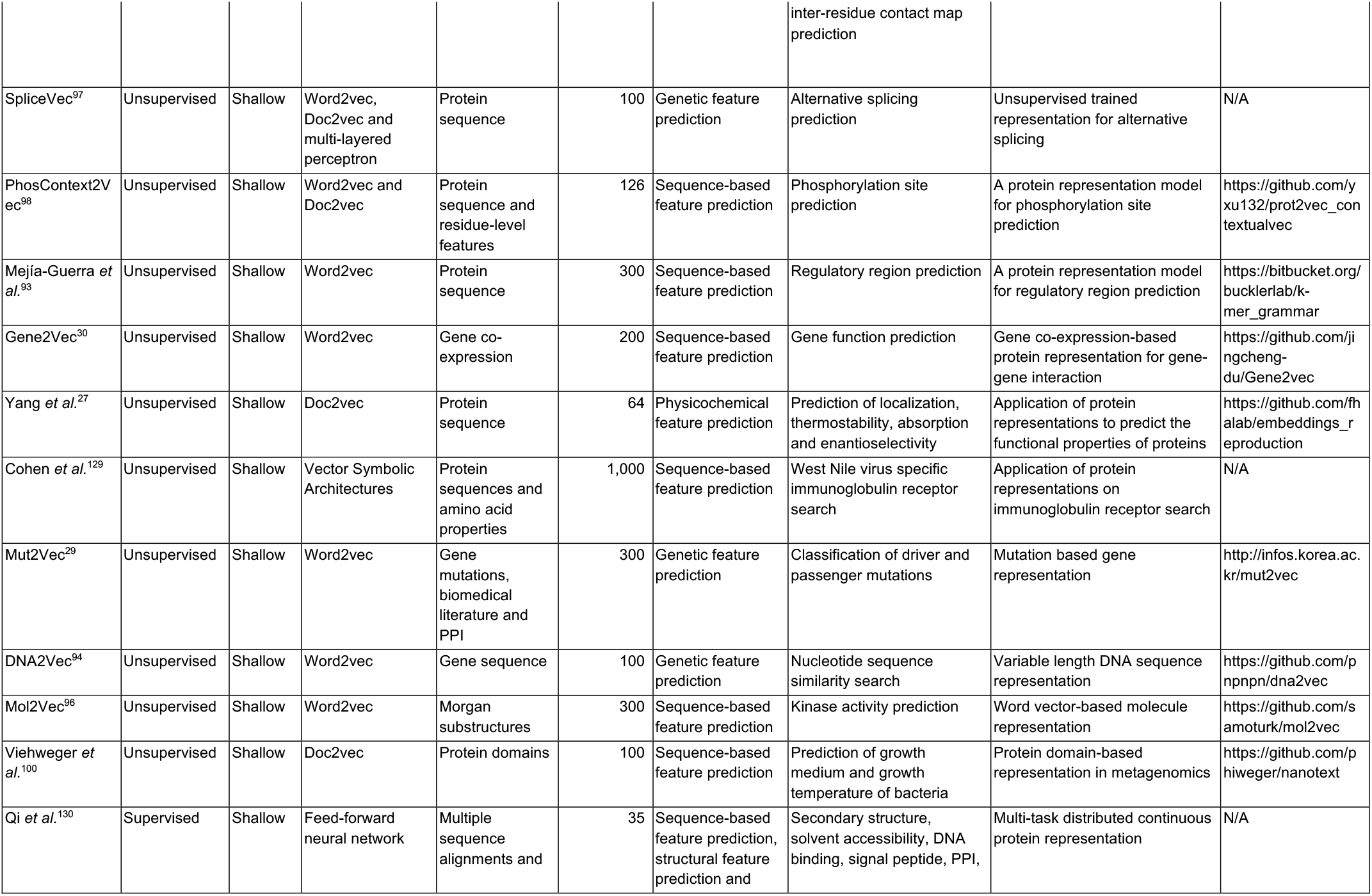
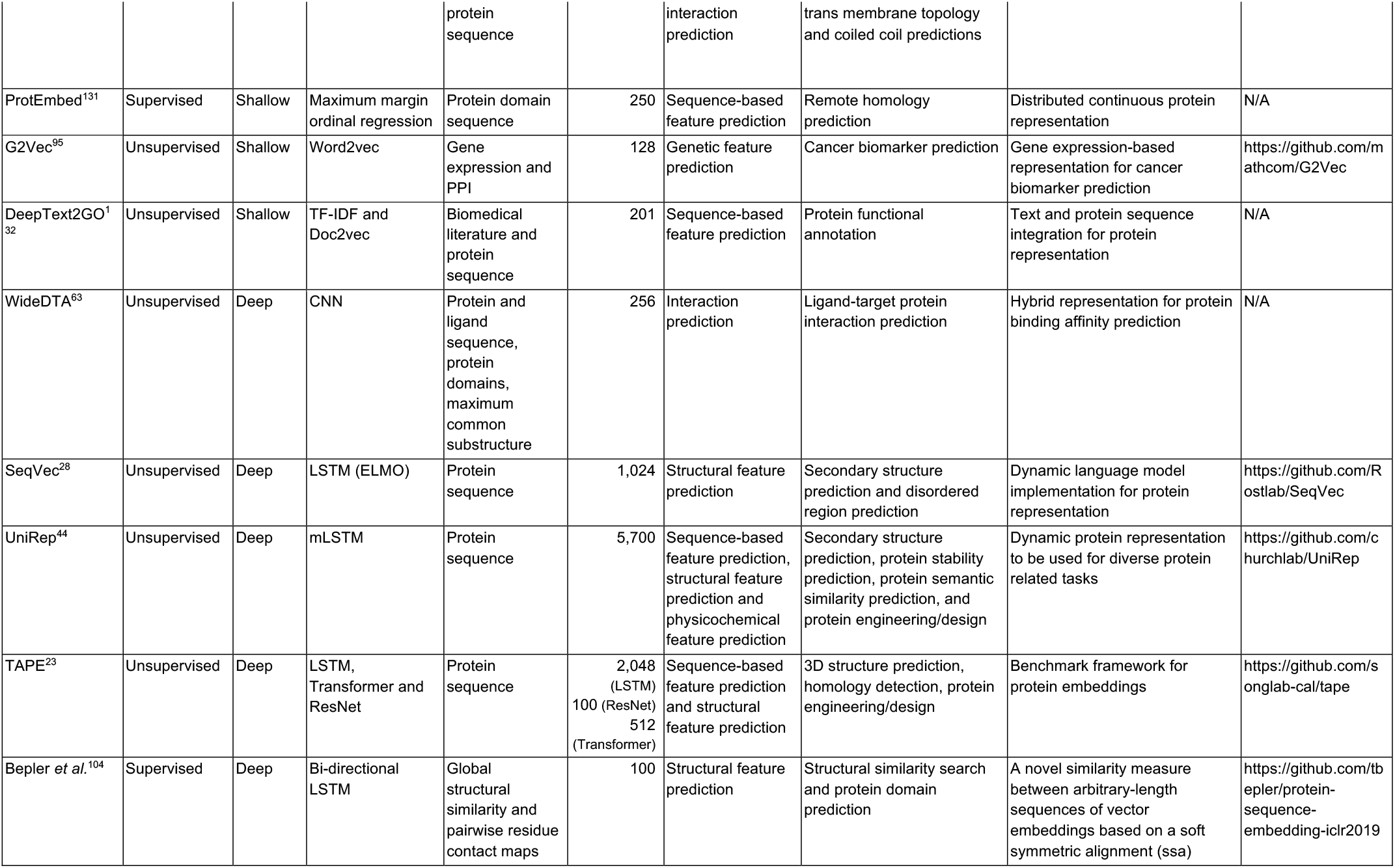
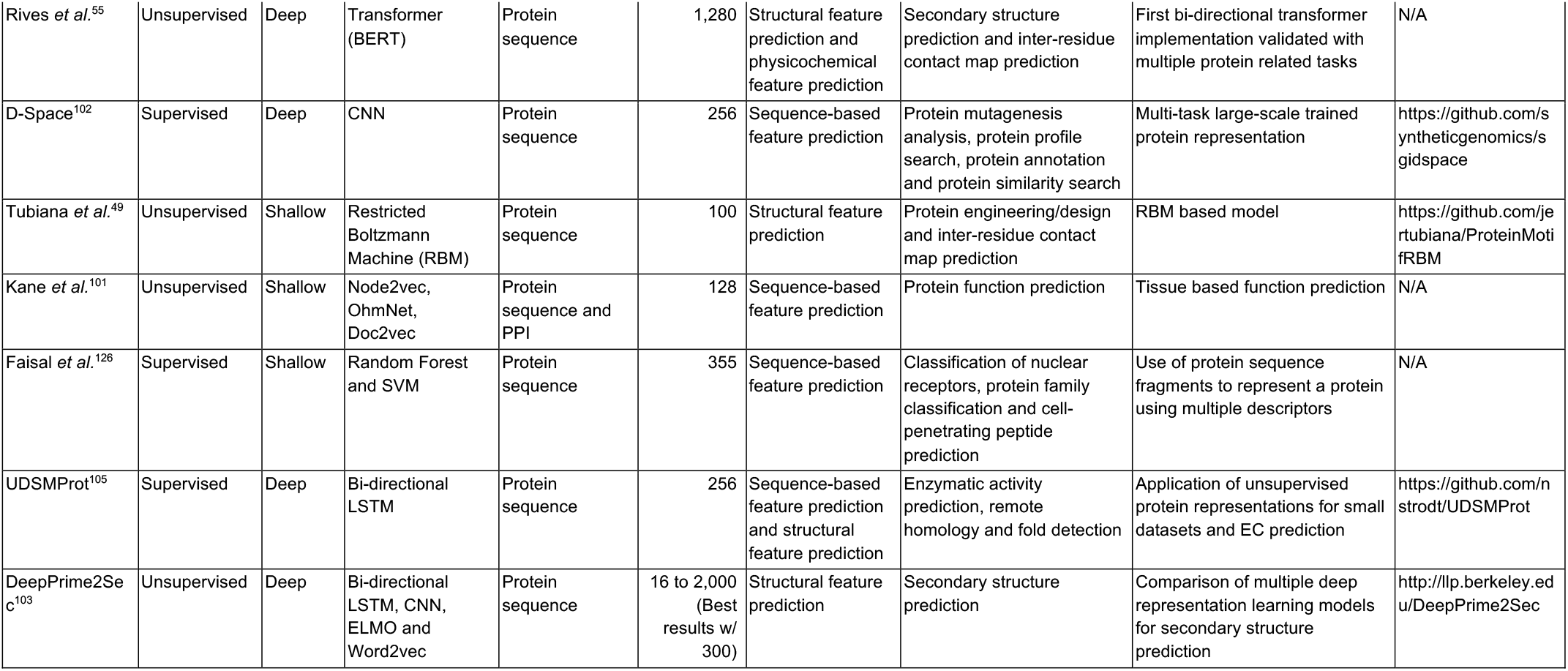
A near-comprehensive list of protein representation learning methods including names, references, learning approach, depth of the system, utilized machine learning algorithm, input data type, vectors sizes, objectives and applications of systems, the importance of studies, and the availability as tools or source code (vector sizes vary for some of the methods, in those cases, we indicate the vector sizes that yield the best predictive performance).

In this study, we conduct an investigation of the available protein representation learning methods that were proposed since 2015, with a detailed benchmark analysis regarding the potential of these methods to capture the functional properties of proteins. We explain both classical and learning-based methods to provide insight into their respective approaches to represent proteins, and we classify these methods according to their technical aspects and objectives (please see Methods section and the supplementary information document). Aiming to evaluate how much each representation model captures different facets of functional information, we constructed and applied benchmarks based on; *(i)* semantic similarity inference between proteins, *(ii)* ontology-based protein function prediction, and *(iii)* drug-target protein family classification (Results section). Finally, we discuss the results and current issues, and provide a perspective on the future of learned protein representations (Discussion section). The whole study is summarized in Fig. 1a. We expect that the discussion and conclusions of this study will inform researchers who would like to apply machine/deep learning-based representation techniques on biomolecular data for predictive modelling. We believe our investigation will be valuable in gaining insight about the potential of machine learning-based protein representation models in retrieving complex functional relationships of proteins, since previous studies mostly evaluated a few methods over tasks related to the structural features^23,26^. Finally, we hope this study will inspire new ideas for the development of novel, sophisticated and robust approaches to solve open problems in protein informatics.

**Figure 1.**
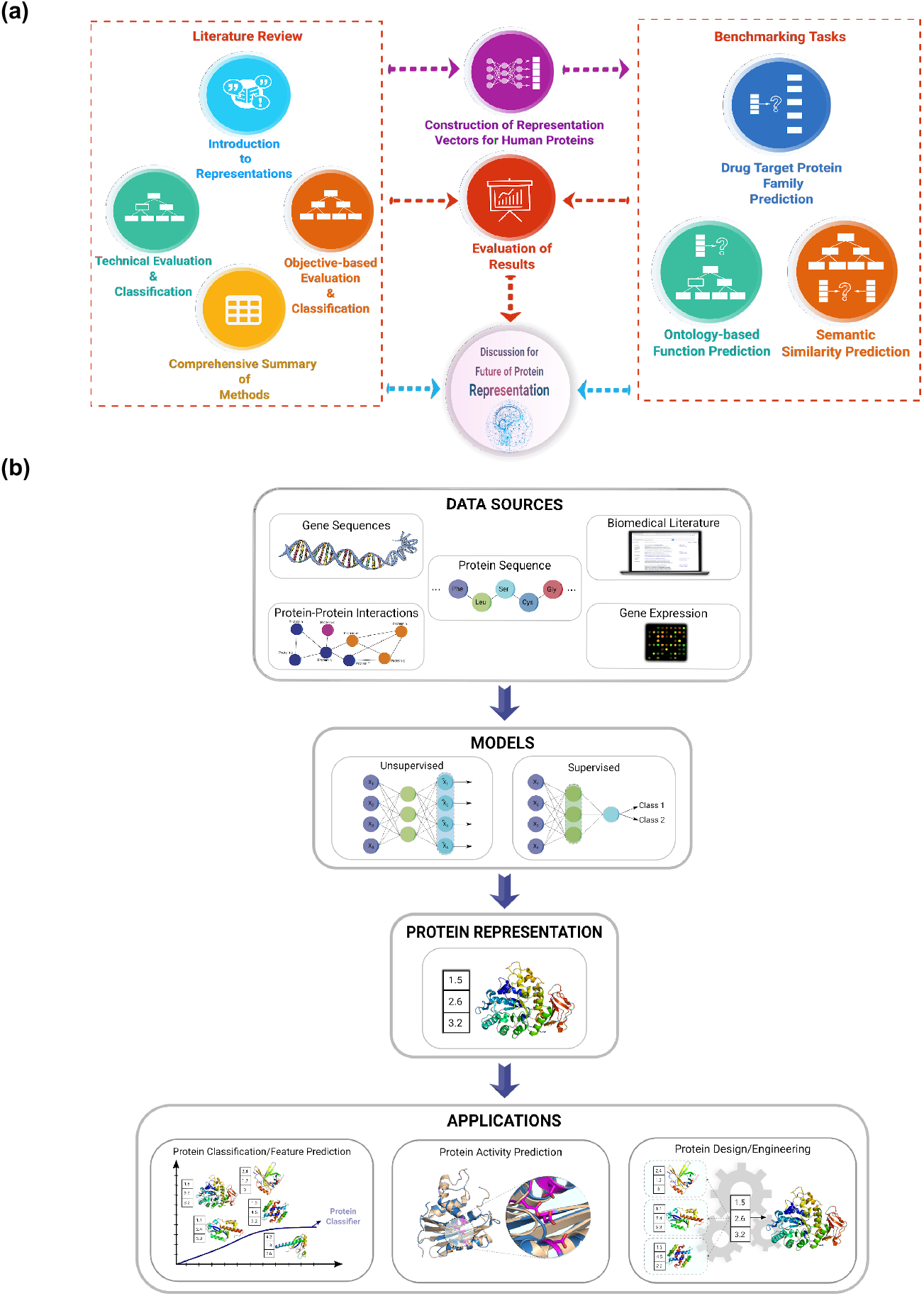
**(a)** Overview of the protein representation benchmark study; **(b)** various data sources/types can be utilized to construct representations, and this data can be used to train unsupervised or supervised models, and the output representation vectors can be used for diverse applications.

## Results

In this section, we focus on protein representation benchmark analyses. The review of the literature, including the construction and application of protein representations (Fig. 1b), and their technical and objective-based classification and evaluation (Fig. 2) are given in the Methods section and in the supplementary information document.

**Figure 2.**
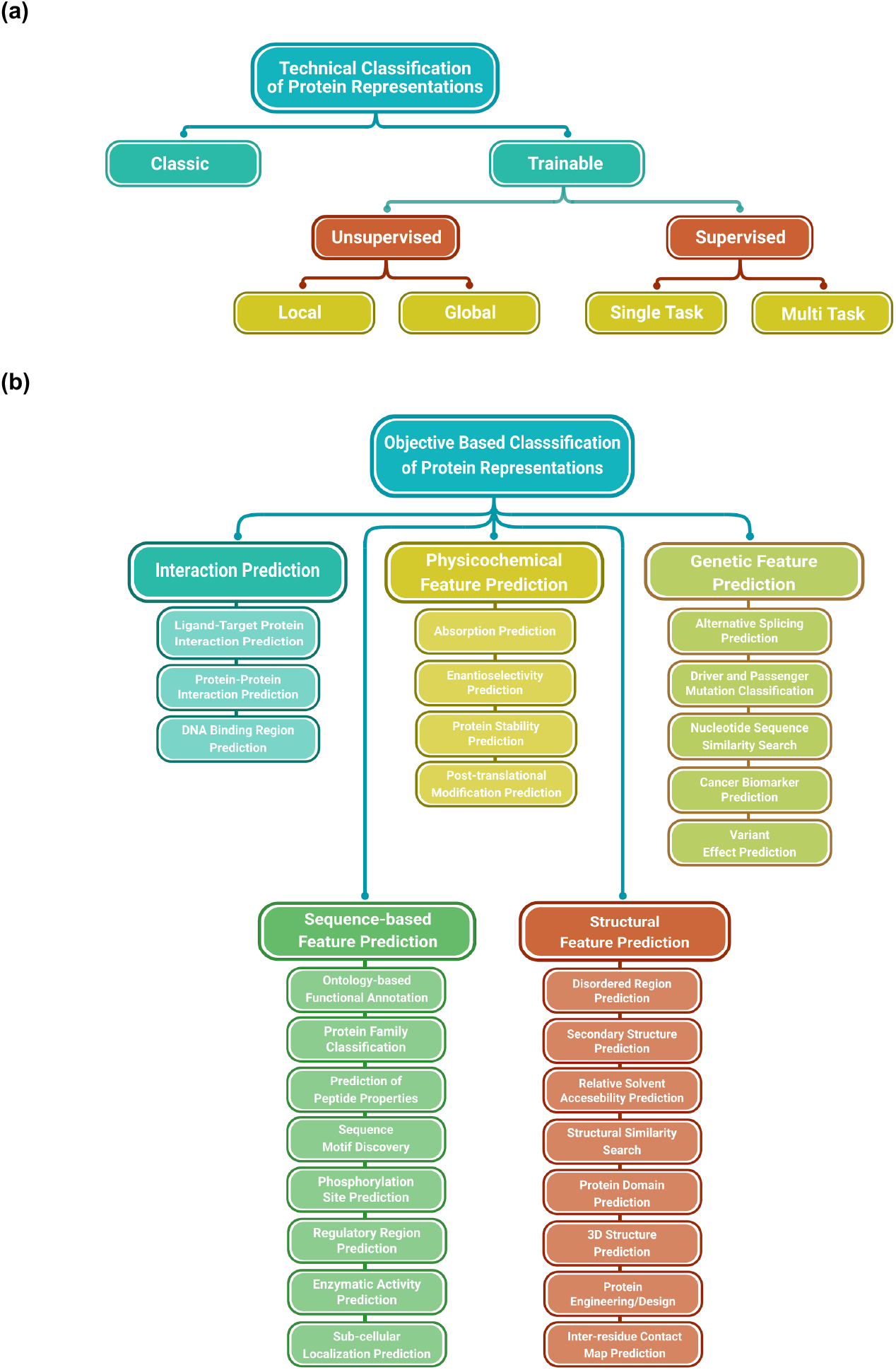
**(a)** The classification of the technical approaches used in protein representation learning, and **(b)** the classification of the objectives of protein representation methods.

Nine different representation learning methods have been selected for our functional property-based benchmarks, according to their previously reported success in predictive tasks, and subject to their availability as open access tools or ready to use pre-constructed feature vectors. During the selection process, we also considered the source protein features/attributes used to train these methods (e.g., sequence, PPIs, etc.) and the algorithmic approaches, with the aim of covering a wide variety of methodologies. The methods included in the benchmark are thus; LearnedEmbeddingVec^27^, SeqVec^28^, Mut2Vec^29^, Gene2Vec^30^, TCGA_Embedding^31^, ProtVec^4^, TAPE-BERT_Avg^23^, TAPE-BERT_Pool ^23^, UniRep^23^, along with two classical representations: APAAC ^32^ and k-sep-bigrams^33^, as baselines. Technical information about these tools is given in the Methods section. A near-comprehensive summary of 35 protein representation learning methods obtained from the literature, including the above-mentioned benchmark methods, is given in Table 1.

### Semantic Similarity Inference

In this analysis, we aim to measure how much information about biomolecular functional similarity is captured by the representation models. In this context, Gene ontology (GO) annotations are utilized, which signify the molecular functions, large-scale biological roles, and subcellular localizations of proteins. We first calculated vector similarities, which are defined in terms of pairwise quantitative similarities (e.g., cosine, Manhattan and Euclidean) between representation vectors of proteins in our dataset. These similarities were then compared to the ground truth functional similarities, which are measured based on the actual GO annotations of these proteins using standard semantic similarity measures (e.g., Lin similarity^34^). To be able to compare the success of different protein representation methods, we calculated Spearman rank-order correlation values between representation vector similarities and the actual GO-based semantic similarities of the same protein pairs, using 4 different test datasets (explained in the Methods section). Higher correlation values indicate higher success. The results based on cosine similarity are shown in Fig. 3. Performance results considering the Manhattan similarity and Euclidean distance measures can be found in Fig. S3 and S4.

**Figure 3.**
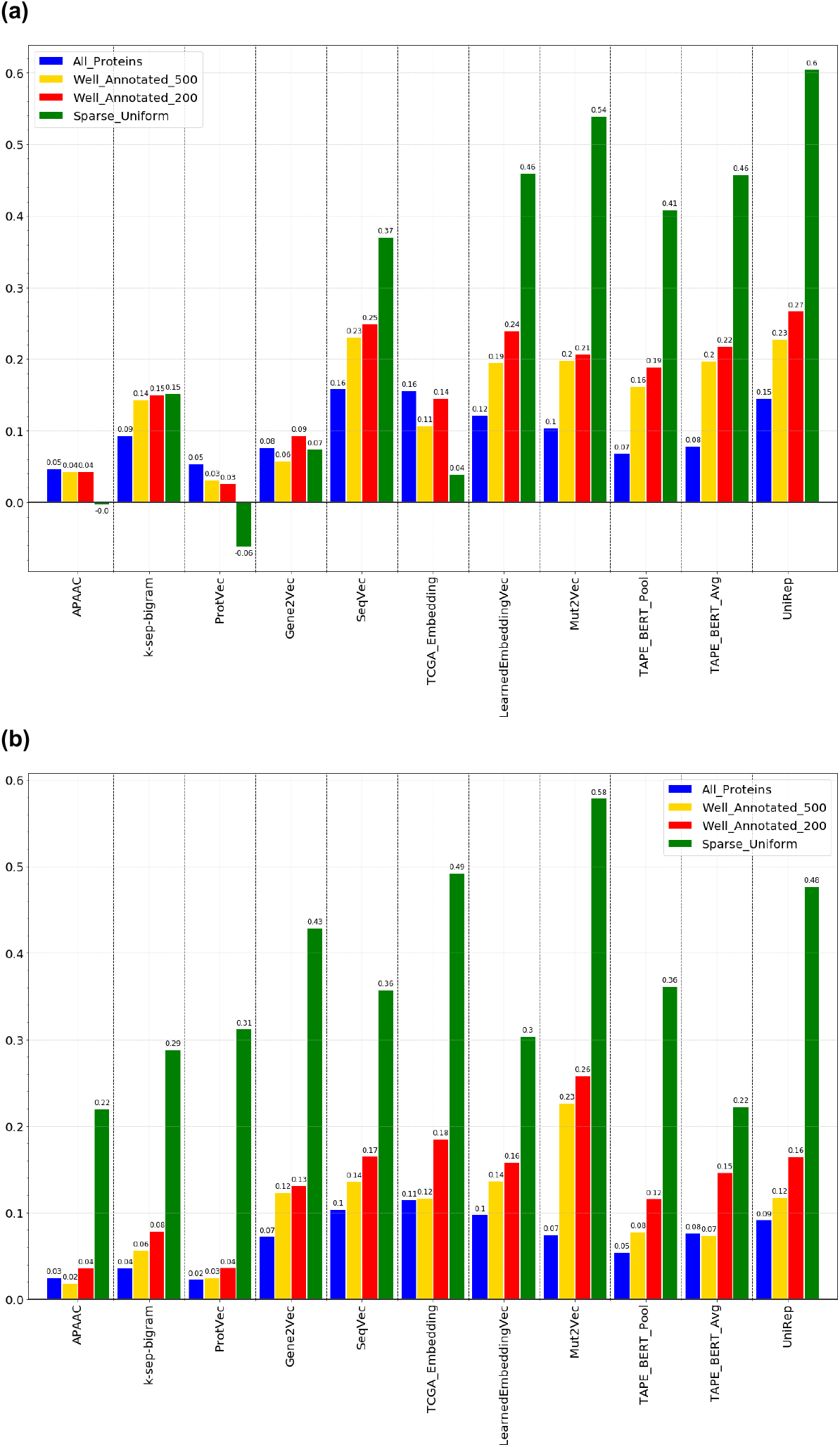
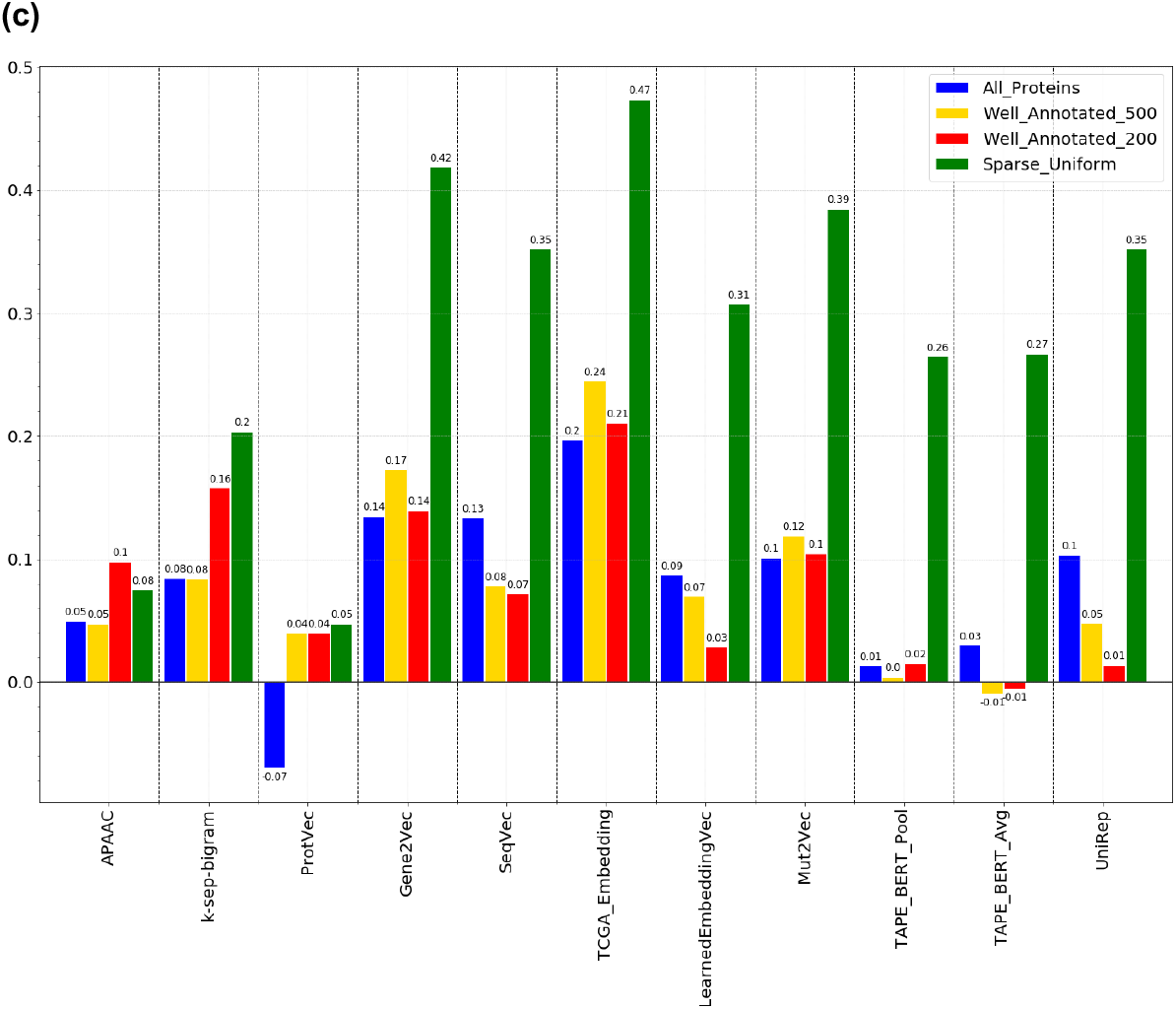
Performance of protein representation learning methods in inferring pairwise semantic similarities between proteins, calculated in terms of Spearman correlation between the ranked true pairwise similarity list (calculated using Lin similarities^34^ between functional annotations of proteins) and the representation-based ranked pairwise similarity list (calculated using cosine similarities between numerical feature vectors of proteins). True semantic similarities are calculated based on GO terms of; **(a)** molecular function, **(b)** biological process, and **(c)** cellular component categories.

According to the results presented in Fig. 3a, UniRep is the most successful representation model in the GO molecular function (MF) category, considering all four datasets. Mut2Vec^29^ is the best performer in the GO biological process (BP) except for the “all proteins” dataset, and TCGA_Embedding achieved the highest correlation score in the GO cellular component (CC) category. Mut2Vec is also among the top three methods in MF and CC categories. TCGA_Embedding is also in the top three in the GO BP category. BERT, Gene2Vec, and LearnedEmbeddingVec are other notable methods but did not achieve the top performance in any of the categories.

UniRep^23^ is based on multiplicative LSTM^35^. We propose two reasons to explain the higher performance of UniRep. The first reason can be that, UniRep constructs and uses a training sequence dataset with low bias (consisting of 24 million UniRef50^36^ protein entries, which are filtered by a 50% similarity threshold from the from UniProtKB, instead of using all available protein sequences in the data source), which might be providing better generalization capability. The second reason may be the size and the information content of the model. In our benchmarks, we employed the “UniRep Fusion’’ model since this version had the highest performance according to the original UniRep study. This model was built with the concatenation of the “final hidden state”, “final cell state”, and “average hidden state” of the LSTM model, each of which has a size of 1×1900, providing a total vector size of 5700. In our opinion, the concatenation of the different states might have enhanced the protein representation vector with different levels of semantic information, since the level and type of information learnt at each layer is claimed to be distinct^16^.

Mut2Vec^29^ was originally developed to predict the effects of mutations, but surprisingly it performed very well in our analysis considering the BP based semantic similarities. The model was developed using patient mutation profiles, biomedical literature and protein-protein interactions. The last two datasets may include information considering the role of the proteins in BPs. For example, considering that two proteins are interacting, then observing them as a part of the same BP is highly probable. Similarly, supposing two proteins had a role in the same BP, they may frequently be observed together in the same text (e.g., article). As a result, those proteins would probably be embedded proximally in the vectorial semantic space. We are suggesting that the top performance of Mut2Vec probably depends on these factors.

TCGA_Embedding^31^ exploited gene expression data and a simple learning system inspired by non-negative matrix factorization. With this approach, the authors constructed a representation with a vector size of 50, which is one of the smallest representations in our benchmark. TCGA_Embedding scored the best performance in the prediction of CC-based semantic similarities, together with a notable performance considering the BP-based similarities. Similar to TCGA_Embedding, the Gene2Vec^30^ model utilizes gene co-expression data with the skip-gram algorithm. The model performed well in both BP and CC based semantic similarity inference tasks. Gene (co)expression profiles are one of the least studied data types for developing protein representations as only a few studies exist; however, it is found to be quite informative to infer the similarities between proteins in terms of the BPs they take part in and the CCs that they localize to. It was reported in the literature that there is a correlation between the expression profiles and the subcellular locations of genes/proteins, which was evaluated in the context of machine learning-based prediction of protein localizations^37,38^. It was also discussed in the results of the CAFA Pi challenge (over the bacterial motility and biofilm formation biological processes) that gene expression is a critical input data type for predicting the biological roles of proteins^15^.

TAPE-BERT^23^ had the second place in the MF-based similarity inference task. TAPE-BERT (a bidirectional transformer) is the only method that uses the self-attention mechanism in this analysis. Technical details of BERT and the self-attention are discussed elsewhere^39,40^. Similar to the success of the BERT model in word sequences (sentences)^41^, TAPE-BERT model can represent protein sequences with high accuracy. It should also be noted that the implementation we used here was directly obtained from the TAPE benchmark study^23^ without any fine-tuning. TAPE-BERT may perform better in protein informatics tasks with further optimization.

Finally, LearnedEmbeddingVec^27^, which is a simple model that takes protein sequences at the input level to process them using doc2vec^42^, scored as good as the TAPE-BERT^23^ models in our semantic similarity based analysis. The size of the TAPE-BERT models were notably larger compared to LearnedEmbeddingVec^27^ (i.e., 12 hidden layers with 768 neurons for each layer, as opposed to 1 hidden layer with 64 neurons). We argue that some of the shallow models still preserve their significance, especially in MF-based semantic similarity inference.

### Ontology-based Protein Function Prediction

As the second benchmark of our study, we aimed to assess the success of protein representation models in terms of automated protein function prediction (PFP). In this analysis, Gene Ontology^14^ (GO) term annotations of proteins were used to train and test the same 11 protein representation models via supervised machine learning based classification. In this benchmark, we preferred to use a linear classifier (i.e., linear support vector classification from scikit-learn^43^) in order to prevent non-linear transformations on the protein representation vectors, since the relevant information hidden in proteins should have been captured and extracted by the representation model beforehand, if the model is successful. This way, we could evaluate the protein representation models in terms of their success in extracting this information without additional factors.

Here, we also discussed a critical topic that was mostly overlooked in previous PFP studies, the assessment of the performance in terms of annotated GO term specificity. This is important since there is a relation between the specificity of a GO term (i.e., its location of the graph of GO) and its informativeness. For example, considering an annotation with the GO term “negative regulation of molecular function” the provided information is too general. These GO terms are generally located near the root of the GO graph and called “shallow terms”. If the same protein was annotated with the GO term “negative regulation of double-stranded telomeric DNA binding”, which is a descendent term of the previous one, the annotation would have been more informative. In order to take this phenomenon into account in our analysis, we simply grouped GO terms under three categories as shallow, normal, and specific; according to their level on the directed acyclic graph of GO (see Methods).

One key problem in applying deep learning to protein informatics is the requirement for high amounts of training data^11^. To examine this issue in our benchmark, we considered a grouping among GO terms regarding the number of proteins they are annotated to. This approach is expected to uncover the performance of representation learning models in terms of learning with only a few training examples, which also is the case for a considerable number of informative GO terms. Furthermore, some of the protein functions are studied well and some others are understudied, which creates a discrepancy in terms of the number of annotated proteins. We expect that this approach will be useful to assess the representations considering their ability to learn under-studied functional properties. For this, we created three categories that point out the number of the proteins annotated to a GO term as; low, middle, and high (see Methods).

PFP performance results are given for 9 different GO groups using F1-score based heat maps in Fig. 4. The overall GO term prediction performance results (averaged over 9 different GO groups) in terms of recall, precision, F1-score, accuracy, and Hamming distance are given in Table S4. It is important to mention that these performances are higher compared to the results of CAFA challenges, due to the way we modelled the experiment. We only run a test sample on the model that contains its true label as one of the 5 tasks (i.e., GO terms), instead of running all test samples on all prediction models. The reason behind this experimental design choice was to prevent the accumulation of the scores of all benchmarked methods in low performance regions (especially for hard-to-predict ontologies such as BP), which would prevent the observation of the performance differences in-between.

**Figure 4.**
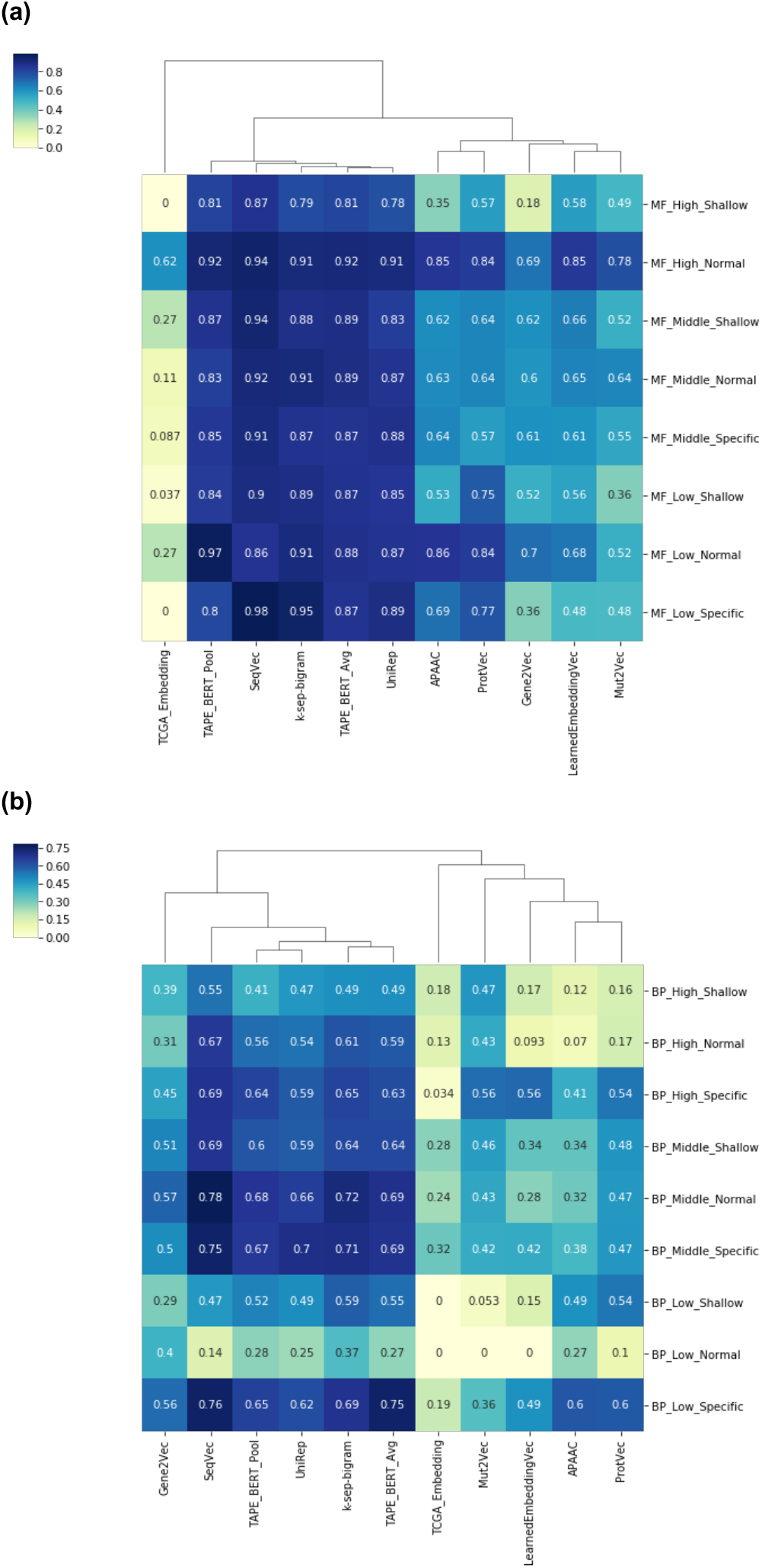
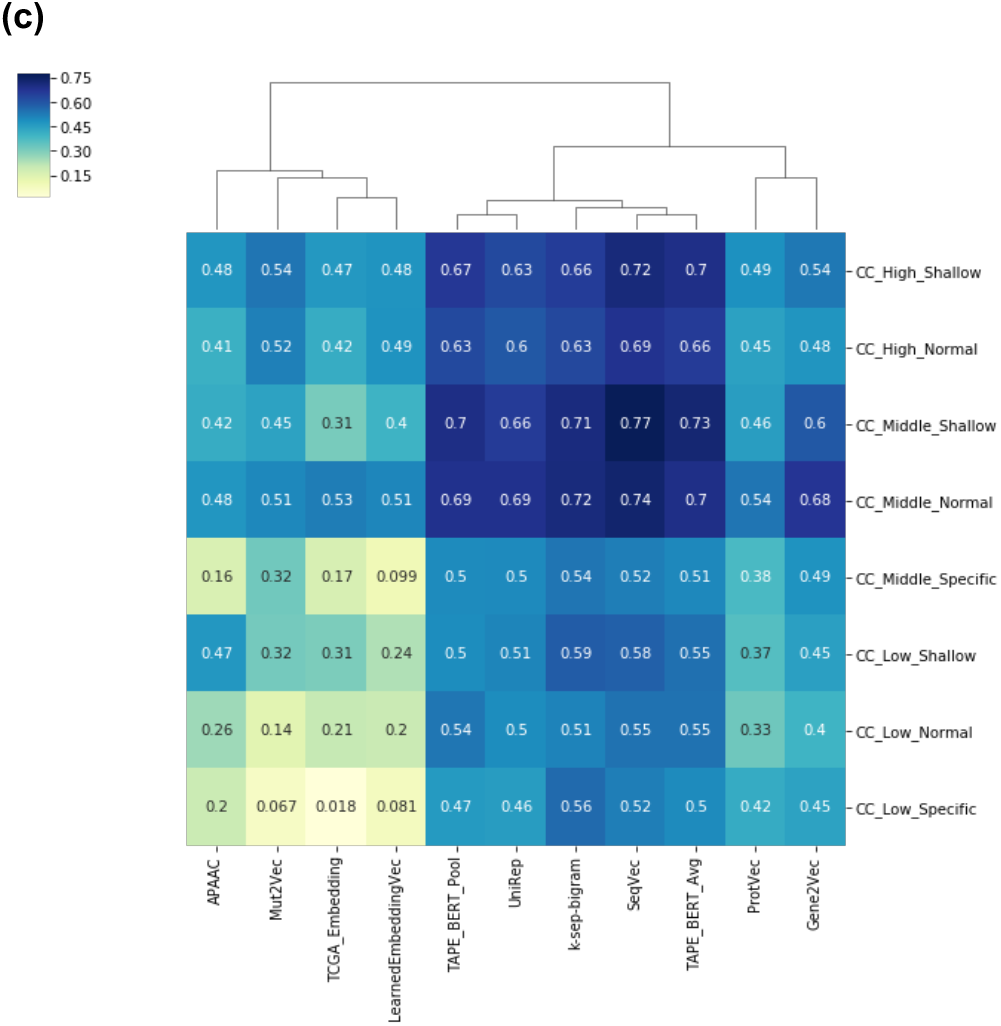
Heat maps indicating the performance results (weighted F1-scores) of protein representation learning methods in ontology-based protein function prediction benchmark in terms of GO; **(a)** molecular function annotations, **(b)** biological process annotations, and **(c)** cellular component annotations.

It is shown in both Fig. 4 and Table S4 that, in the MF prediction task, top methods showed similar performances across almost all GO groups (e.g., low, high, specific, shallow, etc.), among which SeqVec^28^ got the top place, and k-sep-bigrams^33^, TAPE-BERT^23^ models and UniRep^44^ came right after. For BP and CC prediction tasks, SeqVec was the best performer together with k-sep-bigrams, and runner ups were again TAPE-BERT models and UniRep. We also observed that these methods are clustered together in all three heat maps, considering their performances (Fig. 4). These four representation learning methods share common characteristics that can explain their similar performance in the PFP benchmark. First of all, they are all based on large state-of-the-art sequence modelling algorithms: LSTM (SeqVec and UniRep with 93M and 18.2M parameters, respectively) and a Transformer (TAPE-BERT with 110M parameters). Also, they share the same model training objective, prediction of the next amino acid on the sequence. Finally, they were all trained with large datasets (24M sequences for UniRep, 33M for SeqVec and 31M domain sequences for TAPE-BERT).

We also calculated model performances averaged for each GO group (Table S5), especially to observe the scores for challenging groups; the “low” group which contain GO terms that have low effective number of proteins (during dataset preparation, we eliminated highly similar proteins by filtering through UniRef clusters, to be able to take the effective number of samples into account while grouping GO terms), and the “specific” group consisting of GO terms that are leaf nodes, or are close to the leaf nodes, in the GO hierarchy. In Table S5, mean F1-score results (considering the average of all methods) indicate that low number of samples is a problem for BP and CC categories, but not so much for MF category, where there is generally an explicit relation between the input (i.e., sequence) and the label. On the other hand, we could not observe a trend in performance change in terms of GO term specificity. As a result, it can be stated that prediction success for informative specific terms may be solely related to the effective number of training proteins. For the MF-low category, k-sep-bigrams (F1:0.916) and SeqVec (F1:0.914) achieved the best performances. In the BP-low and CC-low categories, k-sep-bigrams had the best F1-scores with 0.548 and 0.556, respectively. For the MF-specific and BP-specific categories, the SeqVec model got the top scores with F1:0.945 and F1:0.732, respectively. Considering the CC-specific category, the k-sep-bigrams model again got first place with F1:0.552. These results showed that, for the tasks where the number of labelled data points are low, the representation capability of classical (model-driven) methods is still higher compared to learning-based (data-driven) models.

Overall, the best performing method in the protein function prediction benchmark, considering all GO categories, was SeqVec^28^, whereas, k-sep-bigrams^33^, a classical protein representation based on evolutionary relationships, got second place with scores close to the top performer.

### Drug Target Protein Family Classification

In our third benchmark analysis, we aimed to measure the performance of protein representations in the framework of drug discovery, with the prediction of drug target proteins’ main families (i.e., enzymes, membrane receptors, transcription factors, ion channels and others). Since these families are made up of proteins with distinct structural characteristics, this benchmark analysis will also reflect the ability of these representations in learning structural properties. Furthermore, by using a data source other than GO annotations, we seek to diversify our benchmark and to evaluate the representations from a different perspective. Similar to the ontology-based protein function prediction benchmark, here we preferred to use a multi-task linear SVM classifier in order to solely measure the ability of learned protein representations in extracting the complex protein attributes/properties.

The number of samples in this analysis was lower compared to the previous benchmarks (Table S6), as we only used human target proteins listed in the ChEMBL database^45^. Since there is an imbalance in terms of the number samples for each task (i.e., protein family), MCC was taken as the most reliable indicator in comparing the representation methods. According to the average cross-validation results of our multi-task classification model (Fig. 5), SeqVec^28^ model is the best method in all metrics, whereas TAPE-BERT_Avg^23^ came a close second. UniRep^44^, TAPE-BERT_Pool^23^, and k-sep-bigrams^33^ models also provided remarkable performances. Family specific prediction scores (Fig. S5) showed that SeqVec provided the best performance in terms of enzymes. For membrane receptors, UniRep provided the best performance, followed by SeqVec. For transcription factors, TAPE-BERT_Avg took the top place. For ion channels and others, SeqVec was again the best performer.

**Figure 5.**
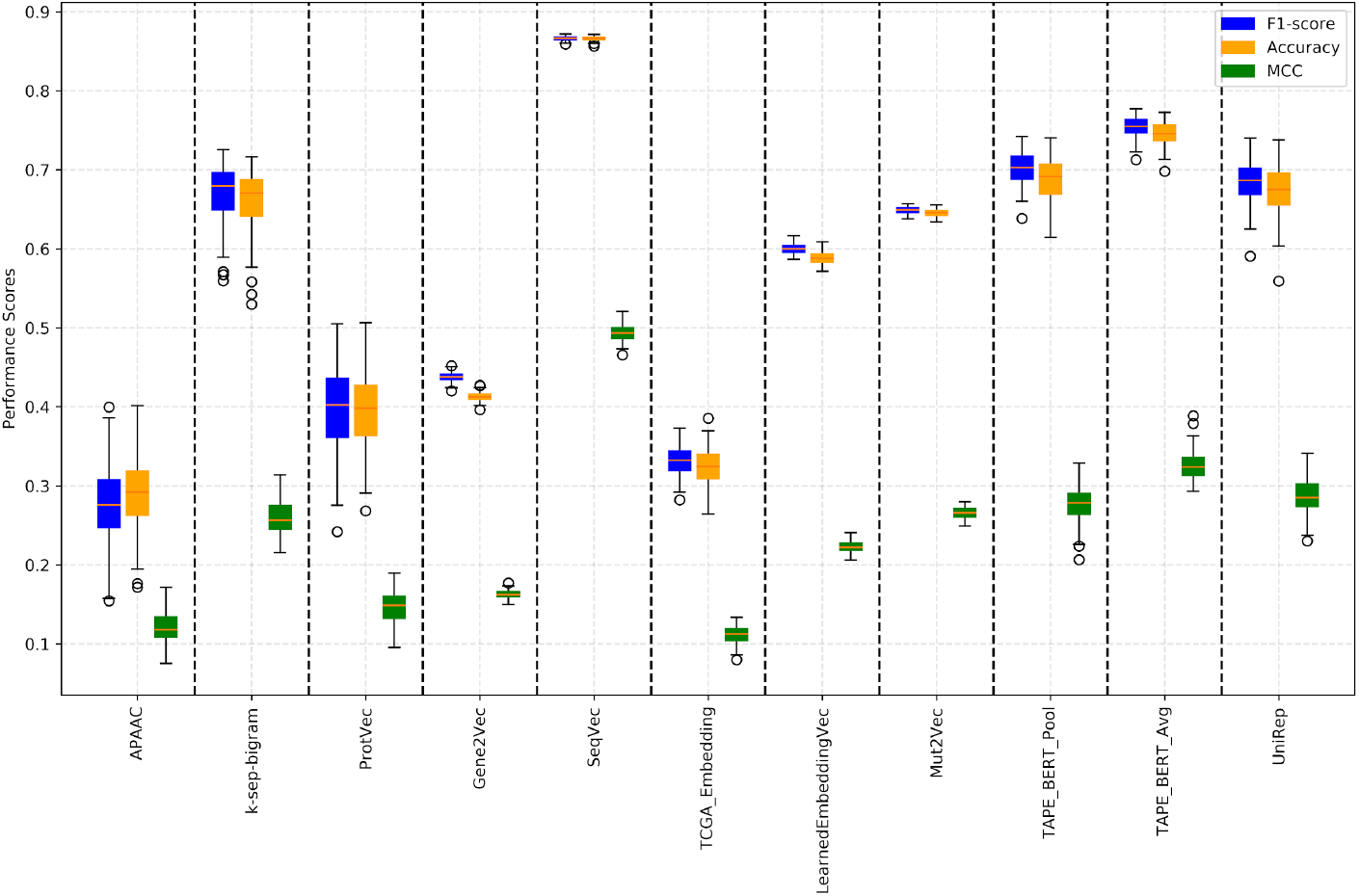
Box plots indicating performance results (iF1-score, accuracy and MCC) of protein representation learning methods in the drug target protein family classification benchmark.

Given that SeqVec was also very successful in the protein function prediction benchmark, especially in MF prediction, and that protein family information is related to functions, there is a plausible correlation between these results. SeqVec uses the ELMO model^46^, a bi-directional LSTM with 93M parameters capable of learning long sequential patterns, which is stated to be highly efficient for language modeling^41^. The most evident difference of SeqVec from the other successful state of the art models in our study (e.g., UniRep and TAPE-BERT) in terms of the model architecture is that SeqVec contains a CNN layer to embed the amino acids in the sequence onto a latent space, before the LSTM layers. In the original ELMO model, the same approach, charCNN^47^, was mainly used to obtain word vectors of fixed size. It is also important to mention that SeqVec displayed a moderate performance on the semantic similarity inference benchmark. This observation indicates that, although protein function prediction and semantic similarity inference can be seen as correlated tasks due to sharing the same information source, specialized solutions are required for each one. The moderate performance of SeqVec on semantic similarity inference might be explained by the noise on the original representation vectors that SeqVec produced. This noise may be filtered out by a simple feature selection during our supervised training in the PFP and drug target protein family classification benchmarks, as a result, SeqVec was successful. However, there was no supervised training in semantic similarity inference. This phenomenon was also observed in the original SeqVec study (see Fig. 2 of the SeqVec paper^28^). In the first t-SNE plot of this figure, protein classes are distributed heterogeneously when the unsupervised model was directly used to generate protein representation vectors. Contrarily, when the representation vectors generated by the supervised model were used, the classes were successfully clustered.

## Discussion

The volume of AI-based protein informatics studies has been growing lately to further the understanding of the complex relations between sequence, structure and function^48^. In this study, we evaluated protein representation learning methods in terms of their ability to capture functional properties of proteins, to be utilized, ultimately, to overcome critical challenges in protein informatics, biotechnology, and biomedicine domains. These models, with their modest resource requirements and high representation performance, can be (re)used for a variety of tasks. Thus, we argue that learned representations will play an essential role in protein research and development in the near future.

In the PFP benchmark, performances observed in CC and BP GO term prediction tasks were lower compared to the MF prediction. This observation is plausible since most of the learningbased methods use protein sequence data as input, and sequence is not a direct marker for localization (without the cleaved signal peptides) or the biological role of proteins. Also, we observed that the success rate in CC term prediction decreases with the decreasing number of annotated proteins. A similar observation was valid for MF and BP categories as well; however, the effect was less pronounced. On the other hand, we did not observe a similar trend in performance change with increasing or decreasing term specificities. Nevertheless, it is possible to state that the issue at hand is still critical since many of the informative “specific” GO terms have “low” number of annotated proteins.

In semantic similarity inference and drug target protein family prediction benchmarks, we observed that some of the learned representations are superior to the classical ones in terms of predictive performance, thus justifying the benefit of the data-driven approach to represent the functional properties of biomolecules. On the other hand, k-sep-bigrams, a classical protein representation method that does not need any training, could compete with deep learning-based protein representation methods in the PFP benchmark. These results indicate that evolutionary relationships are correlated with functional properties of biomolecules to such a degree that a simple representation that utilizes this feature can perform as good as the most complex sequence modelling methods. In the light of these results, we claim that the explicit incorporation of evolutionary information into the training of representation learning models would lead to significant improvements considering predictive performances in protein informatics. Nevertheless, we still claim that learned protein representations, in their current state, are essential for different reasons. First of all, learning-based models produce reusable vectors, which can be optimized towards increasing predictive performance in challenging tasks (e.g., BP GO term or 3-D structure prediction) with further training on selected supervised tasks. Second, studies indicate that protein representation learning models can also be employed for designing new proteins using the learned probability distributions of the proteins in the training set^49,50^. This is not possible with classical representations. The topic of protein design is discussed further, below. Third, the results of our benchmarks indicate that the approach of learning protein representations is an excellent application field for transfer learning, as it is possible to train a model that could generate a representation vector, and use this vector for various related predictive tasks (e.g., training the model via the prediction of the next amino acid in the sequence and later using the trained model for predicting functions, interactions, or structures).

In benchmark studies, the possibility of data leak from training to test is a critical issue that should be considered during performance testing. Data leak can be defined as the accidental share of knowledge between train and test, leading to overoptimistic performance measurements. In our analyses, we observed that certain representation models performed well in tasks that are biologically related to the tasks that these models were trained on; although the data and the actual tasks were different from each other. For example, Gene2Vec utilized a hyperparameter optimization task, which aims to maximize the clustering of genes within MSigDB^51^ functional pathways. This task is assumed to have latent knowledge about BP and CC based protein semantic similarity inference and ontology-based PFP benchmarks, where Gene2Vec showed a notable performance. Likewise, Mut2Vec^29^ uses protein-protein interaction data for training, and we found that this model is successful in predicting BP and CC related tasks. It is highly probable that two interacting proteins are localized to the same cellular compartment or have a role in the same biological process. Finally, TAPE-BERT, one of the best performers especially in the MF-based PFP and drug-target protein family classification, is trained on protein domain sequences provided by the Pfam database. The selection of the sequence fragments based on protein families can be stipulated to have led to knowledge transfer from Pfam to the protein representation vectors calculated by TAPE-BERT. In our opinion, these cases of knowledge transfer are unlikely to be counted as examples of data leak from training to test, since the data and the tasks used in train and test were completely independent. Hence, these protein representation models should be considered successful in terms of inferring relevant information from the input data, in the scope of this study. Nonetheless, particulars of such knowledge transfer is an interesting topic to be further investigated in future studies.

There are several challenges within the field of protein representations. First of them is related to the assessment of newly proposed methods, as the proper evaluation of stability and robustness of models is critical. So far in the literature, protein representation models are tested only with small-scale datasets and limited tasks. On the other hand, there are studies (unrelated to protein informatics) in which authors proposed new approaches for rigorously evaluating the properties of data representation models^52-54^. These studies can be exploited and adapted for the evaluation of learned protein representations. Another key challenge is associated with model sizes. In the NLP domain, the number of parameters is steadily increasing with every new high-performance model (e.g., state-of-the-art GPT-3 model has 175 billion parameters). As most of the successful protein representation learning approaches are based on NLP models, this trend is also observed in the protein representation field^55^. This may pose a critical problem in terms of increasing the computational cost to extreme scales, especially for embedding large samples^56^ using sequencebased protein representations, where each amino acid in a sequence is modelled as a word in a sentence^26^. To make a simple comparison, the average size of a sentence in English is 21.7 words^57^; however, the median number of amino acids in human proteins is 361^58^, which makes the problem even more pronounced for protein informatics. There are potential solutions for this issue in the literature^56,59–61^, mainly proposed for NLP-related tasks. These solutions may also be exploited for protein sequence representations. It should also be noted that model sizes (e.g., number of hidden layers, total number of parameters) are not necessarily correlated with performance in protein representation models^26^. We observed this phenomenon in our benchmarks as well. For example, the UniRep model has 18.2M parameters but could compete with much larger models such as SeqVec (93M parameters) and TAPE-BERT (110M parameters). Therefore, constructing larger and more complex models may not always be the solution for better representations. Instead, investing time and resources on the incorporation of diverse types of biological data into the models would be a better choice.

Model interpretability is a critical topic to understand why a model behaves the way it does. In an interpretable (i.e., explainable) representation, all features are encoded in a distributed form, which means that the feature(s) corresponding to each dimension on the vector is known. However, most of the learned protein representations are not interpretable/explainable. In other words, the meaning of a feature encoded in a dimension of the output vector is not known. For example, presence of a TIM barrel structure in a protein might be encoded in the 5th dimension of its representation vector, whereas, the molecular weight information may be shared between the 3rd and 4th dimensions. In the general field of data science, disentanglement studies try to associate the real properties of input samples with individual dimensions of the output vectors^62^. The disentanglement of protein representations is a new subject, and only a few representation model developers have explored this issue so far^44,49^, as a result, there is yet to be a systematic approach. Therefore, systematic benchmarking platforms are required for the standardized evaluation of protein representation model interpretabilities.

Most of the protein representation models so far are trained using only one type of data (e.g., protein sequence). However, protein knowledge is associated with multiple types of biological data, such as protein-protein interactions (PPI), post-translational modifications, gene/protein (co)expressions, etc., along with sequences. According to the best of our knowledge, only a few of the available protein representation models are trained with more than two types of data^29,63^.

Among the methods in our benchmark study, Mut2Vec^29,63^ harmonized multiple types of data (i.e., PPIs, mutations and biomedical texts), and produced more accurate results than many of the solely sequence based representations in GO BP and CC based PFP. We propose that the integration of additional types of protein related data may further augment the accuracy in predictive tasks. Furthermore, there is a clear requirement in the literature for holistic protein vectors that can effectively represent proteins from a generalized point of view, to be used for various different proteins informatics-related purposes. In our opinion, it may be possible to create these holistic representations by concatenating multiple representation vectors that were previously constructed using different types of biological data (as a means of pre-training), and training a new model using the integrated version of these vectors over high-level supervised tasks such as predicting biological processes and/or complex structural features (Fig. S6).

Protein design is one of the key challenges in biotechnology^64^. Rational protein design involves evaluating the activities and functions of many different alternative sequences/structures to provide the most promising candidates for experimental validation, which can be seen as an optimization problem^65^. The sequence space to be explored for this purpose is enormous. For example, the mean length of human proteins is around 350 amino acids, for which 20^350 different combinations exist. In the last three decades, computational approaches have been utilized for designing proteins with an increasing intensity, which produced promising results considering enzyme design^66–68^, protein folding and assembly^69^, and protein surface design to develop efficient antibodies^70^ and biosensors^71^. Most of these methods use the quantum mechanical calculations^72,73^, molecular dynamics^74,75^ and statistical mechanics^76,77^, all of which have exceptionally high computational costs^78^, and require expert knowledge. On the other hand, methods based on statistical heuristics demand less computing resources, but have lower performance.

Recent studies showed that artificial learning-based generative modelling can be employed for *de novo* protein design. Generative modelling is an approach, as opposed to discriminative modelling, in the machine learning domain^79^, where synthetic samples are produced, that obey a probability distribution learnt from real samples. To accomplish this task, it is required to learn the representations of samples in the training dataset. Recently, deep learning has become the key approach for generative model architectures^80^, which have been applied in various fields including protein/peptide design. For example, Greener *et al.*^81^ utilized variational auto-encoders to design metal-binding proteins. In another study, Gupta and Zou^82^ showed that generative adversarial networks (GANs) could be used for designing proteins through the construction of synthetic encodings of DNA sequences. In the work by Biswas *et al.*, variants of two different proteins (a fluorescent protein and a hydrolase) could successfully be designed with improved functional activity^50^. In another study, Tubiana *et al.* showed that proteins can be designed by defining preferred functions by conditioning a Restricted Boltzmann Machine-based protein representation model^49^. Furthermore, it was possible to generate direct 3-D coordinates of fullatom antibody backbones^83^ and to design peptides with anticancer properties^84^ (validated by *in vitro* experiments) with deep generative modelling. In the field of drug discovery and development, learned representations have been employed for molecular property prediction^85^, drug-target interaction prediction^86^ and *de novo* drug design^87^. These studies indicate that representation learning is critical for novel applications in both protein and ligand (drug) design.

We believe protein representation learning approaches will have an influence on various fields of the protein science with real-world applications, in the near future, thanks to their flexibility to integrate heterogeneous protein data at the input level (i.e., physical and chemical properties/attributes, functional annotations, etc.), and their ability to efficiently extract complex hidden features.

## Methods

In this section, we first explain different approaches in representing proteins, along with an evaluation of representation learning methods from a technical point of view. A section specifically dedicated to classical representation methods is provided under supplementary information section 1. We then summarize representation learning methods according to their training objectives and provide detailed information about those we included in our benchmark analyses (the rest of the protein representation methods are detailed in supplementary information section 3). Finally, we present methodological details regarding the datasets, modelling approaches, training and test procedures and performance metrics for each benchmark task (i.e., semantic similarity inference, ontology-based protein function prediction and drug target protein family classification). We shared the source code, models and datasets of this study in our repository (https://github.com/serbulent/TrainableRepresentationAnalysis) so that the data can be used by other groups for benchmarking new representation models and to compare the results with the ones provided in this study.

### Different Approaches for Representing Proteins

Feature vectors should ideally represent relevant properties of the data at hand (e.g., physical, chemical, or biological properties while representing proteins). For example, within the classical protein representation approach, a protein can be represented as a 2-dimensional numeric vector where the first dimension corresponds to the mean hydrophobicity value, and the second is the mean net charge^88^. Using these simple vectors as input, a classifier can be trained (Fig. S1a). In another example shown in Fig. S1b, the representation learning approach is employed, where Gene Ontology (GO) based functional annotations of proteins are used as the input data. An initial binary matrix that displays the associations between proteins and GO terms, is decomposed into latent protein and GO term matrices using matrix factorization. Afterwards, a predicted protein vs. GO term matrix is calculated with the dot product of the first, and the transpose of the second latent matrices. The error between the original and predicted matrices is used to update the parameters of the model during training. When the training is finished, each row of the finalized latent matrices is used as a feature vector that represent the respective protein or GO term. These feature vectors can then be used as input to other classification or clustering models for different predictive tasks.

In the domain of natural language processing (NLP), one of the first contemporary word representation learning methods, word2vec, was developed by Mikolov *et al.*^89^. Word2Vec is an unsupervised learning network that calculates a vector representation for each word in a text. In word2vec-like NLP models, the learning process is based on the co-occurrence of words. During the training of a word2vec model with the skip-gram architecture, the vector that represents the current word is optimized considering the correct prediction of surrounding words. In a successful representation model, e.g., the words “protein” and “gene” will be proximally located in the feature hyperspace (i.e., they are found to be semantically related to each other), since they are frequently observed together. Additionally, the model may also discover relationships between these words transitively, if, for example, the word pairs “protein - mRNA” and “mRNA - gene” are observed in the input text samples. Word2vec is an example of shallow data representation approaches, in which only one effective data processing layer (i.e., the hidden layer in an artificial neural network) is presented. Word2vec laid the foundation of many widely-used data representation learning methods available today, including protein representations. Later, deeper models, in which there are more than one effective data processing layer^90^, were developed and have achieved far better performance on NLP tasks^91^.

The first examples of learned protein representations were based on the word2vec algorithm ^4,92, 93,94,95,29,96,30^, most of which are still in use today. Since word2vec depends on word cooccurrence in a limited window, it ignores the larger context which may include critical semantic information. For protein sequence-based representations, this larger context can be the whole protein sequence. Another embedding method, doc2vec^42^ includes the whole context to some extent and performs better than word2vec on selected tasks. Several methods use doc2vec to represent proteins^5,27,97–101^. Also, deep language models, such as BERT^91^ and ELMO^46^ were originally developed for NLP, and later employed for protein representations^23,28^. Furthermore, Convolutional Neural networks (CNNs), having the ability to learn to summarize the data with adaptive filters, have been employed to represent proteins^23,63,86,102,103^. Additionally, architectures that are capable of inferring patterns from sequential data (e.g., protein sequences) using the attention mechanism^23,55^, such as Long Short-Term Memory (LSTM) neural networks^23,28,44,104,105^ and transformer based algorithms ^106^, are used in representation methods. However, transformer-based methods have shortcomings considering model explainability^107,108^. For this, Restricted Boltzmann Machines (RBM)^109,110^, with its self-recursive design, are used to construct explainable protein representation models^49^. Finally, hybrid approaches are utilized in the protein representation learning literature^103,111,112^. Most successful protein representations possess certain common hallmarks; and these are explained and discussed under supplementary information document section 2.

### Summary of the Evaluated Protein Representation Learning Methods

We group protein representation learning methods’ technical approaches (Fig. 2a), and objectives and applications reported in their respective publications (Fig. 2b). Here, we formed five main categories according to the application domains; *(i)* protein interaction prediction (essential for understanding molecular mechanisms and pathways), *(ii)* physicochemical feature prediction (important for protein engineering and drug discovery related tasks), *(iii)* genetic feature prediction, *(iv)* protein function prediction, and *(v)* structural feature prediction. Fig. 2b categorizes the main domains and specific application fields under each one. Methods with more than one objective were classified according to their major objective. The methods that we included in our benchmark study (i.e., LearnedEmbeddingVec^27^, SeqVec^28^, Mut2Vec^29^, Gene2Vec^30^, TCGA_Embedding^31^, ProtVec^4^, TAPE-BERT_Avg^23^, TAPE-BERT_Pool^23^, UniRep ^23^, and the two classical representations, APAAC ^32^ and k-sep-bigrams^33^) are described below, the rest of the methods are explained in supplementary information document section 3. All of these protein representation methods are summarized in terms of their technical aspects (e.g., learning approach, algorithm, etc.), input data types, vector sizes, objectives, applications, importance and available data repositories in Table 1.

Asgari *et al.*^4^ were one of the first to create a protein representation learning model to represent proteins by applying word2vec^89^ in a method they named ProtVec. The authors treated each protein sequence as a sentence, and each k-mer (i.e., k length amino acid sequence) as a word. Authors claimed that the method could be employed for different problems in protein biology including protein function prediction and protein interaction prediction. They evaluated the performance of ProtVec in predicting the mass, volume, polarity, hydrophobicity and charge of proteins, as well as its accuracy in disordered protein classification.

In the study by Yang *et al.*^27^, learned protein embeddings (i.e., representation vectors) with sizes ranging from 4 to 128 dimensions are constructed using the doc2vec algorithm^42^ on nonoverlapping k-mers. We will call this method “LearnedEmbeddingVec” in the rest of this study, as the Yang *et al.* did not provide a specific name. The authors measured the performance of LearnedEmbeddingVec and the effects of hyperparameter optimization on four protein property prediction tasks, namely channelrhodopsin (ChR) localization, cytochrome P450 thermostability, rhodopsin absorption wavelength, and epoxide hydrolase enantioselectivity with blocking design, to compare their model with baseline models (e.g., one-hot encoding and classical feature-based representations). Performance values were calculated using mean absolute error (MAE), a measure of variation between predicted and actual values; the Kendall rank correlation coefficient, which calculates the ordinal accuracy; and log-likelihood. For 3 out of 4 tasks, they report that LearnedEmbeddingVec provided the best performance in terms of at least one of these metrics. Authors also provide 2-D t-SNE visualizations, which were consistent with the reported results.

Rao *et al.*^23^ proposed a comparative study entitled “Tasks Assessing Protein Embeddings’’ (TAPE). The authors constructed three original sequence-based representation models based on; *(i)* Bidirectional Encoder Representations from Transformers (BERT)^91^ (we will call this TAPE-BERT), *(ii)* an unsupervised LSTM^113^, and *(iii)* ResNet^114^. These three models were trained on 32 million Pfam^115^ domains. Additionally, They evaluated two previously developed representations, UniRep^44^ and a supervised LSTM^104^. This study employs three groups of tasks which are; structure-based (i.e., secondary structure prediction and contact prediction), evolutionary (i.e., remote homology prediction) and protein engineering (i.e., fluorescence landscape prediction and stability landscape prediction). For structure-based tasks, an alignment-based representation (proposed as part of the baseline models) achieved the best score. In the evolutionary tasks, the pre-trained LSTM model had the top performance. Finally, for protein engineering tasks, TAPE-BERT was the best in terms of fluorescence landscape prediction and shares the top position with ResNet in terms of stability landscape prediction. Results indicated that no single method could dominate all of the benchmarking tasks. TAPE-BERT_Avg and TAPE-BERT_Pool are the two versions of TAPE-BERT, constructed by averaging and max-pooling the final hidden layer of the BERT model. In averaging, a mean value is calculated for each dimension of feature vectors that represents amino acids. In max-pooling, the maximum value of each dimension is used to create the final protein representation vector. We incorporated both TAPE-BERT_Avg and TAPE-BERT_Pool in our benchmark analyses.

In the study conducted by Du *et al.*^30^, the method Gene2Vec is proposed, where 200-dimensional vectors are calculated to represent genes, using skip-gram^89^. Hyperparameter tuning (e.g., vector size and window size optimization) was applied with the objective of maximizing the clusterdness of genes within MSigDB^51^ functional pathways. The input data, gene co-expression profiles, were gathered from the GEO database^116^. The major objective of the study is predicting gene-gene interactions (i.e., the genes acting in the same biological process), in which Gene2Vec was reported to be successful. Additionally, it was indicated that the model could summarize latent semantic information about genes by accurately representing functional similarities over tissue specific gene clusters. We employed gene representation vectors of Gene2Vec in our benchmarks by mapping them to canonical forms of their respective gene products (proteins). Kim *et al.*^29^ trained a mutation representation model named Mut2Vec. The aim of the proposed model was the classification of mutations according to their disease-causing effects. In Mut2Vec, mutation co-occurrence information, protein-protein interaction (PPI) networks (from BioGRID), and biomedical literature abstracts (from PubMed) were used to construct the representation model. Among alternatives, the model that utilizes the co-occurrence information via skip-gram^89^ was chosen as the finalized representation model. In the Mut2Vec workflow, first mutation cooccurrences and PubMed texts were used to calculate representation vectors. PPI data was integrated at the post-processing phase, using a retrofitting process similar to WordNet^117^. The authors stated that Mut2Vec could separate passenger and driver mutations successfully, and that, it produces promising results in the detection of new cancerous mutation candidates.

Heinzinger *et al.*^28^ used Embeddings from Language Models (ELMO), a bi-directional LSTM which is popular in the NLP domain^46^, to represent proteins using unlabelled protein sequence data. The authors aimed to solve the problem of global representation methods’ shortcomings in inferring information from the local context, with their method, SeqVec, using a smaller version of ELMO with 244k parameters, resulting in a significant speed advantage over up-to-date language models such as BERT^91^. However, their results show that the smaller model could not surpass the state-of-the-art methods, especially on sequence level tasks such as secondary structure prediction. On the other hand, the model produced competitive results for protein level tasks such as subcellular localization prediction. In our opinion, this can be attributed to the extensive training dataset and the successful detection of conserved sequence patterns by the LSTM.

Alley *et al.* developed the method UniRep^44^, a Multiplicative LSTM^35^ (mLSTM) backed character based representation. They tested UniRep on different, mostly protein engineering based, protein informatics tasks, including the classification of proteins based on their families and species, and the prediction of physicochemical properties and secondary structural elements. The results indicate that UniRep could create physicochemically meaningful clusters. Moreover, sequentially distant homologous proteins were clustered correctly. Finally, structural information could be extracted from UniRep, shown by the successful clustering of proteins based on SCOP^118^. These results were also verified using functional, evolutionary, and structural similarity labelled datasets such as HOMSTRAD^119^ and OXBench^120^. The authors have also shown that UniRep can predict protein stability and variant effects.

The study conducted by Choy *et al.* indicates that learned protein representations have potential for explaining molecular biological mechanisms of the cell and disease^31^. In the proposed method, first, a gene expression matrix of cancer samples was prepared using data from the TCGA database. The authors then applied a matrix decomposition with a fully connected neural network layer. Next, through matrix multiplication on these decomposed matrices, they created a predicted version of the original matrix. The error between the original and predicted matrices was used for backpropagation. The decomposed matrices consisted of gene-features and samples-features as dimensions. The authors showed that semantic relationships between samples and genes are conserved in their model. Even though the gene expression levels were not correlated, functionally related genes are observed in adjacent locations, when the multidimensional distance was calculated on the representation vectors. Additionally, when the representation vectors were inspected, it was seen that similar cancer types were clustered in the representation space to the extent that the authors claim that molecular subtyping of cancer was possible using the representations. We refer to this method as “TCGA_Embedding’’ throughout this paper, as the authors did not provide a specific name for their model in the original study.

Finally, two widely used classical representations (APAAC and k-sep-bigrams) were included to our benchmark, as baseline models. Amphiphilic Pseudo-Amino Acid Composition (APAAC) utilizes the physicochemical properties of amino acids together with amino acid compositions^32^, and k-sep-bigrams makes use of the evolutionary relationships between proteins^33^. We evaluate the performance of learned representations in comparison to these baselines, to assess the value added by the newer methods. Below, we look at these baseline methods in more detail.

A general issue in amino acid composition based classical representation methods is the difficulty of including residue order information. The Amphiphilic Pseudo Amino Acid Composition (APAAC)^32^ model proposed a solution to this problem by using sequence order coupling and hydrophobic correlations together. The model calculates a representation vector with 80-dimensions (by default), in which the first 20 represent the individual amino acid compositions, and the rest represent the hydrophobicity/hydrophilicity correlation factors. The APAAC method was found successful in predicting enzyme sub-families using a covariant-discriminant predictor^32^.

Evolutionary information is widely used in classical protein representations. In k-separated-bigrams method, row-type matrix transformations on position specific scoring matrices (PSSM), which are constructed using multiple sequence alignments generated from the query sequence and its homologs, are utilized for calculating the bigram transition probabilities between residues that are “k” positions apart from each other. The final representation vectors have the size of 400×1, each dimension representing a specific transition probability from one amino acid to another (20×20). The method was reported to be successful in predicting type IV secretion effectors^33^.

### Semantic Similarity Inference Benchmark

To construct the dataset of semantic similarity inference benchmark, we downloaded all human protein entries in the UniProtKB/Swiss-Prot database and their GO term annotations from the UniProt-GOA database in the 2019_11 release. The electronically inferred annotations, labelled with the “IEA” evidence code, were excluded from the dataset; leaving only the annotations reviewed by human experts. After that, we enriched the dataset by propagating the annotations to the parent terms of the asserted GO terms on the directed acyclic graph (DAG) of GO, according to the true path rule. Our finalized annotation dataset contained 14,625 distinct GO terms (3,374 of them belonged to molecular function - MF, 9,820 belonged to biological process - BP, and 1,431 belonged to cellular component - CC categories) and 326,009 annotations (75,884 of them belonged to MF, 154,532 belonged to BP, and 95,593 belonged to CC categories).

To be used as the ground-truth/reference data in this benchmark, we calculated the true GO-based semantic similarities between all proteins in our dataset independently for all GO aspects (i.e., MF, BP and CC) using Lin similarity in the GoSemSim package^121^. Lin similarity^34^ is based on Shannon’s information theory, which states that the information content (IC) of an event is negatively proportional to the observation probability (P) of the event. Information content (IC) is formulated as;

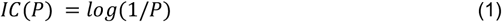

Another concept used in Lin similarity is the least common subsumer (LCS). LCS is the first common ancestor of the two GO terms when traveling to the root in the hierarchical GO graph. Hence, Lin similarity is defined as;

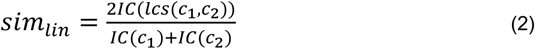

More information about the semantic similarity measures can be found in the literature^122^.

Next, we prepared four protein semantic similarity datasets (i.e., “all proteins”, “well annotated 500”, “well annotated 200” and “sparse uniform”) for each GO category (i.e., MF, BP, and CC), hence, twelve datasets were generated in total. The first dataset includes the pairwise GO-based semantic similarities between all proteins in our dataset (labelled as “all proteins” in the related figures). In this set, 3,077 proteins were used to calculate MF-based pairwise semantic similarities, 6,154 proteins were used for BP-based similarities and 4,531 proteins for CC-based similarities. In the “all proteins” dataset, there are numerous poorly annotated proteins, most of which contain insufficient information about their functional properties. This might introduce a bias in the similarity measurements. To mitigate this, we prepared additional subsets and ran the same analysis on them as well. The first subset, containing only the top 500 proteins sorted by the number of GO annotations (labelled as “well annotated 500” in the related figures). The second subset consists only of the top 200 such proteins (labelled as “well annotated 200” in the related figures). The similarity distribution is not uniform in the three datasets described above, creating very dense similarity score regions (Fig. S2) which significantly decrease the Spearman correlation values due to rank changes among the pairs with proximal similarities. This caused an accumulation around low correlation values that diminished the discriminative power of the measurements. To prevent this, we sampled every thousandth protein pair from the ranked list of pairwise similarities from the “well annotated 500” set to generate a uniformly distributed dataset. This final dataset contains 247 similarity scores between 40 different proteins (labelled as “sparse uniform” in the related figures). Thus, among our 4 datasets, “sparse uniform” is the most trivial one to predict and “all proteins” is the most challenging.

In the benchmark phase, we compiled the protein representation vectors for the human protein entries in our dataset using the selected representation learning methods, which are Gene2Vec^30^, LearnedEmbeddingVec^27^, Mut2Vec^29^, ProtVec^4^, SeqVec^28^, TCGA_embedding^31^, Tape_BERT_Avg^23^, Tape_BERT_Pool^23^ and UniRep^44^. Pre-calculated vectors, when available, were used directly, in other cases these were generated from their respective models. In addition, two classical representation methods (i.e., APAAC^32^ and k-sep-bigrams^33^) were included as baselines. Subsequently, we calculated the pairwise similarities between the proteins, using the compiled representation vectors. Cosine similarity, normalized Manhattan distance, and normalized Euclidean distance measures are used to evaluate pairwise similarity (normalized Manhattan and Euclidean distances are converted to similarities by subtracting them from 1.).

At this point, we had two pairwise similarity arrays at hand; the first one was calculated by taking the GO-derived semantic similarities between the proteins in our dataset into account (i.e., true semantic similarities), and the second one consisted of pairwise similarities calculated directly from the representation vectors.

Finally, to observe and to compare the performance of protein representation models for inferring these semantic similarities, we calculated the Spearman rank-order correlation^123^ values (explained below under “Performance metrics” sub-section) between the ranked lists of representation vector similarities and true semantic similarities.

### Ontology-based Protein Function Prediction Benchmark

The details of the dataset preparation procedure for the protein function prediction benchmark is explained below in six steps. For each GO category (i.e., MF, BP, CC);

1. We obtained human proteins and their GO term annotations from the “2019_10” version of UniProtKB/Swiss-Prot and UniProtGOA databases, respectively.
2. We excluded all electronically made annotations (evidence code: IEA) from the list of GO term annotations with the aim increasing the reliability of annotations and to prevent error propagation during prediction.
3. For each GO term, we created an individual list that includes the accessions of the annotated proteins, to be used in model training and testing via cross-validation. We filtered each protein list using the UniRef clusters^36^ by only selecting the representative protein entry from each cluster. UniRef provides protein clusters that are formed based on sequence similarity. We used UniRef50 clusters, to ensure that there are no protein sequences with more than 50% sequence similarity in each list. Here, the aim is to create train/test datasets without similar proteins that could otherwise introduce a bias to the analysis.
4. GO terms were grouped as either “low”, “middle”, or “high” according to the number of annotated proteins. GO terms with 2 to 30 annotated proteins were placed in the “low” group, terms with 100 to 500 annotated were placed in the “middle” group and terms with more than 1000 annotated proteins were placed in the “high” group. We deliberately left margins between groups to obtain a clear separation.
5. The specificity of the GO terms was determined as either “shallow”, “normal”, and “specific”. In the directed acyclic graph of GO, terms in the first ⅓ of the max depth of that branch were considered as “shallow”, terms in the second ⅓ of the max depth of that branch were categorized to “normal”, and the deepest ⅓ were placed to “specific” group. It should be noted that the max depth varies according to GO category.
6. Based on the combinations of groups constructed in steps 4 and 5; a total of 9 GO term groups (3×3) were formed for each GO category (i.e., MF-low-specific, BP-high-shallow and etc.), making a total of 27 groups (9×3). There were no GO terms that correspond to two of these groups (e.g., MF-high-specific and CC-high-specific), and therefore, these groups are left out of this analysis. Since most of the remaining 25 groups were highly crowded, we selected 5 terms from each group for further evaluation (4 groups already had less than five GO terms, thus, they were directly incorporated without further selection). We intended to select dissimilar GO terms to be able to generalize the results over the whole functional spectrum, as much as possible. For this, we calculated pairwise semantic similarities between GO terms using Lin similarity, and 5 most dissimilar terms were chosen for each group. The statistics of the finalized datasets are given in Table S2 and the identifiers of the selected GO terms are given in Table S3.

Using these datasets, prediction models were constructed (one for each group, mostly made up of 5 GO terms) for each protein representation model using the “Linear Support Vector Classification” module of the scikit-learn library^43^ within a multi-task modelling approach, making a total number of 275 prediction models (25 GO groups x 11 representation models). A 5-fold cross-validation was used to evaluate performance for each model. The hyperparameters of the SVM, for all models, were selected based on the default values; the regularization parameter (C) was set to 1.0, L2 norm was selected for error penalty, and the squared hinge was chosen for the loss function. Since the linear classification model is simple, we assumed that the effect of the hyper-parameter selection would be minimal.

### Drug Target Protein Family Classification Benchmark

To construct our family classification benchmark dataset, we employed ChEMBL database (v.25)^45^, which contains curated collections of drug/compound-target protein interaction data (i.e., bioactivities) to be utilized for experimental and computational research in drug discovery and development. Taking into account the hierarchical target protein categorization system presented in ChEMBL, we use 4 broad target protein families and grouped the rest of the targets as a fifth category (i.e., enzymes, membrane receptors, transcription factors, ion channels, and others). The number of proteins for each family and representation method is shown in Table S6. Small differences between the dataset sizes of different representation methods was due to the availability of vectors, and assumed to be negligible (the largest difference was around 3%). UniRef filtering was not used in this benchmark due to small sizes of original datasets (e.g., transcription factors class was composed of 82 human target proteins). The family information is used as class labels for the multi-task training of the target protein family classification model.

The stochastic gradient descent classifier-SGDClassifier-(i.e., a linear SVM) from the scikit-learn library^43^ was used with “OneVsRestClassifier” option for the multi-task classification. The classifier was used with default parameters; SVM for fitting the SGDClassifier, hinge as the loss function, and L2 norm as the error penalty. The model was trained and tested with 10-fold crossvalidation. The whole process was repeated 100 times, and the average results are reported.

### Performance metrics

In our semantic similarity inference benchmark, we used Spearman rank correlation^123^. For a sample with size *n* and the ranks of variables *rg_xi_*. and *rg_yi_*., Spearman rank correlation (*r_s_*) can be defined as:

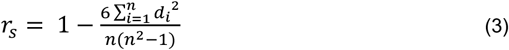

where difference between ranks for observations is defined by:

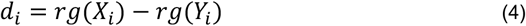

For ontology-based protein function prediction and drug-target protein family classification benchmarks, we mainly used recall, precision, F1-score, accuracy, Matthews correlation coefficient^124^ (MCC) and Hamming distance^125^ metrics, to evaluate the predictive performance of protein representation learning methods. The formulae of these evaluation metrics are given below:

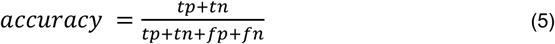

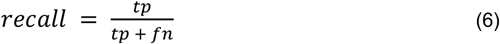

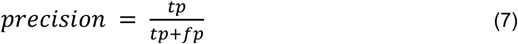

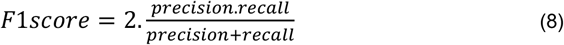

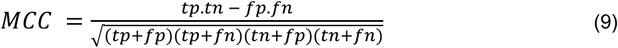

where *tp* denotes number of true positive predictions, *fp* denotes number of false positive predictions, *fn* denotes number of false negative predictions, *tn* denotes number of true negative predictions. Finally, Hamming distance (*D_H_*) is defined by:

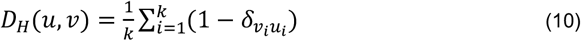

where *u* and *v* are 1-dimensional arrays of real and predicted class labels, respectively, $is the Kronecker delta function, and k is the vector dimension.

In ontology-based protein function prediction benchmark, F1-score and its components precision and recall are weighted inversely proportional to the class sizes. Since the classes were highly imbalanced, the weighting operation was required for an unbiased analysis. In both ontology-based protein function prediction and drug target protein family classification benchmarks, models were designed as multi-task (i.e., 5 GO terms are predicted by one function prediction model, and 5 proteins families are predicted by one family classification model). In ontology-based protein function prediction benchmark, the models were also designed as multi-label, where more than one GO term can be predicted to a test protein (since a protein can have more than one function). In this setting, a random predictor would produce a correct prediction once out of 32 cases (i.e., 2^5^ different combinations exist for a label vector of size 5×1, one of which is the true label vector). Whereas, models are designed as single-label in the drug target protein family prediction benchmark (since each protein can only belong one of the main families), meaning that a random predictor would produce a correct prediction once out of 5 cases (i.e., only 5 different combinations exist for a label vector of size 5×1, one of which is the true label vector).

## Availability

All of the datasets, results and the source code of this project are available for download at: https://github.com/serbulent/TrainableRepresentationAnalysis.

## Acknowledgements

This work was supported by TUBITAK (project no:318S218). We thank Dr Gizem Tatar (faculty member, KTU, Turkey) for reading and commenting on the manuscript, and to Gülbahar Merve Çakmak Şilbir (PhD Candidate, KTU, Turkey) for contributing to the drawing of figures.

## Author Contributions

S.U., T.D. and A.C.A. conceived the idea and planned the work. S.U. evaluated the literature, constructed representation vectors, and prepared the datasets and carried out the analysis for the semantic similarity inference benchmark. H.A. and S.U. prepared the datasets and carried out the analysis for the ontology-based protein family classification benchmark. M.A. and S.U. prepared the datasets and carried out the analysis for drug-target protein family classification benchmark. S.U., A.C.A. and T.D. have written the manuscript. T.D., A.C.A and K.T. supervised the overall study. All authors have approved the manuscript.

## Competing Interests statement

The authors declare no competing interests.

## Additional information

Supplementary information is available for this paper.

## Footnotes

† Biotechnology Market Size by Application (Biopharmacy, Bioservices, Bioagriculture, Bioindustries, Bioinformatics), By Technology (Fermentation, Tissue Engineering and Regeneration, PCR Technology, Nanobiotechnology, Chromatography, DNA Sequencing, Cell Based Assay), Industry Analysis Report, Regional Outlook, Application Potential, Competitive Market Share & Forecast, 2019 – 2025.

## Notes

https://github.com/serbulent/TrainableRepresentationAnalysis

### Competing Interest Statement

The authors have declared no competing interest.

## References

1. Dalkiran, A. et al. ECPred: a tool for the prediction of the enzymatic functions of protein sequences based on the EC nomenclature. BMC Bioinformatics 19, 334 (2018).

2. Dobson, P. D. & Doig, A. J. Distinguishing enzyme structures from non-enzymes without alignments. J. Mol. Biol. 330, 771–783 (2003).

3. Latino, D. A. R. S. & Aires-de-Sousa, J. Assignment of EC numbers to enzymatic reactions with MOLMAP reaction descriptors and random forests. J. Chem. Inf. Model. 49, 1839–1846 (2009).

4. Asgari, E. & Mofrad, M. R. K. Continuous Distributed Representation of Biological Sequences for Deep Proteomics and Genomics. PLoS One 10, e0141287 (2015).

5. Kimothi, D., Soni, A., Biyani, P. & Hogan, J. M. Distributed Representations for Biological Sequence Analysis. arXiv [cs.LG] (2016).

6. Nguyen, S., Li, Z. & Shang, Y. Deep Networks and Continuous Distributed Representation of Protein Sequences for Protein Quality Assessment. in 2017 IEEE 29th International Conference on Tools with Artificial Intelligence (ICTAI) 527–534 (IEEE, 2017).

7. Keskin, O., Tuncbag, N. & Gursoy, A. Predicting Protein-Protein Interactions from the Molecular to the Proteome Level. Chem. Rev. 116, 4884–4909 (2016).

8. Moult, J., Fidelis, K., Kryshtafovych, A., Schwede, T. & Tramontano, A. Critical assessment of methods of protein structure prediction (CASP)-Round XII. Proteins 86 Suppl 1, 7–15 (2018).

9. Ovchinnikov, S., Park, H., Kim, D. E., DiMaio, F. & Baker, D. Protein structure prediction using Rosetta in CASP12. Proteins 86 Suppl 1, 113–121 (2018).

10. Wang, S., Li, W., Liu, S. & Xu, J. RaptorX-Property: a web server for protein structure property prediction. Nucleic Acids Res. 44, W430–5 (2016).

11. Sureyya Rifaioglu, A., Doğan, T., Jesus Martin, M., Cetin-Atalay, R. & Atalay, V. DEEPred: Automated Protein Function Prediction with Multi-task Feed-forward Deep Neural Networks. Sci. Rep. 9, 7344 (2019).

12. You, R. et al. GOLabeler: improving sequence-based large-scale protein function prediction by learning to rank. Bioinformatics vol. 34 2465–2473 (2018).

13. Jain, A. & Kihara, D. Phylo-PFP: improved automated protein function prediction using phylogenetic distance of distantly related sequences. Bioinformatics 35, 753–759 (2019).

14. The Gene Ontology Consortium & The Gene Ontology Consortium. The Gene Ontology Resource: 20 years and still GOing strong. Nucleic Acids Research vol. 47 D330–D338 (2019).

15. Zhou, N. et al. The CAFA challenge reports improved protein function prediction and new functional annotations for hundreds of genes through experimental screens. Genome Biol. 20, 244 (2019).

16. LeCun, Y., Bengio, Y. & Hinton, G. Deep learning. Nature 521, 436–444 (2015).

17. Esteva, A. et al. A guide to deep learning in healthcare. Nat. Med. 25, 24–29 (2019).

18. Liu, L. et al. Deep Learning for Generic Object Detection: A Survey. International Journal of Computer Vision vol. 128 261–318 (2020).

19. Zhang, C., Patras, P. & Haddadi, H. Deep Learning in Mobile and Wireless Networking: A Survey. IEEE Communications Surveys & Tutorials vol. 21 2224–2287 (2019).

20. Zou, J. et al. A primer on deep learning in genomics. Nat. Genet. 51, 12–18 (2019).

21. Weiss, K., Khoshgoftaar, T. M. & Wang, D. A survey of transfer learning. Big Data 3, 1817 (2016).

22. Raffel, C. et al. Exploring the Limits of Transfer Learning with a Unified Text-to-Text Transformer. arXiv [cs.LG] (2019).

23. Rao, R. et al. Evaluating Protein Transfer Learning with TAPE. arXiv [cs.LG] (2019).

24. Brown, T. B. et al. Language Models are Few-Shot Learners. arXiv [cs.CL] (2020).

25. Huang, E. H., Socher, R., Manning, C. D. & Ng, A. Y. Improving word representations via global context and multiple word prototypes. in Proceedings of the 50th Annual Meeting of the Association for Computational Linguistics (Volume 1: Long Papers) 873–882 (aclweb.org, 2012).

26. Elnaggar, A. et al. ProtTrans: Towards Cracking the Language of Life’s Code Through SelfSupervised Deep Learning and High Performance Computing. arXiv [cs.LG] (2020).

27. Yang, K. K., Wu, Z., Bedbrook, C. N. & Arnold, F. H. Learned protein embeddings for machine learning. Bioinformatics 34, 2642–2648 (2018).

28. Heinzinger, M. et al. Modeling the language of life – Deep Learning Protein Sequences. Bioinformatics 540 (2019).

29. Kim, S., Lee, H., Kim, K. & Kang, J. Mut2Vec: distributed representation of cancerous mutations. BMC Med. Genomics 11, 33 (2018).

30. Du, J. et al. Gene2vec: distributed representation of genes based on co-expression. BMC Genomics 20, 82 (2019).

31. Choy, C. T., Wong, C. H. & Chan, S. L. Infer related genes from large scale gene expression dataset with embedding. Cancer Biology 2524 (2018).

32. Chou, K.-C. Using amphiphilic pseudo amino acid composition to predict enzyme subfamily classes. Bioinformatics 21, 10–19 (2005).

33. Wang, J. et al. POSSUM: a bioinformatics toolkit for generating numerical sequence feature descriptors based on PSSM profiles. Bioinformatics 33, 2756–2758 (2017).

34. Lin, D. & Others. An information-theoretic definition of similarity. in Icml vol. 98 296–304 (1998).

35. Krause, B., Lu, L., Murray, I. & Renals, S. Multiplicative LSTM for sequence modelling. arXiv [cs.NE] (2016).

36. Suzek, B. E., Huang, H., McGarvey, P., Mazumder, R. & Wu, C. H. UniRef: comprehensive and non-redundant UniProt reference clusters. Bioinformatics 23, 1282–1288 (2007).

37. Ryngajllo, M. et al. SLocX: Predicting Subcellular Localization of Arabidopsis Proteins Leveraging Gene Expression Data. Front. Plant Sci. 2, 43 (2011).

38. Deng, M., Zhang, K., Mehta, S., Chen, T. & Sun, F. Prediction of protein function using proteinprotein interaction data. J. Comput. Biol. 10, 947–960 (2003).

39. Coenen, A. et al. Visualizing and Measuring the Geometry of BERT. arXiv [cs.LG] (2019).

40. Clark, K., Khandelwal, U., Levy, O. & Manning, C. D. What Does BERT Look At? An Analysis of BERT’s Attention. arXiv [cs.CL] (2019).

41. Peng, Y., Yan, S. & Lu, Z. Transfer Learning in Biomedical Natural Language Processing: An Evaluation of BERT and ELMo on Ten Benchmarking Datasets. arXiv [cs.CL] (2019).

42. Le, Q. & Mikolov, T. Distributed Representations of Sentences and Documents. in 1188–1196 (PMLR, 2014).

43. Pedregosa, F., Varoquaux, G. & Gramfort, A. Scikit-learn: Machine learning in Python. the Journal of machine (2011).

44. Alley, E. C., Khimulya, G., Biswas, S., AlQuraishi, M. & Church, G. M. Unified rational protein engineering with sequence-only deep representation learning. Synthetic Biology e1005786 (2019).

45. Mendez, D. et al. ChEMBL: towards direct deposition of bioassay data. Nucleic Acids Res. 47, D930–D940 (2019).

46. Peters, M. E. et al. Deep contextualized word representations. arXiv [cs.CL] (2018).

47. Kim, Y., Jernite, Y., Sontag, D. & Rush, A. M. Character-aware neural language models. in Proceedings of the Thirtieth AAAI Conference on Artificial Intelligence 2741–2749 (AAAI Press, 2016).

48. Senior, A. W. et al. Improved protein structure prediction using potentials from deep learning. Nature 577, 706–710 (2020).

49. Tubiana, J., Cocco, S. & Monasson, R. Learning protein constitutive motifs from sequence data. Elife 8, (2019).

50. Biswas, S., Khimulya, G., Alley, E. C., Esvelt, K. M. & Church, G. M. Low-N protein engineering with data-efficient deep learning. Synthetic Biology 113 (2020).

51. Liberzon, A. et al. Molecular signatures database (MSigDB) 3.0. Bioinformatics 27, 1739–1740 (2011).

52. Anselmi, F., Leibo, J. Z., Rosasco, L. & Mutch, J. Unsupervised learning of invariant representations. Theor. Comput. Sci. (2016).

53. Bonassi, F., Terzi, E. & Farina, M. LSTM neural networks: Input to state stability and probabilistic safety verification. Learning for Dynamics (2020).

54. Bietti, A. & Mairal, J. Invariance and stability of deep convolutional representations. Adv. Neural Inf. Process. Syst. (2017).

55. Rives, A. et al. Biological structure and function emerge from scaling unsupervised learning to 250 million protein sequences. Synthetic Biology 7 (2019).

56. Zafrir, O., Boudoukh, G., Izsak, P. & Wasserblat, M. Q8bert: Quantized 8bit bert. arXiv preprint arXiv (2019).

57. Conneau, A. et al. XNLI: Evaluating Cross-lingual Sentence Representations. arXiv [cs.CL] (2018).

58. Brocchieri, L. & Karlin, S. Protein length in eukaryotic and prokaryotic proteomes. Nucleic Acids Res. 33, 3390–3400 (2005).

59. Sanh, V., Debut, L., Chaumond, J. & Wolf, T. DistilBERT, a distilled version of BERT: smaller, faster, cheaper and lighter. arXiv [cs.CL] (2019).

60. Bhargava, P. Adaptive Transformers for Learning Multimodal Representations. arXiv [cs.CL] (2020).

61. Merity, S. Single Headed Attention RNN: Stop Thinking With Your Head. arXiv [cs.CL] (2019).

62. Higgins, I. et al. Towards a Definition of Disentangled Representations. arXiv [cs.LG] (2018).

63. Öztürk, H., Ozkirimli, E. & Özgür, A. WideDTA: prediction of drug-target binding affinity. arXiv [q-bio.QM] (2019).

64. Burk, M. J. & Van Dien, S. Biotechnology for Chemical Production: Challenges and Opportunities. Trends Biotechnol. 34, 187–190 (2016).

65. Gainza, P., Nisonoff, H. M. & Donald, B. R. Algorithms for protein design. Curr. Opin. Struct. Biol. 39, 16–26 (2016).

66. Baker, D. An exciting but challenging road ahead for computational enzyme design. Protein Sci. 19, 1817–1819 (2010).

67. Röthlisberger, D. et al. Kemp elimination catalysts by computational enzyme design. Nature 453, 190–195 (2008).

68. Privett, H. K. et al. Iterative approach to computational enzyme design. Proc. Natl. Acad. Sci. U. S. A. 109, 3790–3795 (2012).

69. Chan, H. S., Shimizu, S. & Kaya, H. Cooperativity Principles in Protein Folding. in Energetics of Biological Macromolecules, Part E vol. 380 350–379 (Elsevier, 2004).

70. Lippow, S. M., Wittrup, K. D. & Tidor, B. Computational design of antibody-affinity improvement beyond in vivo maturation. Nat. Biotechnol. 25, 1171–1176 (2007).

71. Looger, L. L., Dwyer, M. A., Smith, J. J. & Hellinga, H. W. Computational design of receptor and sensor proteins with novel functions. Nature 423, 185–190 (2003).

72. Duan, Y. et al. A point-charge force field for molecular mechanics simulations of proteins based on condensed-phase quantum mechanical calculations. J. Comput. Chem. 24, 1999–2012 (2003).

73. Brunk, E. & Rothlisberger, U. Mixed Quantum Mechanical/Molecular Mechanical Molecular Dynamics Simulations of Biological Systems in Ground and Electronically Excited States. Chem. Rev. 115, 6217–6263 (2015).

74. Childers, M. C. & Daggett, V. Insights from molecular dynamics simulations for computational protein design. Mol Syst Des Eng 2, 9–33 (2017).

75. Hollingsworth, S. A. & Dror, R. O. Molecular Dynamics Simulation for All. Neuron 99, 1129–1143 (2018).

76. Camilloni, C. & Vendruscolo, M. Statistical mechanics of the denatured state of a protein using replica-averaged metadynamics. J. Am. Chem. Soc. 136, 8982–8991 (2014).

77. Huang, S.-Y. & Zou, X. Statistical mechanics-based method to extract atomic distance-dependent potentials from protein structures. Proteins 79, 2648–2661 (2011).

78. Pierce, N. A. & Winfree, E. Protein design is NP-hard. Protein Eng. 15, 779–782 (2002).

79. Ng, A. Y. & Jordan, M. I. On Discriminative vs. Generative Classifiers: A comparison of logistic regression and naive Bayes. in Advances in Neural Information Processing Systems 14 (eds. Dietterich, T. G., Becker, S. & Ghahramani, Z.) 841–848 (MIT Press, 2002).

80. Salakhutdinov, R. Learning Deep Generative Models. Annu. Rev. Stat. Appl. 2, 361–385 (2015).

81. Greener, J. G., Moffat, L. & Jones, D. T. Design of metalloproteins and novel protein folds using variational autoencoders. Sci. Rep. 8, 16189 (2018).

82. Gupta, A. & Zou, J. Feedback GAN for DNA optimizes protein functions. Nature Machine Intelligence 1, 105–111 (2019).

83. Eguchi, R. R., Anand, N., Choe, C. A. & Huang, P.-S. IG-VAE: Generative Modeling of Immunoglobulin Proteins by Direct 3D Coordinate Generation. Bioinformatics 29 (2020).

84. Grisoni, F. et al. Designing Anticancer Peptides by Constructive Machine Learning. ChemMedChem 13, 1300–1302 (2018).

85. Coley, C. W., Barzilay, R., Green, W. H., Jaakkola, T. S. & Jensen, K. F. Convolutional Embedding of Attributed Molecular Graphs for Physical Property Prediction. J. Chem. Inf. Model. 57, 1757–1772 (2017).

86. Öztürk, H., Özgür, A. & Ozkirimli, E. DeepDTA: deep drug-target binding affinity prediction. Bioinformatics 34, i821–i829 (2018).

87. Gómez-Bombarelli, R. et al. Automatic Chemical Design Using a Data-Driven Continuous Representation of Molecules. ACS Cent Sci 4, 268–276 (2018).

88. Uversky, V. N., Gillespie, J. R. & Fink, A. L. Why are ‘natively unfolded’ proteins unstructured under physiologic conditions? Proteins 41, 415–427 (2000).

89. Mikolov, T., Sutskever, I., Chen, K., Corrado, G. & Dean, J. Distributed Representations of Words and Phrases and their Compositionality. arXiv [cs.CL] (2013).

90. Schmidhuber, J. Deep learning in neural networks: An overview. Neural Networks vol. 61 85–117 (2015).

91. Devlin, J., Chang, M.-W., Lee, K. & Toutanova, K. BERT: Pre-training of Deep Bidirectional Transformers for Language Understanding. arXiv [cs.CL] (2018).

92. Wan, F. & Zeng, J. (michael). Deep learning with feature embedding for compound-protein interaction prediction. Bioinformatics e1004157 (2016).

93. Mejía-Guerra, M. K. & Buckler, E. S. A k-mer grammar analysis to uncover maize regulatory architecture. BMC Plant Biol. 19, 103 (2019).

94. Ng, P. dna2vec: Consistent vector representations of variable-length k-mers. arXiv [q-bio.QM] (2017).

95. Choi, J., Oh, I., Seo, S. & Ahn, J. G2Vec: Distributed gene representations for identification of cancer prognostic genes. Sci. Rep. 8, 13729 (2018).

96. Jaeger, S., Fulle, S. & Turk, S. Mol2vec: Unsupervised Machine Learning Approach with Chemical Intuition. J. Chem. Inf. Model. 58, 27–35 (2018).

97. Dutta, A., Dubey, T., Singh, K. K. & Anand, A. SpliceVec: Distributed feature representations for splice junction prediction. Computational Biology and Chemistry vol. 74 434–441 (2018).

98. Xu, Y., Song, J., Wilson, C. & Whisstock, J. C. PhosContext2vec: a distributed representation of residue-level sequence contexts and its application to general and kinase-specific phosphorylation site prediction. Sci. Rep. 8, 8240 (2018).

99. You, R., Huang, X. & Zhu, S. DeepText2GO: Improving large-scale protein function prediction with deep semantic text representation. Methods 145, 82–90 (2018).

100. Viehweger, A., Krautwurst, S., Parks, D. H., König, B. & Marz, M. An encoding of genome content for machine learning. Genomics 1533 (2019).

101. Kané, H., Coulibali, M., Abdalla, A. & Ajanoh, P. Augmenting protein network embeddings with sequence information. Bioinformatics 1080 (2019).

102. Schwartz, A. S. et al. Deep Semantic Protein Representation for Annotation, Discovery, and Engineering. Bioinformatics D36 (2018).

103. Asgari, E., Poerner, N., McHardy, A. C. & Mofrad, M. R. K. DeepPrime2Sec: Deep Learning for Protein Secondary Structure Prediction from the Primary Sequences. 705426 (2019) doi:10.1101/705426.

104. Bepler, T. & Berger, B. Learning protein sequence embeddings using information from structure. arXiv [cs.LG] (2019).

105. Strodthoff, N., Wagner, P., Wenzel, M. & Samek, W. UDSMProt: universal deep sequence models for protein classification. Bioinformatics 36, 2401–2409 (2020).

106. Vaswani, A. et al. Attention is All you Need. in Advances in Neural Information Processing Systems 30 (eds. Guyon, I. et al.) 5998–6008 (Curran Associates, Inc., 2017).

107. Jain, S. & Wallace, B. C. Attention is not Explanation. arXiv [cs.CL] (2019).

108. Brunner, G. et al. On Identifiability in Transformers. arXiv [cs.CL] (2019).

109. Smolensky, P. Information processing in dynamical systems: Foundations of harmony theory. https://apps.dtic.mil/sti/citations/ADA620727 (1986).

110. Hinton, G. E. & Salakhutdinov, R. R. Reducing the dimensionality of data with neural networks. Science 313, 504–507 (2006).

111. Oubounyt, M., Louadi, Z., Tayara, H. & To Chong, K. Deep Learning Models Based on Distributed Feature Representations for Alternative Splicing Prediction. IEEE Access 6, 58826–58834 (2018).

112. Mirabello, C. & Wallner, B. rawMSA: End-to-end Deep Learning using raw Multiple Sequence Alignments. PLoS One 14, e0220182 (2019).

113. Klapper-Rybicka, M., Schraudolph, N. N. & Schmidhuber, J. Unsupervised Learning in LSTM Recurrent Neural Networks. Artificial Neural Networks — ICANN 2001 684–691 (2001) doi:10.1007/3-540-44668-0_95.

114. He, K., Zhang, X., Ren, S. & Sun, J. Identity Mappings in Deep Residual Networks. in Computer Vision – ECCV 2016 630–645 (Springer International Publishing, 2016).

115. Finn, R. D. et al. The Pfam protein families database. Nucleic Acids Res. 36, D281–8 (2008).

116. Clough, E. & Barrett, T. The Gene Expression Omnibus Database. Methods Mol. Biol. 1418, 93–110 (2016).

117. Miller, G. A. WordNet. Communications of the ACM vol. 38 39–41 (1995).

118. Andreeva, A., Howorth, D., Chothia, C., Kulesha, E. & Murzin, A. G. SCOP2 prototype: a new approach to protein structure mining. Nucleic Acids Res. 42, D310–4 (2014).

119. Mizuguchi, K., Deane, C. M., Blundell, T. L. & Overington, J. P. HOMSTRAD: a database of protein structure alignments for homologous families. Protein Sci. 7, 2469–2471 (1998).

120. Raghava, G. P. S., Searle, S. M. J., Audley, P. C., Barber, J. D. & Barton, G. J. OXBench: a benchmark for evaluation of protein multiple sequence alignment accuracy. BMC Bioinformatics 4, 47 (2003).

121. Yu, G. et al. GOSemSim: an R package for measuring semantic similarity among GO terms and gene products. Bioinformatics 26, 976–978 (2010).

122. McInnes, B. T. & Pedersen, T. Evaluating measures of semantic similarity and relatedness to disambiguate terms in biomedical text. J. Biomed. Inform. 46, 1116–1124 (2013).

123. Spearman, C. The Proof and Measurement of Association between Two Things. Am. J. Psychol. 15, 72–101 (1904).

124. Matthews, B. W. Comparison of the predicted and observed secondary structure of T4 phage lysozyme. Biochim. Biophys. Acta 405, 442–451 (1975).

125. Bookstein, A., Kulyukin, V. A. & Raita, T. Generalized Hamming Distance. Inf. Retr. Boston. 5, 353–375 (2002).

126. Faisal, M. R. et al. Improving Protein Sequence Classification Performance Using Adjacent and Overlapped Segments on Existing Protein Descriptors. JBiSE 11, 126–143 (2018).

127. Asgari, E., McHardy, A. & Mofrad, M. R. K. Probabilistic variable-length segmentation of protein sequences for discriminative motif discovery (DiMotif) and sequence embedding (ProtVecX). Bioinformatics 707 (2018).

128. Mirabello, C. & Wallner, B. rawMSA: End-to-end Deep Learning Makes Protein Sequence Profiles and Feature Extraction obsolete. Bioinformatics 228 (2018).

129. Cohen, T., Widdows, D., Heiden, J. A. V., Gupta, N. T. & Kleinstein, S. H. Graded Vector Representations of Immunoglobulins Produced in Response to West Nile Virus. in Quantum Interaction (eds. de Barros, J. A., Coecke, B. & Pothos, E.) vol. 10106 135–148 (Springer International Publishing, 2017).

130. Qi, Y., Oja, M., Weston, J. & Noble, W. S. A unified multitask architecture for predicting local protein properties. PLoS One 7, e32235 (2012).

131. Melvin, I., Weston, J., Noble, W. S. & Leslie, C. Detecting remote evolutionary relationships among proteins by large-scale semantic embedding. PLoS Comput. Biol. 7, e1001047 (2011).

132. You, R. & Zhu, S. DeepText2Go: Improving large-scale protein function prediction with deep semantic text representation. in 2017 IEEE International Conference on Bioinformatics and Biomedicine (BIBM) 42–49 (2017).

