## Supplementary Information for "Evaluation of Methods for Protein Representation Learning: A Quantitative Analysis"

#### 1. Classical Protein Representation Methods

In classical protein representations, fixed sized numerical feature vectors are generated by applying predefined rules (statistical calculations in some cases) as a means of data transformation on previously known quantitative measurements of selected physical, chemical and/or biological properties either at the level of individual amino acids, or sub-sequence fragments. Classical methods are model-driven, as these predefined rules are determined by the expert according to the known properties of biomolecular systems, as opposed to the data-driven approach used in learned representations, where the data at hand is directly utilized for extracting information in an automated fashion.

Physicochemical properties of amino acids are widely utilized in early classical protein representation approaches, as they can be easily obtained using the sequence, and they correlate well with the structural and functional properties of proteins. To the best of our knowledge, one of the first studies in which proteins are represented as vectors was conducted by Klein *et al.*[1]. In this study, proteins are represented using the number of hydrophilic and hydrophobic residues, to detect and classify membrane-spanning proteins. In another study, proteins are represented using an 18-dimensional vector, which is calculated based on hydrophobicity/hydrophilicity and appearance of short characteristic amino acid patterns such as signal peptides[2], in which functions of proteins are predicted with high accuracy based on 26 different groups.

One of the first applications of using high throughput data for representing proteins is by Liao and Noble[3], where the authors attempted to detect the structural and taxonomic relations between proteins using support vector machines (SVM). They represent proteins with fixed vectors of real numbers composed of pairwise sequence similarity scores against a corpus of proteins. The algorithm considers both positive and negative samples in the vectors to represent proteins.

Chen *et al.* proposed iFeature[4], a Python package for constructing structural and physicochemical feature based protein feature vectors, solely using sequences as input. The tool incorporates 53 different representation methods including amino acid composition, sequence order, secondary structure and protein disorder-based approaches.

Another widely used input to construct protein representations are the evolutionary relationships. It is possible to identify sites/regions with critical importance (e.g., functional regions) using evolutionary conservation between homologs. In their respective study, Wang *et al.* proposed a tool, POSSUM (both as an online service and a stand-alone version), for constructing protein feature vectors from 21 different position specific scoring matrix (PSSM) based classical protein representation methods, where the initial input is the protein sequence. The resulting vectors of 21 PSSM-based methods are evaluated on the prediction of type IV secretion effectors[5]

A variety of biological data types are employed to represent proteins using the classical approach. Here, we grouped the top-performing protein function prediction methods in the CAFA2 challenge[6], in terms of their input type, together with the data transformation approach they utilized for constructing their quantitative vectors (supplementary material Table S1).

### 2. Traits of Successful Protein Representations

Traits of good data representations are first defined in a study by Bengio *et al.*[7] and it is still widely accepted in the representation learning literature. In this section, we evaluated these properties in the context of protein representations.

**Smoothness:** If small changes on the input sample cause small changes in the output representation vectors, then the representation can be considered as smooth. Evaluating this information in terms of protein representations; small perturbations in the protein sequence (e.g., single residue variations between the amino acids with very similar physicochemical properties, on a non-critical/non-conserved region of the protein sequence) generally do not cause significant changes in the structure and function of the protein, hence, a successful protein representation should be smooth. However, if there is a single amino acid variation at a critically important region of the protein sequence and structure, such as a binding site, the variation should be observed on the representation vector as well, to reflect the change in the functional traits of the protein.

**Explanatory factors:** Explanatory properties are important for representation vectors. When the biological, chemical or the physical feature that corresponds to a specific dimension on the representation vector is known (e.g., the $n^{th}$ dimension of a protein representation vector corresponds to the hydrophobicity value), feature importance, which is just a quantitative measure of how much that specific feature contributes to the task at hand (e.g., prediction the function of a protein), can be associated with a real-world biological/chemical/physical property. Another advantage is that the vector can be reused partially with only its relevant selected features. Explanatory properties can also be utilized to construct more compact representation vectors. Explainability is usually not a problem for classic representations since the calculated values that correspond to different biological/chemical/physical features are just concatenated to generate the feature vectors; however, trained representation vectors are not inherently explanatory. Nevertheless, there are methods to uncover the explanatory features of a high dimensional representation[8–10]. When the explanatory features are discovered, it is also possible to build a task-specific low dimensional representation from the original high dimensional vector. Moreover, these features may be used towards completely different tasks such as the design of novel proteins with desired properties[11].

**A hierarchical organization of explanatory factors:** Proteins can be classified using hierarchical organizations. Hierarchical classification systems for proteins such as SCOP[12] and CATH[13] are widely used in the protein informatics domain. In these systems, hierarchical organizations are created based on the experimental knowledge on the proteins. For example, in SCOP classes are groups created from fundamental secondary structures such as alpha-helices or beta-sheets, moreover, folds are groups placed under the classes and consisted of combinations of secondary structures (e.g. “7-bladed beta-propeller” - consists of seven 4-stranded beta-sheet motifs, meander). Similarly, the hierarchical organization of explanatory factors contributes to the development of modular representation models for learned representation vectors. The modularity might be important for the reusability of representation vectors. For example, some features of the protein representation vectors can be used to predict high-level entities such as the SCOP classes (e.g. “All alpha proteins”) and other features might be used to predict more specific entities such as SCOP families (e.g. “Eukaryotic proteases”).

**Shared properties across different tasks**: A protein representation model, which was trained based on a specific task (e.g., contact map prediction), should be used towards other tasks (e.g., protein function prediction) as long as there is a biological, chemical or physical relationship between these tasks.

**Natural clustering**: Manifolds can be defined as lower-dimensional representations of information and this dimensionality reduction can be useful for data processing such as noise reduction. When the probability distribution of the training data has high-density regions, a manifold can be discovered along these regions which lower the dimension of the data representation. Naturally, low-density regions are observed between these manifolds which are similar to valleys between mountains. These low-density regions can be used to cluster samples. Manifolds are also important for defining simple/linear dependencies between features of the representation vector (also known as the **simplicity of factor dependencies**). The output of a successful representation model should be easily separable using a simple linear classifier/regressor, even though the input samples have complex and non-linear relations between each other. In the benchmarks of this study, we use linear classifiers on protein representation vectors to classify them. Hence, a good protein representation can be naturally clustered.

**Sparsity**: In general, only a few of the dimensions in a representation vector includes information relevant to a task of interest. From a statistical point of view, most of the factors are not sensitive to small changes. This property manifests itself as sparsity in the representation vectors. For example, active sites of a protein cover only a small fraction of the protein sequence, but an amino acid substitution in this region might cause significant changes in protein function, and this information may be captured by a sparse feature on the representation vector. This property can be utilized for interpreting/explaining the trained models by associating the output features with certain biological/physical/chemical properties of input proteins.

### 3. Objective-based Classification of Protein Representation Methods

In this section, we grouped and elaborate protein representation learning methods according to the objectives and applications reported in their respective publications (Fig. 2b). The methods that we included in our benchmark study (i.e., LearnedEmbeddingVec[14], SeqVec[15], Mut2Vec[16], Gene2Vec[17], TCGA_Embedding[18], ProtVec[19], TAPE-BERT_Avg[20], TAPE-BERT_Pool[20], UniRep[20], together with two classical representations: APAAC [21] and k-sep-bigrams[5],) are explained in the Methods section of the main text, so the information about them is not repeated below.

#### 3.1. Methods for Physicochemical Feature Prediction

State-of-the-art NLP methods are gaining importance in the protein representation domain with their context-based dynamic inference capabilities. Rives *et al.*[22] uses one of these, Bidirectional Encoder Representations from Transformers (BERT)[22], and adapts it to the protein representation domain by predicting *masked* (hidden) amino acids on sequences where 15% of amino acids are hidden. Using 250M protein sequences from the Uniparc database[23], the model can successfully predict the masked amino acids' features such as polarity, charge, hydrophobicity; and protein secondary structure and activities such as contact and variant effects. The authors also showed that 25M samples would have been sufficient to achieve performance results on par with 250M samples. Moreover, the study results indicated that the model is far better than the baseline representation models, such as random and n-gram models. The model was also tested against an untrained LSTM and shows better performance. When internal mechanics were inspected using orthology based tests, it was observed that the BERT model-based phylogenetic organization has a good correlation with the natural phylogenetic organization even for long homologous proteins. The model was also tested for vectorial stability and protein similarity prediction. For the vectorial stability task, at first, the average distance between species is calculated in the representation space. This vector distance is added to the source protein's representation vector, and ortholog proteins are searched around the new position. The representation's vectorial stability is measured based on the precision of this search in finding true orthologs. Finally, one primary application of this representation is prediction of mutational effects which includes two subtasks: intra-protein variant effect prediction using known mutations of the target protein and generalization of mutational fitness landscape for new proteins without any prior knowledge. The proposed representation model performs on par with thre state-of-the-art variant effect prediction methods, without any added knowledge about the protein, such as the structure.

The rest of the methods, where the objective is the prediction of physicochemical features (e.g., ProtVec, LearnedEmbeddingVec and TAPE_BERT models) are summarized in the Methods section of the main text.

3.2. Methods for Sequence-based Feature Prediction

To the best of our knowledge, the first application of learned protein representations was conducted by Melvin *et al.*[24] with the aim of developing a search algorithm for protein sequences. They trained a 2-dimensional protein representation vector, by integrating 3-D structural similarity and protein class label information. The accuracy of the method was reported to be notably higher compared to known alignment-based homology search methods. The method was also trained and tested using SCOP (Structural Classification of Proteins) labels. Moreover, it was shown that a faster protein sequence search is possible with simple protein representation vectors.

Qi *et al.*[25] exploited multi-task learning and developed a deep learning framework for training a protein representation model to predict various local protein properties. Previous studies showed that multi-task learning is advantageous since information extracted from different features creates a synergy, which leads to better performance[26]. The proposed model learned and predicted the secondary structure, signal peptide transmembrane topology, solvent accessibility, and protein/DNA binding residues simultaneously. In the first step of the method, a protein representation model was trained to predict naturally occurring protein sequences. During the second step, sequential feature extraction was applied to capture information at flanking regions of amino acid sequences. As a third step, a neural network composed of feed-forward layers does the classification job. Finally, a post-processing step was added using a Hidden Markov model to utilize information in position-based patterns, such as repeated sequences. The study was critical since the authors used protein representation learning models and multi-task labelling for protein feature prediction, and achieved better performance compared to previous studies.

In the study conducted by Kimothi *et al.*[27], the authors proposed a context-aware protein representation method based on doc2vec[28] algorithm, and named it “seq2vec”. The authors claimed that a previous method, ProtVec[19], which employs word2vec algorithm, does not fully capture the order information in the protein sequence. One important shortcoming of word2vec based models is their lack of global context awareness, which means that the order of the words is not considered in terms of the whole document in which the word exists, during the calculation of the representation vector. The authors solved this problem using the doc2vec algorithm, which calculates a vector for each word using neighbouring vectors and a document vector. The document vector represents features of the whole document, such its topic. According to the results of the study, seq2vec approach is notably better than ProtVec on protein family classification task.

In a follow-up study to ProtVec by Asgari *et al.*[29], a variable-length protein sequence segmentation approach is introduced. In the proposed method, ProtVecX, the byte-pair encoding[30] which had also been used in the field of neural machine translation, was adapted to the protein sequence representation domain. Also, as one of the first applications in this domain, a baseline is defined using k-mers occurrences. Ablation studies indicated that this is a successful baseline for protein classification that can be used in future studies as well. It is also important to note that there still is a strong requirement for baseline models in the domain of protein representation learning. ProtVecX, could not display a significant performance advantage against the k-mer-based method on protein classification tasks.

Xu *et al.*[31] exploited learned protein representations in their method called PhosContext2vec, with the aim of recognizing phosphorylation sites on protein sequences. Phosphorylation is one of the fundamental control mechanisms for the cells; thus, the identification of phosphorylation sites is a critical problem in protein science. In PhosContext2vec, both word2vec and doc2vec algorithms were applied together with overlapping n-grams, which produced a better performance compared to non-overlapping n-grams. For the identification of the phosphorylation sites, the support vector machine (SVM) algorithm[32] was utilized. The model used learned representation vectors and six other protein residue-level features such as Shannon entropy, relative entropy, disordered protein regions, secondary structures, Taylor's overlapping properties, and the average cumulative hydrophobicity. The results indicated that word2vec and doc2vec were successful in different cases, thus complementing each other, and neither one was superior in terms of the overall performance. When PhosContext2vec was compared with other phosphorylation site prediction methods on semi-independent tests, it was shown that PhosContext2vec outperforms or competes with them on different cases.

The method D-Space (Deep Semantic Protein Annotation Classification and Exploration[33]) is based on a convolutional neural network (CNN), which produces 256-dimensional representation vectors from protein sequences. In the respective study, these vectors are used to predict labels from 13 different sources including PFAM[34], InterPro[35], EC Number[36], GO[37] and PROSITE[38]. Multi-task modeling is one of the solutions proposed and applied for the scarcity of labelled data[39–41]. When the representation model is tested, it is observed that over 400.000 proteins can be grouped in correlation with their OrthoDB[42] cluster label. OrthoDB is a database of orthologous protein-coding genes, and the correlation is assumed to be a marker of the accuracy of D-Space. The method's speed and sensitivity were tested on 109M protein entries from the UniprotKB, and compared against the results of a standard BLAST search. The results showed that the proposed method is fast (i.e., 5 seconds to find homologous sequences of a query protein, on average, compared to several minutes when BLAST is used). A correlation analysis between sequence identity and representation vector similarity supported the results with R=0.84 with p-value<2.2e-16. The study also included promising results on searching functionally related proteins and variant effect analysis; however, these results were mostly case-based and insufficient to infer generalization.

Cohen *et al.*[43] used vector symbolic architectures[44] with a set of quantum related compositional operators to generate protein representation vectors. These orthographic vectors are able to represent words or k-mers and employed to overcome the dependency of protein representations on the exact locations of amino acids. Different physicochemical properties of amino acids are encoded into these vectors to represent the proteins. The representation and similarity measures are tested using immunoglobulin (Ig) sequences gathered from patients infected with the West

Nile Virus (WNV). The task was the identification of WNV-specific clonal lineages between thousands of different Ig sequences. The study showed that the best overall results were obtained by the proposed method, where the alternatives were models based on bag-of-amino-acids and bag-of-properties. This approach may be promising not only in terms of the provided results but also considering the potential applications on quantum computers.

It is known that the smallest fully independent functional building block of a protein is a domain. In the study conducted by Viehweger *et al.*[45] authors use protein domains where a word vector model is trained via the doc2vec[28] algorithm, using protein domain sequences as words. The proposed method is called nanotext. The study was the first to train a protein representation using metagenomes (from 32 thousand genome assemblies). Training models for metagenomes is a critical issue since metagenomes have billions of records and most of their functions are unknown. The accuracy of the representation model was tested with the semantic odd man out (SOMO) approach, which can be defined as identifying the odd word in a context, where the method achieved more than 99% accuracy. When the representations were visualized with t-SNE, it was observed that clusters were correlated with the enzymatic functions of proteins. When genome vectors were used instead of domains, it was noted that nanotext could infer the taxonomic information, even for highly incomplete genomes. Finally, using nanotext, various features of bacteria, such as the culture medium and water temperatures, in which the bacteria were sampled, could successfully be predicted. The results indicated that there might be many other potential applications of protein representations.

You and Zhu[46] utilized text data for automated protein function prediction in terms of gene ontology-based annotations, in their method called DeepText2GO. The texts are one of the least exploited type of data in protein representation learning since the inference of relevant information is difficult. The main novelty of DeepText2GO was that the authors used the text data (trained on abstracts of MEDLINE[47]) and the homology information together, since each was reported to be successful in different tasks (i.e., sequence-based homology gave a better performance on molecular function prediction, and text-based information was more successful in predicting biological processes and cellular components). In DeepText2GO, TF-IDF and doc2vec[28] methods were used for the text-based classification, and BLAST-KNN and logistic regression were used for the sequence-based classification on InterPro[35] where domains, families, and motifs were used. The algorithm produced comparable results to the state-of-the-art methods on the CAFA2[6] benchmark dataset. One interesting finding of the study was the better performance of TF-IDF over doc2vec, which was also consistent with the latest study of Asgari *et al.*[48]. This finding indicates the shortcomings of static word vectors and the requirement for more sophisticated representation models.

Jaeger *et al.*[50] developed a cheminformatics-based application of word2vec. The method, named Mol2Vec, was trained as an unsupervised representation model. Here, the aim is to create a representation for molecular substructures that could be used for various tasks, including drug discovery and repositioning. SMILES[52] representations and Morgan fingerprints[51] were used to create the input to the method. Continuous Bag-of-Words (CBOW) and skip-gram algorithms of the word2vec[53] were tested on different tasks (e.g., regression-based prediction of solubilities, classification of mutagenic and non-mutagenic compounds, and the prediction of compound toxicities), and the most successful one was selected for each task to represent molecules. Also, a combination of ProtVec[19] and Mol2Vec was created and tested (named as PCM2Vec). It was shown that PCM2Vec could successfully predict kinase bioactivities. Also, both Mol2Vec and PCM2Vec could attain success comparable to the state-of-the-art methods at every task. Finally, Mol2Vec was used to visually illustrate the molecular substructures.

There are also additional studies aiming for the prediction of sequence-based features, that are worth mentioning. In the study conducted by Faisal *et al.*[49], the authors divided the protein sequence into segments and calculated position-free descriptors, such as amino acid composition, dipeptide composition, and normalized Moreau-Broto, on each segment, together with position-based numerical features. When implemented with SVM algorithm for feature selection and classification, the method scored a slightly higher accuracy over ProtVec in terms of protein family prediction. Kane *et al.*[54] aimed to predict the functions of proteins by augmenting protein sequence representation vectors with protein-protein interactions. Their most successful representation vectors were compared to random representation vectors, and there was a slight increase in terms of the area under the ROC curve (from 0.5 to 0.6). Strodthoff *et al.*[55] showed that a pre-trained protein representation could transfer knowledge from large unlabelled datasets to labelled small data sets. This approach was tested for enzymatic function prediction using EC numbers. Moreover, their AWD-LSTM[56] based model performed well compared to other state-of-the-art methods for protein function prediction and homology detection.

#### 3.3. Methods for Interaction Prediction

Wan and Zeng[57] exploited vectorial representations of proteins for the prediction of interactions between compounds and target proteins. The authors employed a 3-D structure-free drug-target interaction prediction method using latent semantic analysis and word2vec[53] algorithm to learn both the compound and protein representation vectors. In this work, the substructures of compounds (which are created using Morgan fingerprints) and k-mers of protein sequences were considered as words, and the whole compounds and protein sequences were considered as sentences. The method uses skip-gram with negative sampling. After constructing the independent compound and protein representations, authors concatenated the vectors and input to a deep neural network, to predict the interaction between the compound and protein of interest. The performance of the model was evaluated using known interactions of drugs and drug-candidate compounds against proteins. Well known databases such as ChEMBL[58] and DrugBank[59] were used as data sources. According to the authors, results are promising and better than conventional approaches.

In their method called DeepDTA[60], Ozturk *et al.* created learned representation vectors for proteins and compounds to predict the drug-target binding affinities. The authors used SMILES notations of compounds and protein sequences to create the representation vectors. Using character-based representation vectors for ligand representation, instead of words, is novel since most of the representation learning methods utilized the latter for similar tasks. DeepDTA utilizes CNN to train the representation vectors. Afterwards, the representation vectors are aggregated and given to a fully connected deep neural network for prediction. Concordance Index (CI) and mean square error (MSE) measures were used as evaluation metrics on two different benchmark datasets. The authors also conducted a performance analysis, which indicated that DeepDTA had a lower MSE and a higher CI value than the compared methods. In a follow-up study to from the same group, authors utilized protein domains, motifs and the maximum common substructure information within a similar algorithmic framework and developed the method WideDTA[61]. The results indicated that this data did not contribute to the model performance, but interestingly, using only domain and motif data produced competing results to using the full protein sequence, which indicates that a significant amount of ligand binding information is located in domain/motif regions.

Yao *et al.*[62] developed a deep learning pipeline named DeepFE-PPI to predict protein-protein interactions (PPI). In this method, proteins are represented using a word2vec[53] based algorithm (skip-gram) named Res2vec, which calculates a vector for each interaction between residues. Then, multiple dense layers were formed to predict PPIs. PPI networks belong to S. Cerevisiae, and human were used to test the proposed method. According to the authors, DeepFE-PPI achieved the best performance for most of the test cases.

Zhang and Kabuka[63] claimed that they developed the first deep multi-modal PPI prediction system. The authors stated that traditional PPI prediction techniques mostly rely on the protein sequence. In the proposed method, features based on the whole protein context (amino acid composition), protein sequence (using a stacked autoencoder), and PPI graph (using continuous bag-of-words algorithm) are utilized together to predict PPIs. Also, protein family prediction capabilities of the output representation were tested. The results indicated that the performance of the proposed approach was reported to be better compared to the state-of-the-art methods.

#### 3.4. Methods for Structural Feature Prediction

Nguyen *et al.*[64] developed DeepCon-QA, a deep convolutional neural network model that uses continuously distributed protein representation (ProtVec) and protein profiles as input to predict a protein distance matrix that represents protein structure for quality assessment of predicted structures. Quality assessment is part of protein structure prediction process which can be defined as measuring the similarity between the true and the predicted 3-D protein structures. The study results indicated that the utilization of protein representation vectors based on word2vec models created a significant performance improvement compared to using the protein profiles alone. It was also shown in the paper that DeepCon-QA competes with the state-of-the-art quality assessment methods. Hence, it is possible to comment that protein representation models have the potential for various types of *in silico* protein analysis tasks.

Mirabello and Wallner[65] trained protein representation models using a deep learning architecture and the multiple sequence alignment (MSA) data. The model predicted the secondary structural elements, relative solvent accessibility, and protein contact maps. In the first step, the input MSA was represented with an embedding layer. After a 2-D CNN with pooling was applied, an LSTM was employed to generate the predictions for secondary structures and relative-solvent accessibility. Besides, another CNN was employed for the contact-map prediction. Bias due to homology contamination (i.e., precise separation of training and test sets according to protein sequence similarity and protein family classification) were also taken into account and filtered during the model development phase. The results were indicated that the proposed approach was successful for all of the above-mentioned prediction tasks. The method could compete with the state-of-the-art approaches in protein contact map prediction, based on the benchmarks.

Asgari *et al.*[48] conducted a comprehensive study on the secondary structure prediction task. The authors evaluated one-hot vector representations, biophysical scores of amino acids, amino acid protein vectors, contextualized embeddings, and a Position-Specific Scoring Matrix (PSSM) as input. On the model side, various combinations of CNNs and LSTMs were applied to predict secondary structures in the CB513 Q8[66], which is a challenging dataset with eight different classes. The results showed that models that combine biophysical features, one-hot encoding, and PSSM achieved the best results. Still, it was shown that most of this accuracy (more than 99%) originated from the PSSM. On the model side, the ensemble of 100 different neural networks achieves the highest performance, but similarly, most of this score (more than 99%) originated from CNN-BiLSTMs. Finally, an inspection of confusion matrices showed that prediction accuracy decreased dramatically in the border regions of protein secondary structures.

There have been various efforts[15,19,67] to predict the structural features of proteins with unsupervised representation learning, but Bepler and Berger[68] took these efforts one step further. The authors developed a Bidirectional LSTM model that was trained on labelled structure data. A critical problem in sequence-based protein representation learning model development is the loss of positional correspondence during the representation vector calculation. Authors proposed a new solution to this problem with a new similarity measure they call “soft symmetric alignment” (SSA), a symmetrisation of the directional alignment commonly used in attention mechanisms. Using SSA, the model achieved the best structural similarity classification performance based on SCOP[69]. Moreover, the paper states that the developed model also produced improved contact and transmembrane prediction results.

The study conducted by Tubiana *et al.*[11] has addressed two critical problems in the protein representation learning domain. The first one was the interpretability of representation models. Since most of the models in this domain are based on black-box deep learning methods, it is hard to interpret and understand the internals. Moreover, finding the essential features of a protein representation vector is also an intricate effort. The second issue is related to the trend towards huge models[70]. Although there are ablation studies made for preventing overfitting, these models might tend to memorize patterns. Ramajuan *et al.* [71] showed that even with random weights, deep neural network models include optimal subnetworks, representing any mathematical function. The authors, Tubiana *et al.* used Restricted Boltzmann Machine (RBM) to model protein sequences using multiple sequence alignments of protein families. The network learned stochastic functions, which defined a two-way mapping between the protein sequence and the representation space. Since RBM depends on the Boltzmann distribution, the interpretability of the network is not a complex problem, as it is in black-box methods. Also, RBM’s low number of parameters reduced the training cost. The study results were also remarkable, in that, using the statistical base of the RBM, that it was possible to design proteins. Moreover, they demonstrate that it is possible to activate or deactivate the desirable features of a protein by conditioning the model. In the application phase, an accurate contact prediction could be made using the proposed model. In this study, a new type of activation function, a double Rectified Linear Unit (dReLU), was introduced, which contributed to the success of the model.

#### 3.5. Methods for Genetic Feature Prediction

Oubounyt *et al.*[72] exploited word2vec[53] and doc2vec[28] based sequence representation models to predict the percentage of splicing inclusion (PSI) in the context of alternative splicing. Using datasets from different tissues, the authors indicated that sequence representation learning models may be effective in predicting PSI. In this method, learned sequence representation vectors are given to the Inception Network[73] for PSI classification (as low, medium or high), and after that, PSI calculation with regression. The results of the study are indicated that word2vec and doc2vec models could capture similar features, and the deep learning model trained using these features outperformed the state-of-the-art methods for PSI calculation.

In their study, Dutta *et al.* [74] utilized word embeddings to calculate representation vectors using word2vec[53] and doc2vec[28]. The study aimed to solve the RNA-Seq data-based intron boundary recognition problem, since RNA-Seq suffers from misalignment of short reads. During the model development, 3-mers of the genetic sequences are considered as words, sequences with 2000 nucleotides are used as sentences, and splice junction sequences are used as the context. After the representation model is trained, a simple multilayer perceptron is used to detect the splice junctions and to annotate intron boundaries. The accuracy of the doc2vec based model was found to be better than word2vec. The results of the study also indicated that classification after representation training was invariant to class imbalance problem.

Mejía-Guerra and Buckler[75] developed a model to represent k-mers and classified different regions of the genetic sequences either as regulatory or random by learning complex patterns in regulatory regions. The authors employed word2vec and bag-of-words methods to define k-mer representations. A logistic regression model was used to create the bag-of-words representation ("bag-of-k-mers"), and a recurrent neural network was used to train the "vector-k-mers" representation. Although the accuracies of both models are satisfactory (over 90%), bag-of-k-mers outperformed vector-k-mers model.

Ng[76] developed a DNA sequence representation model and demonstrated the stability of their model empirically. The model is named dna2vec. The author used word2vec with the skip-gram algorithm to train the model, using a three-step procedure. The first step is dividing sequences into long non-overlapping fragments using gap characters (such as "X", "-" and etc.). This approach was also used in phylogenetic analysis[77]. The second step is converting long fragments to overlapping variable-length k-mers. In the third step, skip-gram and negative sampling is applied to train the model. Two different tasks were used to prove the stability of the trained representation model. The first one examined whether the summation of two k-mer sequence vectors was equal to the vector of the concatenated sequence. The second one was searching for a correlation between similar k-mers and the global alignment. For both tasks, the developed method was deemed successful.

Choi *et al.* trained a model named G2Vec[78] using random paths of functional interaction networks as sentences and genes as words. G2Vec, which is based on the continuous bag-of-words (CBOW) algorithm, was utilized for finding prognostic genes. The study showed that genes that imply good and bad prognosis could be separated in the representation space, and it can be used to discover new biomarkers.

### References


1. Klein, P., Kanehisa, M. & DeLisi, C. The detection and classification of membrane-spanning proteins. *Biochim. Biophys. Acta* 815, 468–476 (1985).

2. Klein, P., Jacquez, J. A. & Delisi, C. Prediction of protein function by discriminant analysis. *Math. Biosci.* 81, 177–189 (1986).

3. Liao, L. & Noble, W. S. Combining pairwise sequence similarity and support vector machines for detecting remote protein evolutionary and structural relationships. *J. Comput. Biol.* 10, 857–868 (2003).

4. Chen, Z. *et al.* iFeature: a Python package and web server for features extraction and selection from protein and peptide sequences. *Bioinformatics* 34, 2499–2502 (2018).

5. Wang, J. *et al.* POSSUM: a bioinformatics toolkit for generating numerical sequence feature descriptors based on PSSM profiles. *Bioinformatics* 33, 2756–2758 (2017).

6. Jiang, Y. *et al.* An expanded evaluation of protein function prediction methods shows an improvement in accuracy. *Genome Biol.* 17, 184 (2016).

7. Bengio, Y., Courville, A. & Vincent, P. Representation Learning: A Review and New Perspectives. *arXiv [cs.LG]* (2012).

8. Chen, R. T. Q., Li, X., Grosse, R. B. & Duvenaud, D. K. Isolating Sources of Disentanglement in Variational Autoencoders. in *Advances in Neural Information Processing Systems 31* (eds. Bengio, S. *et al.*) 2610–2620 (Curran Associates, Inc., 2018).

9. Achille, A. *et al.* Life-Long Disentangled Representation Learning with Cross-Domain Latent Homologies. in *Advances in Neural Information Processing Systems 31* (eds. Bengio, S. *et al.*) 9873–9883 (Curran Associates, Inc., 2018).

10. Jain, S., Banner, E., van de Meent, J.-W., Marshall, I. J. & Wallace, B. C. Learning Disentangled Representations of Texts with Application to Biomedical Abstracts. *arXiv [cs.CL]* (2018).

11. Tubiana, J., Cocco, S. & Monasson, R. Learning protein constitutive motifs from sequence data. *Elife* 8, (2019).

12. Andreeva, A., Kulesha, E., Gough, J. & Murzin, A. G. The SCOP database in 2020: expanded classification of representative family and superfamily domains of known protein structures. *Nucleic Acids Res.* 48, D376–D382 (2020).

13. Sillitoe, I. *et al.* CATH: expanding the horizons of structure-based functional annotations for genome sequences. *Nucleic Acids Res.* 47, D280–D284 (2019).

14. Yang, K. K., Wu, Z., Bedbrook, C. N. & Arnold, F. H. Learned protein embeddings for machine learning. *Bioinformatics* 34, 2642–2648 (2018).

15. Heinzinger, M. *et al.* Modeling the language of life – Deep Learning Protein Sequences. *Bioinformatics* 540 (2019).

16. Kim, S., Lee, H., Kim, K. & Kang, J. Mut2Vec: distributed representation of cancerous mutations. *BMC Med. Genomics* 11, 33 (2018).

17. Du, J. *et al.* Gene2vec: distributed representation of genes based on co-expression. *BMC Genomics* 20, 82 (2019).

18. Choy, C. T., Wong, C. H. & Chan, S. L. Infer related genes from large scale gene expression dataset with embedding. *Cancer Biology* 2524 (2018).

19. Asgari, E. & Mofrad, M. R. K. Continuous Distributed Representation of Biological Sequences for Deep Proteomics and Genomics. *PLoS One* 10, e0141287 (2015).

20. Rao, R. *et al.* Evaluating Protein Transfer Learning with TAPE. *arXiv [cs.LG]* (2019).

21. Chou, K.-C. Using amphiphilic pseudo amino acid composition to predict enzyme subfamily classes. *Bioinformatics* 21, 10–19 (2005).

22. Rives, A. *et al.* Biological structure and function emerge from scaling unsupervised learning to 250 million protein sequences. *Synthetic Biology* 7 (2019).

23. Leinonen, R. *et al.* UniProt archive. *Bioinformatics* 20, 3236–3237 (2004).

24. Melvin, I., Weston, J., Noble, W. S. & Leslie, C. Detecting remote evolutionary relationships among proteins by large-scale semantic embedding. *PLoS Comput. Biol.* 7, e1001047 (2011).

25. Qi, Y., Oja, M., Weston, J. & Noble, W. S. A unified multitask architecture for predicting local protein properties. *PLoS One* 7, e32235 (2012).

26. Collobert, R. & Weston, J. A unified architecture for natural language processing: deep neural networks with multitask learning. in *Proceedings of the 25th international conference on Machine learning* 160–167 (Association for Computing Machinery, 2008).

27. Kimothi, D., Soni, A., Biyani, P. & Hogan, J. M. Distributed Representations for Biological Sequence Analysis. *arXiv [cs.LG]* (2016).

28. Le, Q. & Mikolov, T. Distributed Representations of Sentences and Documents. in 1188–1196 (PMLR, 2014).

29. Asgari, E., McHardy, A. & Mofrad, M. R. K. Probabilistic variable-length segmentation of protein sequences for discriminative motif discovery (DiMotif) and sequence embedding (ProtVecX). *Bioinformatics* 707 (2018).

30. Gage, P. A new algorithm for data compression. *C Users J.* 12, 23–38 (1994).

31. Xu, Y., Song, J., Wilson, C. & Whisstock, J. C. PhosContext2vec: a distributed representation of residue-level sequence contexts and its application to general and kinase-specific phosphorylation site prediction. *Sci. Rep.* 8, 8240 (2018).

32. Cortes, C. & Vapnik, V. Support-vector networks. *Mach. Learn.* 20, 273–297 (1995).

33. Schwartz, A. S. *et al.* Deep Semantic Protein Representation for Annotation, Discovery, and Engineering. *Bioinformatics* D36 (2018).

34. Finn, R. D. *et al.* The Pfam protein families database. *Nucleic Acids Res.* 36, D281–8 (2008).

35. Hunter, S. *et al.* InterPro: the integrative protein signature database. *Nucleic Acids Res.* 37, D211–5 (2009).

36. Bairoch, A. The ENZYME database in 2000. *Nucleic Acids Res.* 28, 304–305 (2000).

37. The Gene Ontology Consortium & The Gene Ontology Consortium. The Gene Ontology Resource: 20 years and still GOing strong. *Nucleic Acids Research* vol. 47 D330–D338 (2019).

38. Hulo, N. *et al.* The PROSITE database. *Nucleic Acids Res.* 34, D227–30 (2006).

39. Lin, Q., Liang, L., Huang, Y. & Jin, L. Learning to Generate Realistic Scene Chinese Character Images by Multitask Coupled GAN. in *Pattern Recognition and Computer Vision* 41–51 (Springer International Publishing, 2018).

40. Chen, D., Mak, B., Leung, C. & Sivadas, S. Joint acoustic modeling of triphones and trigraphemes by multi-task learning deep neural networks for low-resource speech recognition. in *2014 IEEE International Conference on Acoustics, Speech and Signal Processing (ICASSP)* 5592–5596 (ieeexplore.ieee.org, 2014).

41. Schulz, C., Eger, S., Daxenberger, J., Kahse, T. & Gurevych, I. Multi-Task Learning for Argumentation Mining in Low-Resource Settings. *arXiv [cs.CL]* (2018).

42. Kriventseva, E. V. *et al.* OrthoDB v10: sampling the diversity of animal, plant, fungal, protist, bacterial and viral genomes for evolutionary and functional annotations of orthologs. *Nucleic Acids Res.* 47, D807–D811 (2019).

43. Cohen, T., Widdows, D., Heiden, J. A. V., Gupta, N. T. & Kleinstein, S. H. Graded Vector Representations of Immunoglobulins Produced in Response to West Nile Virus. in *Quantum Interaction* (eds. de Barros, J. A., Coecke, B. & Pothos, E.) vol. 10106 135–148 (Springer International Publishing, 2017).

44. Levy, S. D. & Gayler, R. Vector Symbolic Architectures: A New Building Material for Artificial General Intelligence. in *Proceedings of the 2008 conference on Artificial General Intelligence 2008: Proceedings of the First AGI Conference* 414–418 (IOS Press, 2008).

45. Viehweger, A., Krautwurst, S., Parks, D. H., König, B. & Marz, M. An encoding of genome content for machine learning. *Genomics* 1533 (2019).

46. You, R. & Zhu, S. DeepText2Go: Improving large-scale protein function prediction with deep semantic text representation. in *2017 IEEE International Conference on Bioinformatics and Biomedicine (BIBM)* 42–49 (2017).

47. Lindberg, D. A. Internet access to the National Library of Medicine. *Eff. Clin. Pract.* 3, 256–260 (2000).

48. Asgari, E., Poerner, N., McHardy, A. C. & Mofrad, M. R. K. DeepPrime2Sec: Deep Learning for Protein Secondary Structure Prediction from the Primary Sequences. 705426 (2019) doi:10.1101/705426.

49. Faisal, M. R. *et al.* Improving Protein Sequence Classification Performance Using Adjacent and Overlapped Segments on Existing Protein Descriptors. *JBiSE* 11, 126–143 (2018).

50. Jaeger, S., Fulle, S. & Turk, S. Mol2vec: Unsupervised Machine Learning Approach with Chemical Intuition. *J. Chem. Inf. Model.* 58, 27–35 (2018).

51. Figueras, J. Morgan revisited. *J. Chem. Inf. Comput. Sci.* 33, 717–718 (1993).

52. Weininger, D., Weininger, A. & Weininger, J. L. SMILES. 2. Algorithm for generation of unique SMILES notation. *J. Chem. Inf. Comput. Sci.* 29, 97–101 (1989).

53. Mikolov, T., Sutskever, I., Chen, K., Corrado, G. & Dean, J. Distributed Representations of

Words and Phrases and their Compositionality. *arXiv [cs.CL]* (2013).

54. Kané, H., Coulibali, M., Abdalla, A. & Ajanoh, P. Augmenting protein network embeddings with sequence information. *Bioinformatics* 1080 (2019).

55. Strodthoff, N., Wagner, P., Wenzel, M. & Samek, W. UDSMProt: universal deep sequence models for protein classification. *Bioinformatics* 36, 2401–2409 (2020).

56. Merity, S., Keskar, N. S. & Socher, R. Regularizing and Optimizing LSTM Language Models. *arXiv [cs.CL]* (2017).

57. Wan, F. & Zeng, J. (michael). Deep learning with feature embedding for compound-protein interaction prediction. *Bioinformatics* e1004157 (2016).

58. Mendez, D. *et al.* ChEMBL: towards direct deposition of bioassay data. *Nucleic Acids Res.* 47, D930–D940 (2019).

59. Wishart, D. S. *et al.* DrugBank 5.0: a major update to the DrugBank database for 2018. *Nucleic Acids Res.* 46, D1074–D1082 (2018).

60. Öztürk, H., Özgür, A. & Ozkirimli, E. DeepDTA: deep drug-target binding affinity prediction. *Bioinformatics* 34, i821–i829 (2018).

61. Öztürk, H., Ozkirimli, E. & Özgür, A. WideDTA: prediction of drug-target binding affinity. *arXiv [q-bio.QM]* (2019).

62. Yao, Y., Du, X., Diao, Y. & Zhu, H. An integration of deep learning with feature embedding for protein-protein interaction prediction. *PeerJ* 7, e7126 (2019).

63. Zhang, D. & Kabuka, M. Multimodal deep representation learning for protein interaction identification and protein family classification. *BMC Bioinformatics* 20, 531 (2019).

64. Nguyen, S., Li, Z. & Shang, Y. Deep Networks and Continuous Distributed Representation of Protein Sequences for Protein Quality Assessment. in *2017 IEEE 29th International Conference on Tools with Artificial Intelligence (ICTAI)* 527–534 (IEEE, 2017).

65. Mirabello, C. & Wallner, B. rawMSA: End-to-end Deep Learning using raw Multiple Sequence Alignments. *PLoS One* 14, e0220182 (2019).

66. Cuff, J. A. & Barton, G. J. Evaluation and improvement of multiple sequence methods for protein secondary structure prediction. *Proteins* 34, 508–519 (1999).

67. Alley, E. C., Khimulya, G., Biswas, S., AlQuraishi, M. & Church, G. M. Unified rational protein engineering with sequence-only deep representation learning. *Synthetic Biology* e1005786 (2019).

68. Bepler, T. & Berger, B. Learning protein sequence embeddings using information from structure. *arXiv [cs.LG]* (2019).

69. Andreeva, A., Howorth, D., Chothia, C., Kulesha, E. & Murzin, A. G. SCOP2 prototype: a new approach to protein structure mining. *Nucleic Acids Res.* 42, D310–4 (2014).

70. Elnaggar, A. *et al.* ProtTrans: Towards Cracking the Language of Life's Code Through Self-Supervised Deep Learning and High Performance Computing. *arXiv [cs.LG]* (2020).

71. Ramanujan, V., Wortsman, M., Kembhavi, A., Farhadi, A. & Rastegari, M. What's Hidden in a Randomly Weighted Neural Network? *arXiv [cs.CV]* (2019).

72. Oubounyt, M., Louadi, Z., Tayara, H. & To Chong, K. Deep Learning Models Based on Distributed Feature Representations for Alternative Splicing Prediction. *IEEE Access* 6, 58826–58834 (2018).

73. Szegedy, C. *et al.* Going deeper with convolutions. in *2015 IEEE Conference on Computer Vision and Pattern Recognition (CVPR)* 1–9 (2015).

74. Dutta, A., Dubey, T., Singh, K. K. & Anand, A. SpliceVec: Distributed feature representations for splice junction prediction. *Computational Biology and Chemistry* vol. 74 434–441 (2018).

75. Mejía-Guerra, M. K. & Buckler, E. S. A k-mer grammar analysis to uncover maize regulatory architecture. *BMC Plant Biol.* 19, 103 (2019).

76. Ng, P. dna2vec: Consistent vector representations of variable-length k-mers. *arXiv [q-bio.QM]* (2017).

77. Simmons, M. P. & Ochoterena, H. Gaps as characters in sequence-based phylogenetic analyses. *Syst. Biol.* 49, 369–381 (2000).

78. Choi, J., Oh, I., Seo, S. & Ahn, J. G2Vec: Distributed gene representations for identification of cancer prognostic genes. *Sci. Rep.* 8, 13729 (2018).

79. Lin, D. & Others. An information-theoretic definition of similarity. in *Icml* vol. 98 296–304 (1998).

80. Cozzetto, D., Buchan, D. W. A., Bryson, K. & Jones, D. T. Protein function prediction by

massive integration of evolutionary analyses and multiple data sources. *BMC Bioinformatics* 14 Suppl 3, S1 (2013).

81. Lan, L., Djuric, N., Guo, Y. & Vucetic, S. MS-kNN: protein function prediction by integrating multiple data sources. *BMC Bioinformatics* 14 Suppl 3, S8 (2013).

82. Hawkins, T., Chitale, M., Luban, S. & Kihara, D. PFP: Automated prediction of gene ontology functional annotations with confidence scores using protein sequence data. *Proteins* 74, 566–582 (2009).

83. Cao, R. & Cheng, J. Integrated protein function prediction by mining function associations, sequences, and protein–protein and gene–gene interaction networks. *Methods* 93, 84–91 (2016).

84. Piovesan, D., Giollo, M., Leonardi, E., Ferrari, C. & Tosatto, S. C. E. INGA: protein function prediction combining interaction networks, domain assignments and sequence similarity. *Nucleic Acids Res.* 43, W134–40 (2015).

85. Oates, M. E. *et al.* D2P2: database of disordered protein predictions. *Nucleic Acids Res.* 41, D508–D516 (2012).

86. Youngs, N., Penfold-Brown, D., Drew, K., Shasha, D. & Bonneau, R. Parametric Bayesian priors and better choice of negative examples improve protein function prediction. *Bioinformatics* 29, 1190–1198 (2013).

87. Sasidharan, R., Nepusz, T., Swarbreck, D., Huala, E. & Paccanaro, A. GFam: a platform for automatic annotation of gene families. *Nucleic Acids Res.* 40, e152 (2012).

88. Van Landeghem, S. *et al.* Exploring Biomolecular Literature with EVEX: Connecting Genes through Events, Homology, and Indirect Associations. *Adv. Bioinformatics* 2012, 582765 (2012).

### Supplementary Figures

**(a)**

| | Hydrophobicity | Net Charge |
|---|---|---|
| Protein 1 | 0.10 | 0.50 |
| Protein 2 | 0.15 | 0.45 |
| Protein 3 | 0.13 | 0.20 |
| Protein 4 | 0.10 | 0.15 |
| Protein 5 | 0.40 | 0.48 |
| Protein 6 | 0.45 | 0.40 |
| Protein 7 | 0.47 | 0.30 |
| Protein 8 | 0.45 | 0.25 |

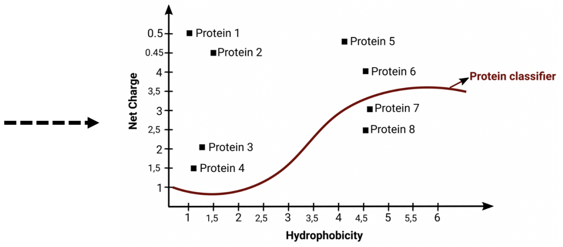


**(b)**

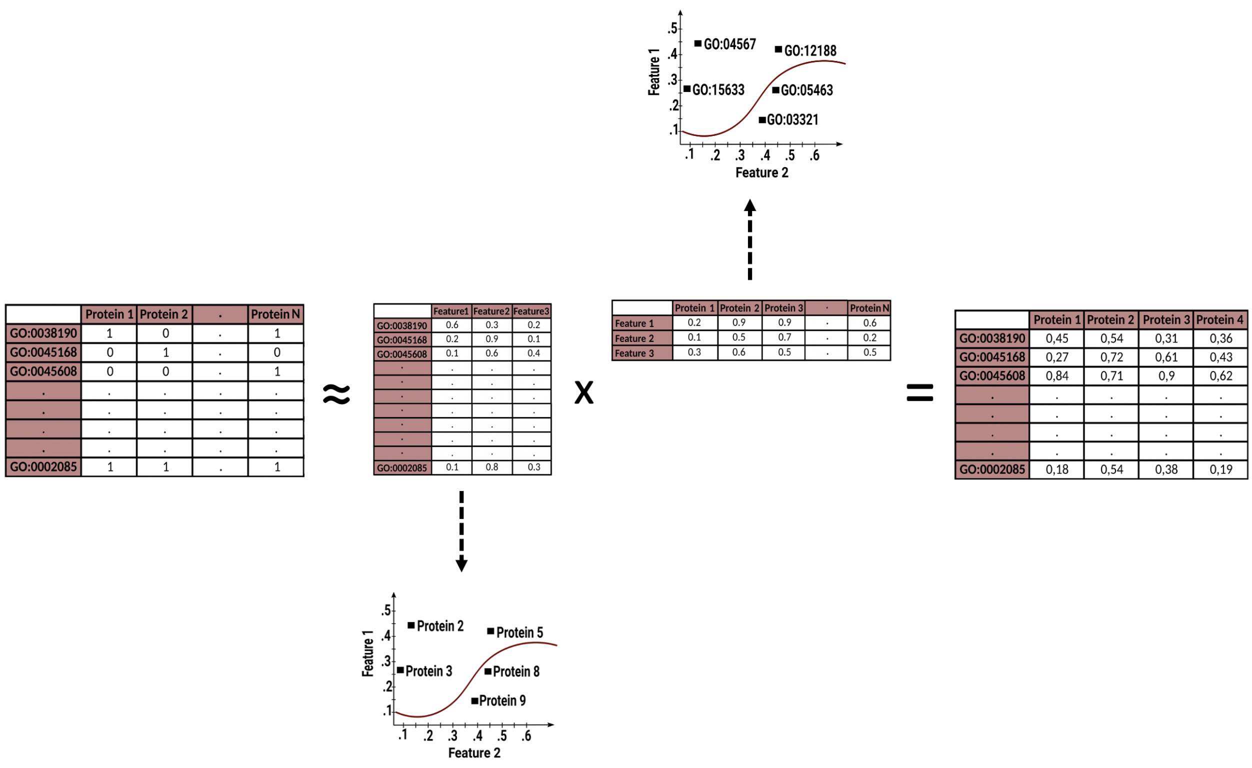


**Figure S1.** Examples for two different types of protein representation methods and their usage; **(a)** a classic protein representation example: Measurements regarding 2 different physicochemical properties (i.e., hydrophobicity and net charge) are used to represent each protein (left-side panel). 2-D protein representations are used as input to a second-order nonlinear logistic regression model, to classify the proteins into 2 groups (right-side panel); **(b)** a protein representation learning example: The source (i.e., training) data is a protein-GO term matrix displaying annotations. Feature vectors representing proteins and GO terms are calculated by training a model using a machine learning algorithm (e.g., matrix factorization with stochastic gradient descent) where the objective is the minimization of the difference between the real and predicted (i.e., dot product of the intermediate -latent- matrices) GO-protein associations. At the end of the training procedure, rows of the intermediate (latent) matrices of proteins and GO terms correspond to their representation vectors, which can later be used in various predictive tasks.

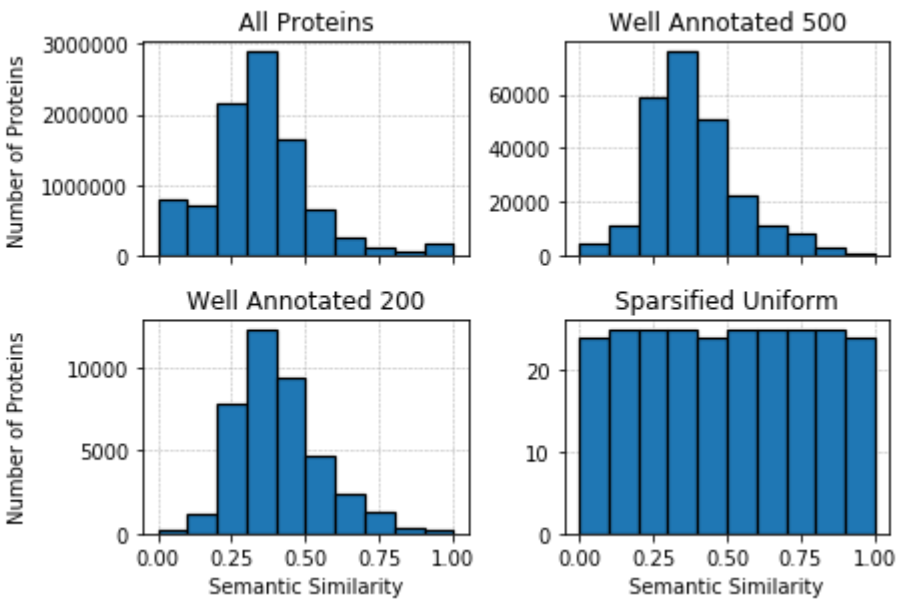


**Figure S2.** Distribution of pairwise protein similarities of GO terms before and after sparsification. Distribution of pairwise protein similarities for “all proteins”, “well annotated 500”, “well annotated 200”, and “sparse uniform” datasets for GO molecular function annotation dataset.

**(a)**

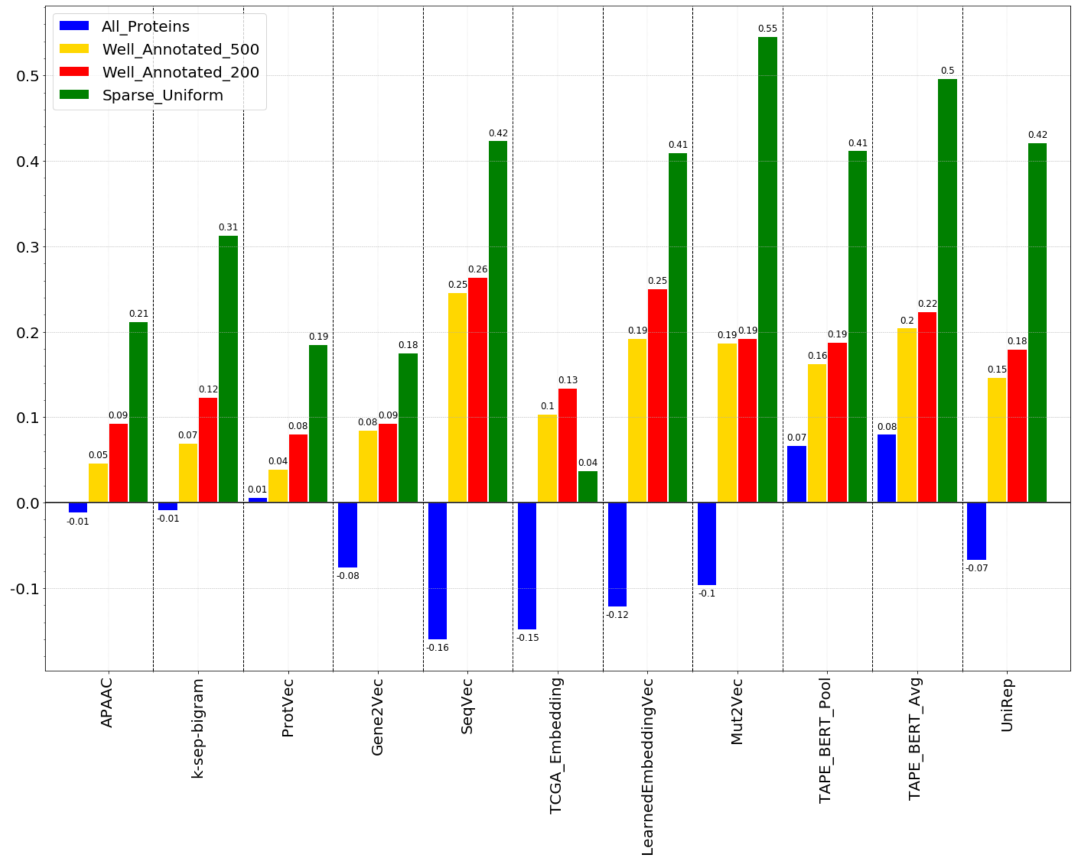
All_Proteins
Well_Annotated_500
Well_Annotated_200
Sparse_Uniform
0.5
0.4
0.3
0.2
0.1
0.0
-0.1
APAAC
k-sep-bigram
ProtVec
Gene2Vec
SeqVec
TCGA_Embedding
LearnedEmbeddingVec
Mut2Vec
TAPE_BERT_Pool
TAPE_BERT_Avg
UniRep

**(b)**

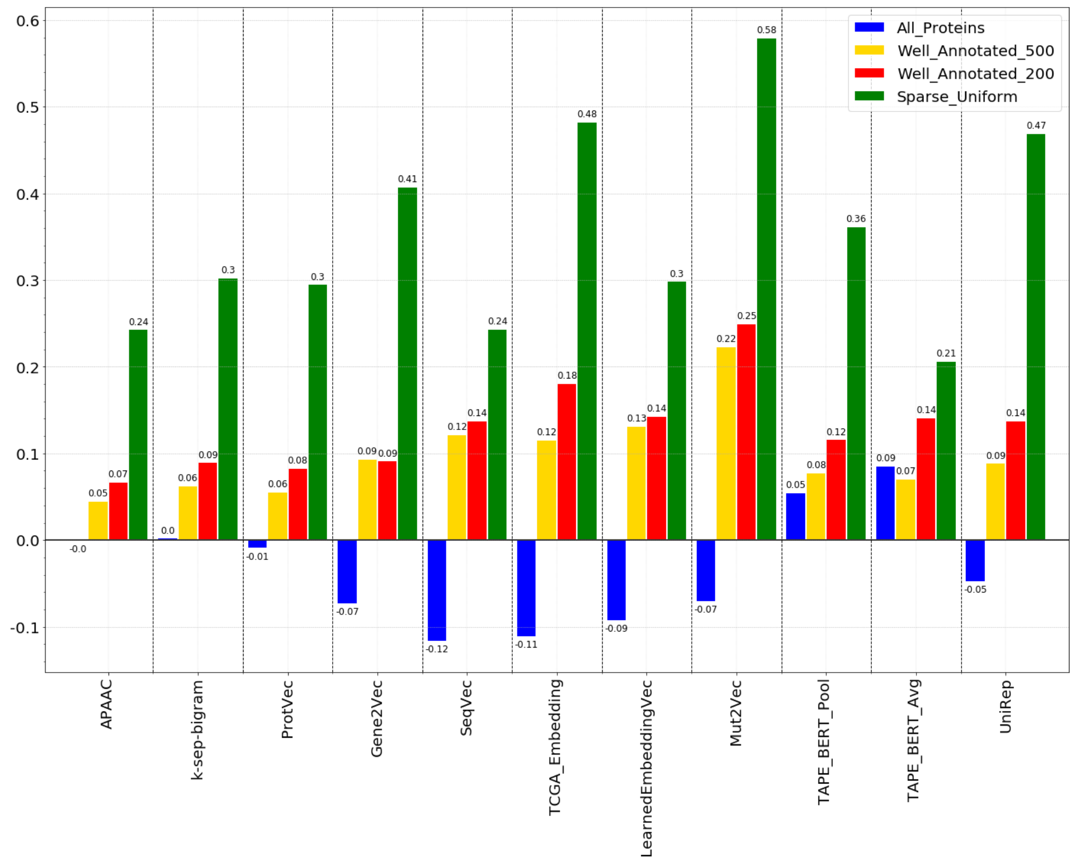
All_Proteins
Well_Annotated_500
Well_Annotated_200
Sparse_Uniform
0.6
0.5
0.4
0.3
0.2
0.1
0.0
-0.1
APAAC
k-sep-bigram
ProtVec
Gene2Vec
SeqVec
TCGA_Embedding
LearnedEmbeddingVec
Mut2Vec
TAPE_BERT_Pool
TAPE_BERT_Avg
UniRep

**(c)**

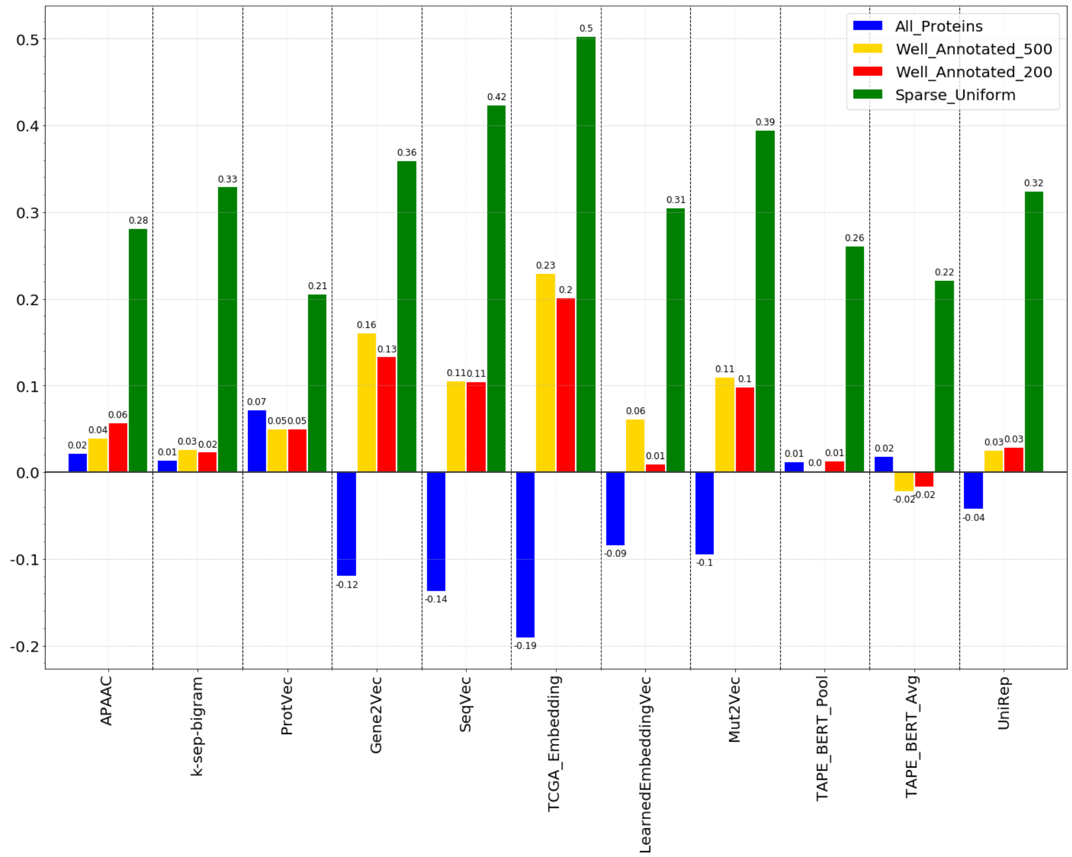


**Figure S3.** Performance of protein representation learning methods in inferring pairwise semantic similarities between proteins, calculated in terms of Spearman correlation between the ranked true pairwise similarity list (calculated using Lin similarities[79] between ontology-based functional annotations of proteins) and the representation-based ranked pairwise similarity list (calculated using “1 - normalized Manhattan distance” between numerical feature vectors of proteins). True semantic similarities are calculated based on GO terms of; **(a)** molecular function, **(b)** biological process, and **(c)** cellular component categories.

**(a)**

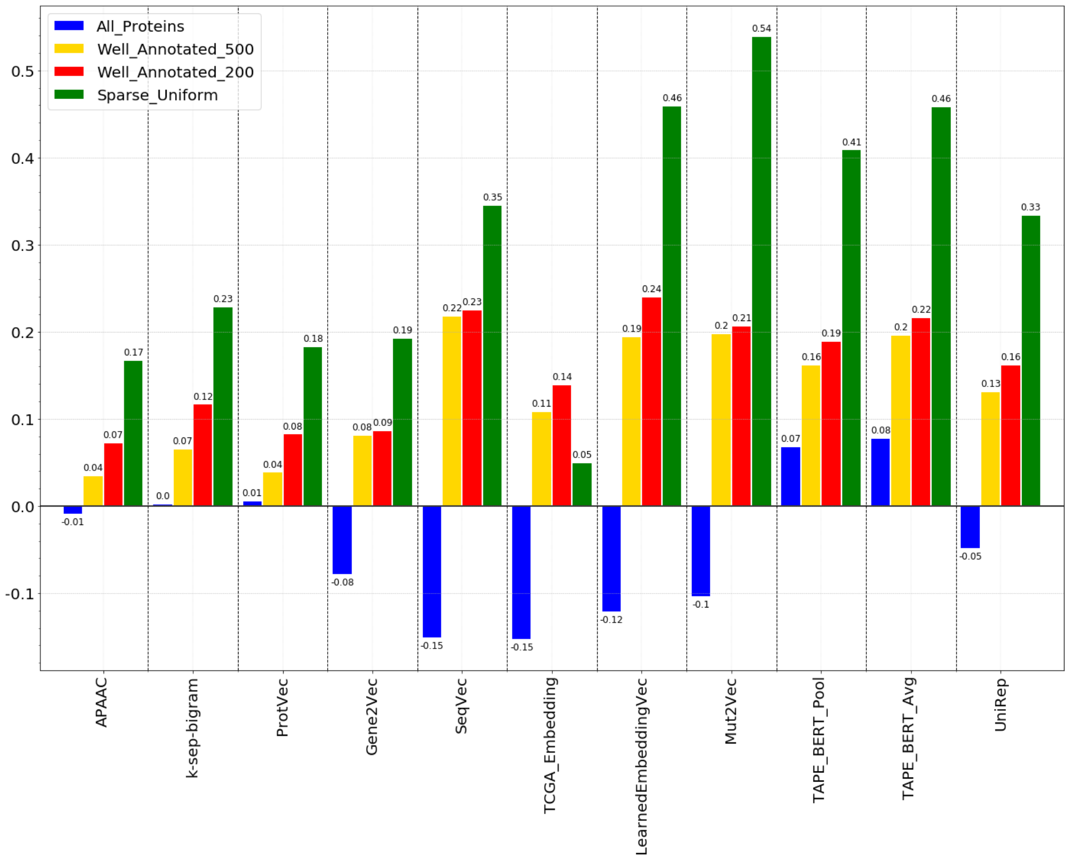
All_Proteins
Well_Annotated_500
Well_Annotated_200
Sparse_Uniform
APAAC
k-sep-bigram
ProtVec
Gene2Vec
SeqVec
TCGA_Embedding
LearnedEmbeddingVec
Mut2Vec
TAPE_BERT_Pool
TAPE_BERT_Avg
UniRep

**(b)**

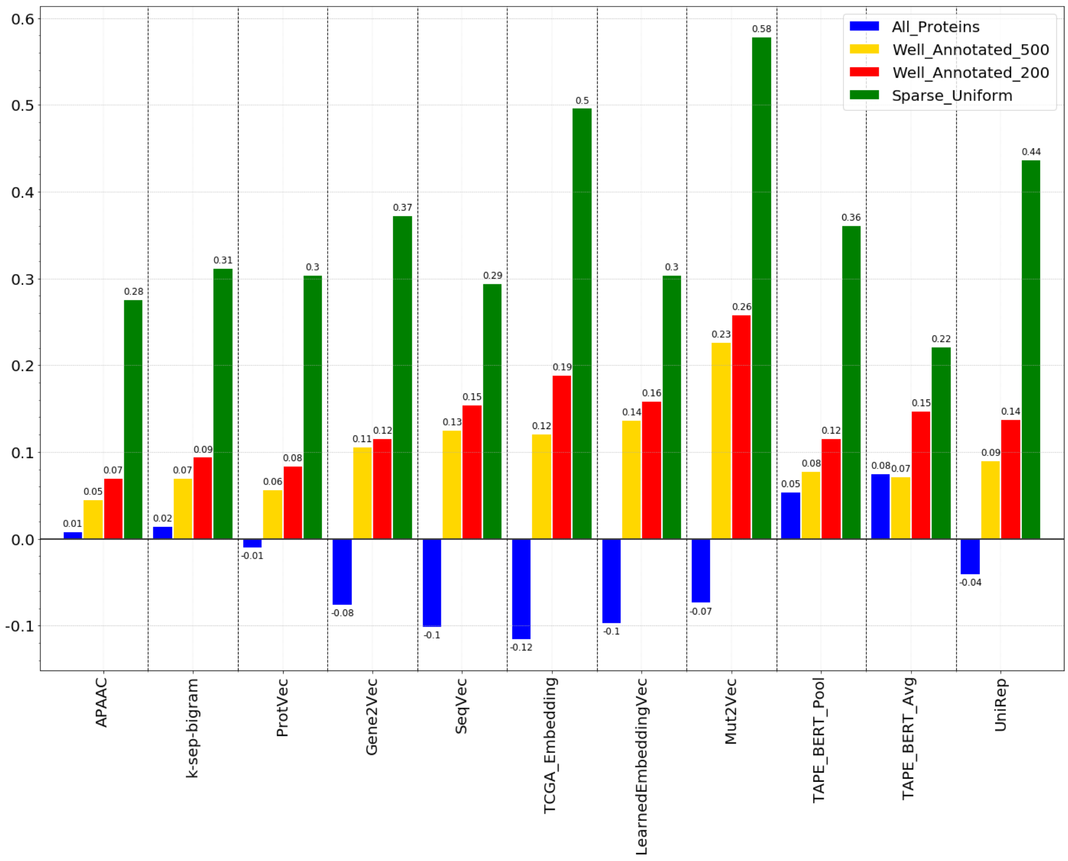
All_Proteins
Well_Annotated_500
Well_Annotated_200
Sparse_Uniform
APAAC
k-sep-bigram
ProtVec
Gene2Vec
SeqVec
TCGA_Embedding
LearnedEmbeddingVec
Mut2Vec
TAPE_BERT_Pool
TAPE_BERT_Avg
UniRep

**(c)**

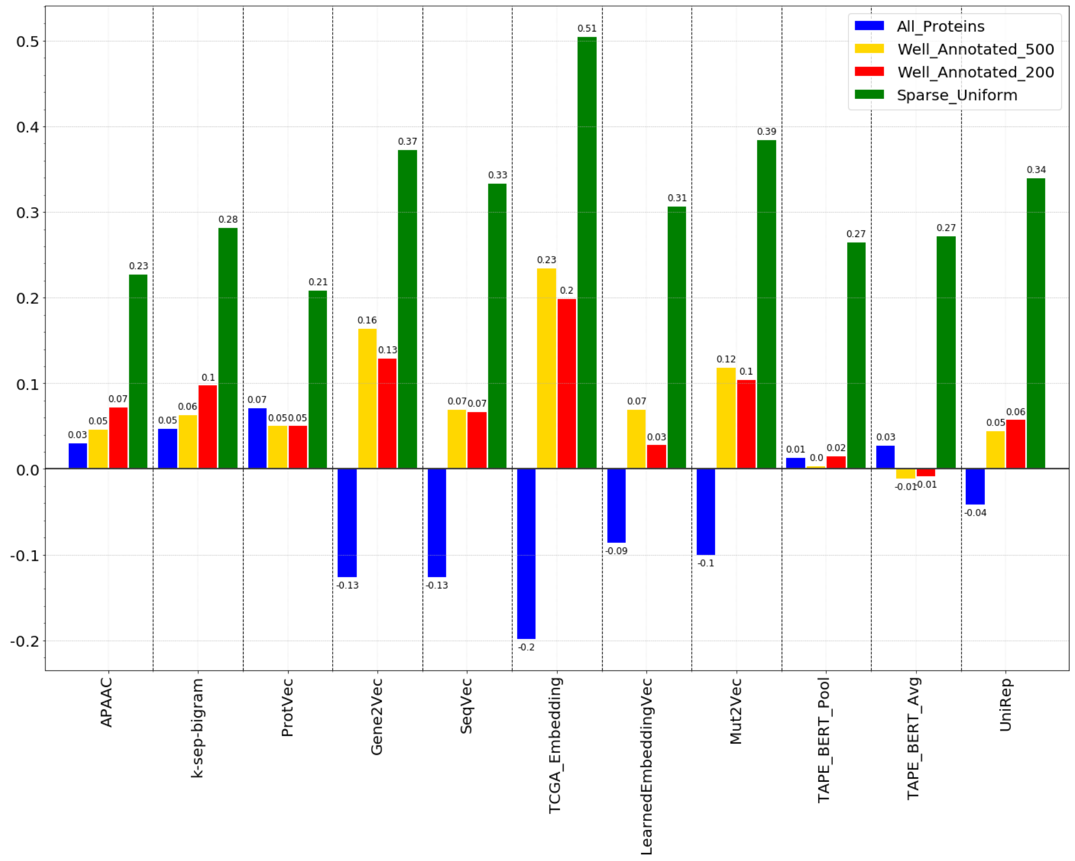


**Figure S4.** Performance of protein representation learning methods in inferring pairwise semantic similarities between proteins, calculated in terms of Spearman correlation between the ranked true pairwise similarity list (calculated using Lin similarities[79] between ontology-based functional annotations of proteins) and the representation-based ranked pairwise similarity list (calculated using "1 - Euclidean distance" between numerical feature vectors of proteins). True semantic similarities are calculated based on GO terms of; **(a)** molecular function, **(b)** biological process, and **(c)** cellular component categories.

**(a)**

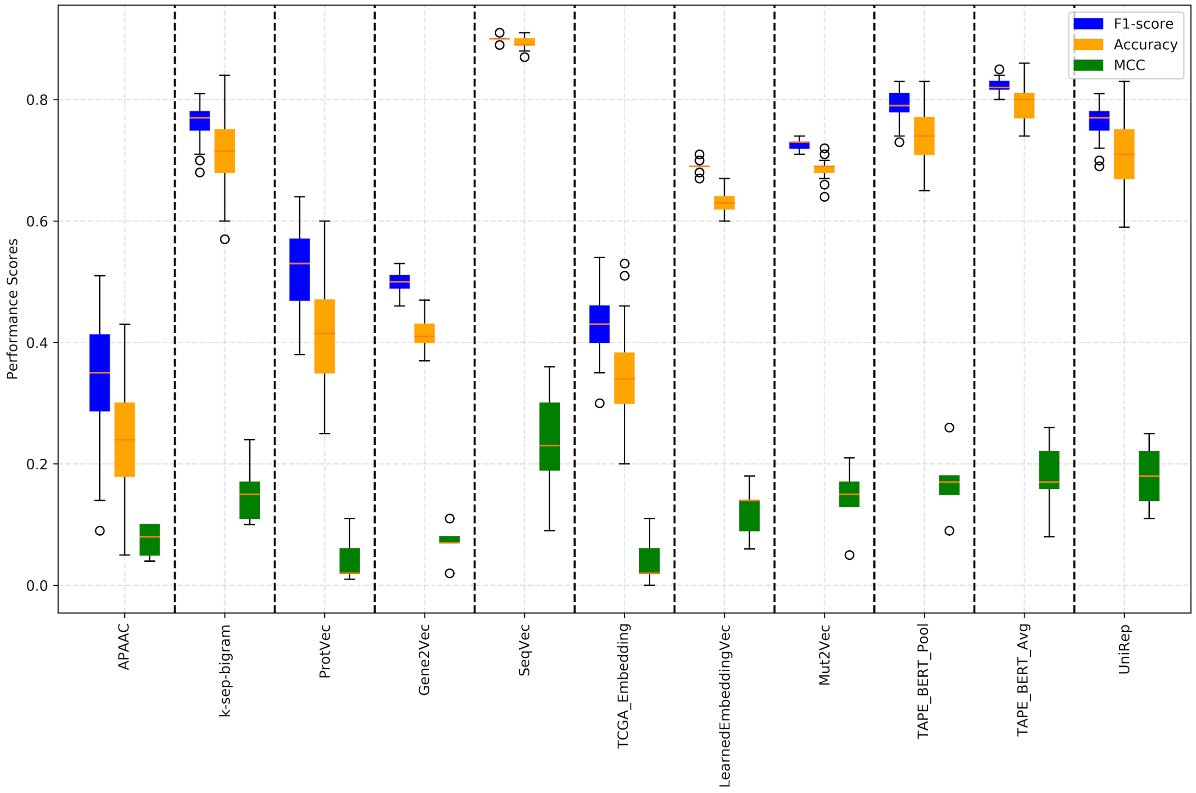

F1-score
Accuracy
MCC
Performance Scores
0.0
0.2
0.4
0.6
0.8
APAAC
k-sep-bigram
ProtVec
Gene2Vec
SeqVec
TCGA_Embedding
LearnedEmbeddingVec
Mut2Vec
TAPE_BERT_Pool
TAPE_BERT_Avg
UniRep


**(b)**

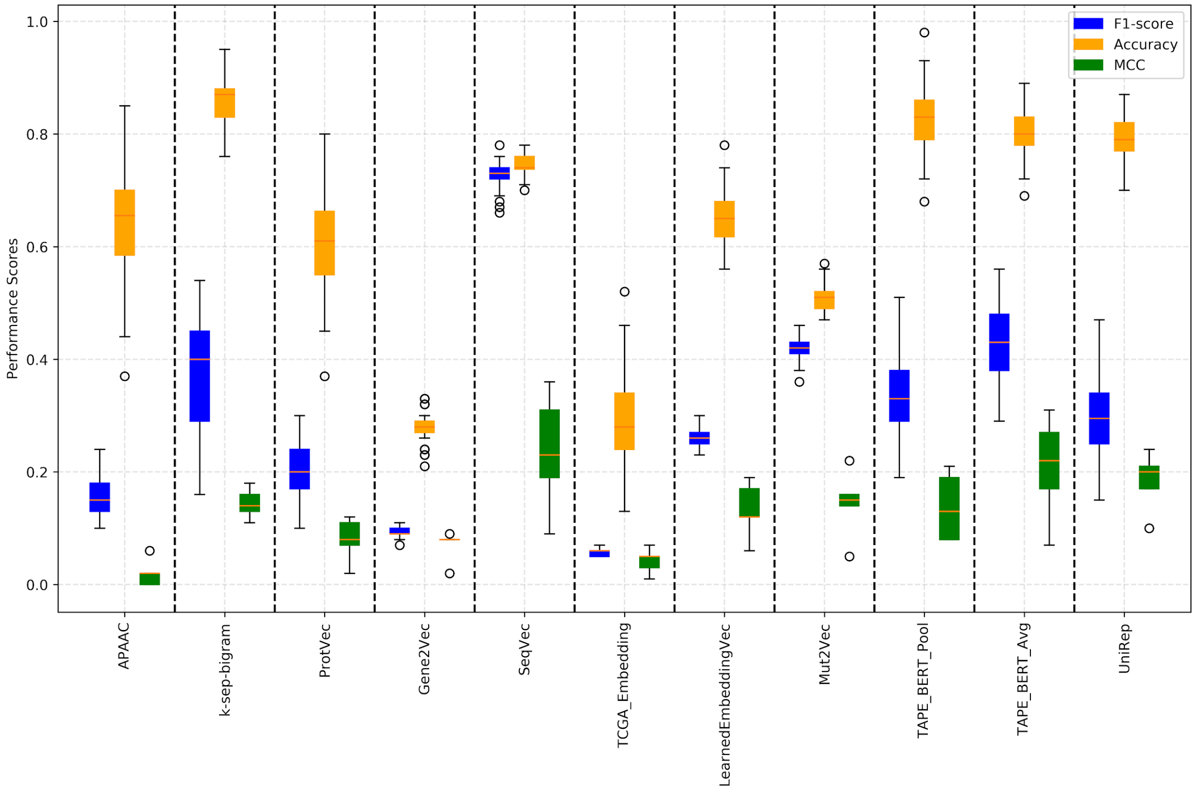

F1-score
Accuracy
MCC
Performance Scores
0.0
0.2
0.4
0.6
0.8
1.0
APAAC
k-sep-bigram
ProtVec
Gene2Vec
SeqVec
TCGA_Embedding
LearnedEmbeddingVec
Mut2Vec
TAPE_BERT_Pool
TAPE_BERT_Avg
UniRep

(c)

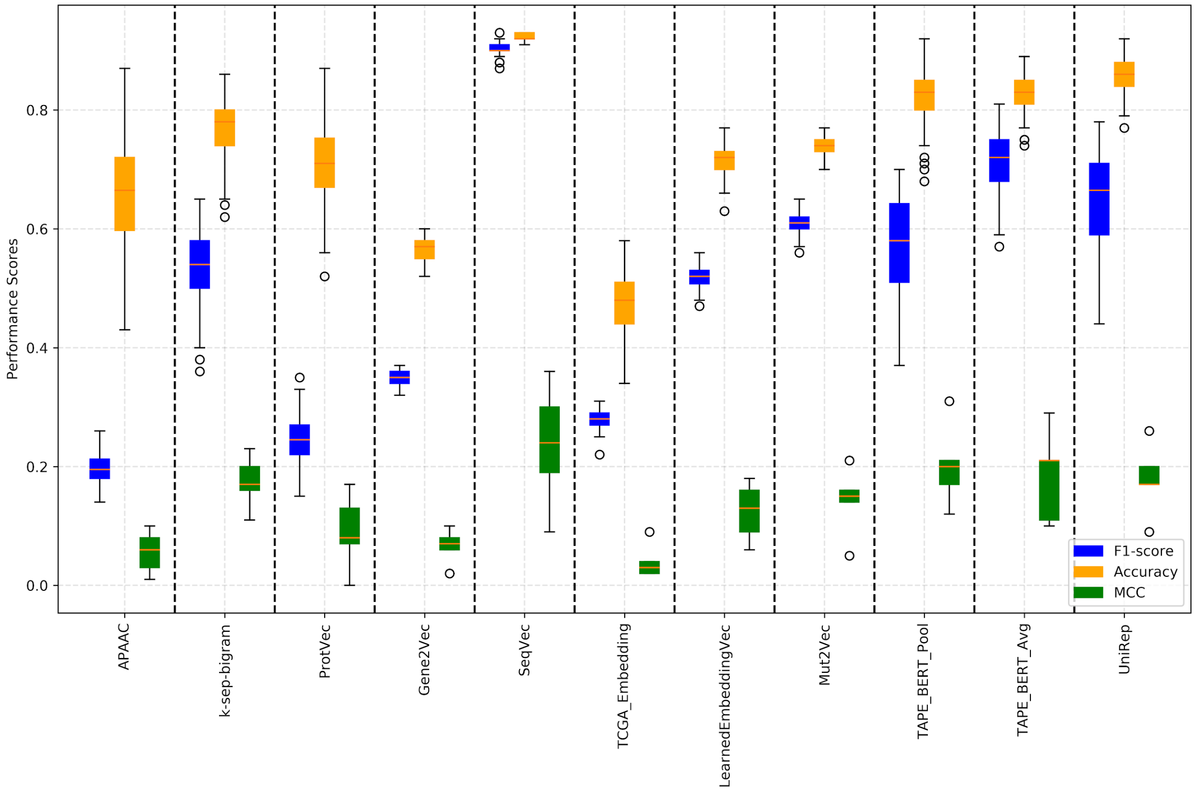

Performance Scores
0.0
0.2
0.4
0.6
0.8
F1-score
Accuracy
MCC
APAAC
k-sep-bigram
ProtVec
Gene2Vec
SeqVec
TCGA_Embedding
LearnedEmbeddingVec
Mut2Vec
TAPE_BERT_Pool
TAPE_BERT_Avg
UniRep


(d)

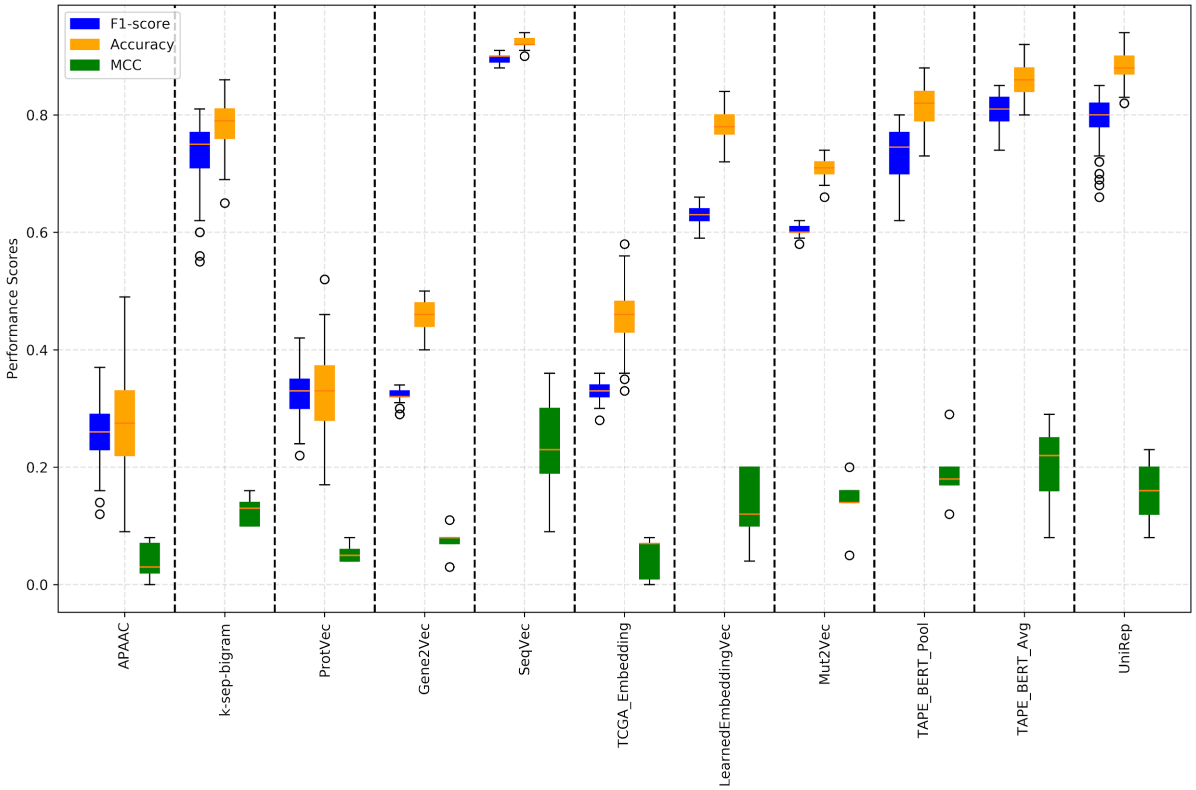

Performance Scores
0.0
0.2
0.4
0.6
0.8
F1-score
Accuracy
MCC
APAAC
k-sep-bigram
ProtVec
Gene2Vec
SeqVec
TCGA_Embedding
LearnedEmbeddingVec
Mut2Vec
TAPE_BERT_Pool
TAPE_BERT_Avg
UniRep

**(e)**

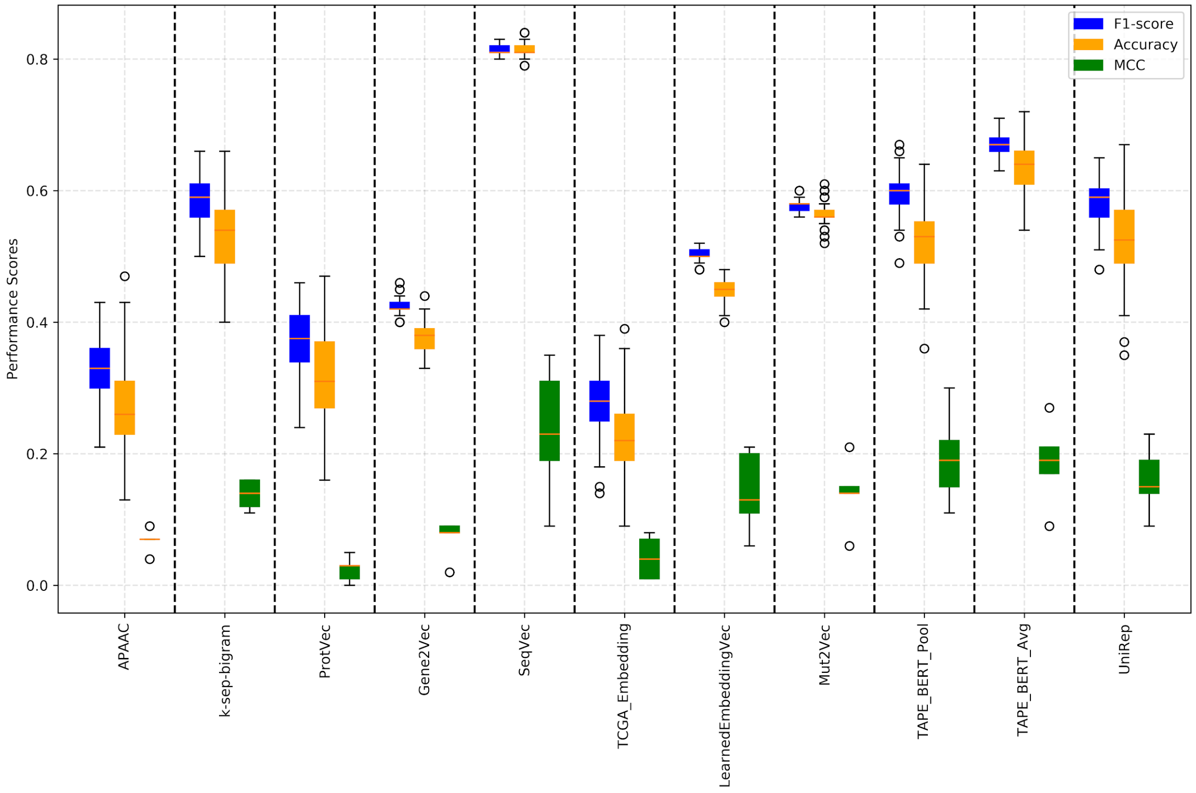


**Figure S5.** Protein family-based box plots indicating performance results (F1-score, accuracy and MCC) of protein representation learning methods in the drug target protein family classification benchmark, based on families; **(a)** enzymes, **(b)** membrane receptors, **(c)** transcription factors, **(d)** ion channels, and **(e)** others.

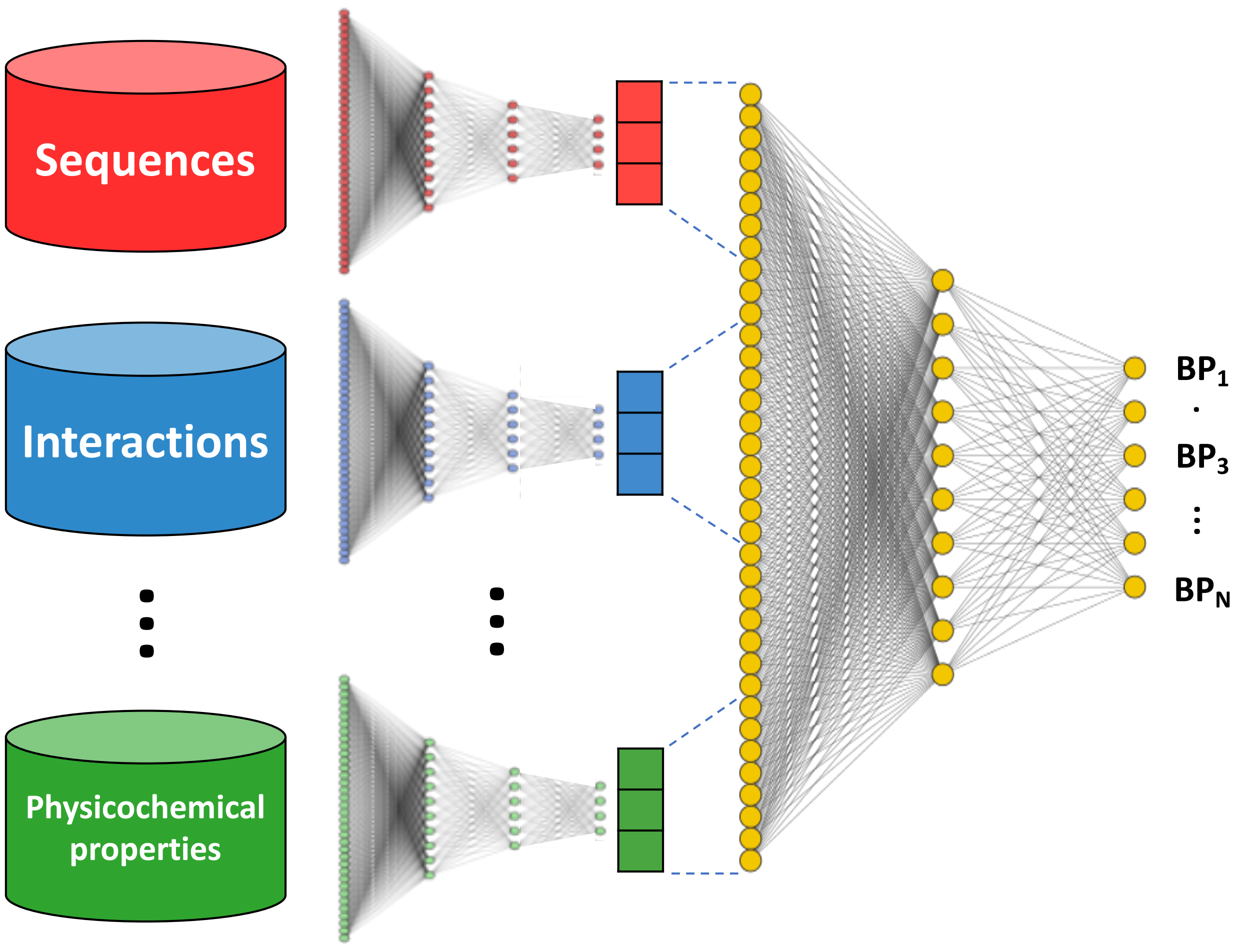


**Figure S6.** Schematic representation of an example ensemble-based protein representation vector construction model. Individual protein representation models are trained separately on different protein data (e.g., sequence, PPIs, physicochemical properties and etc.) and a holistic protein representation is trained on the integration of the vectorial output of these models. The holistic representation is trained using high-level tasks in a supervised manner, such as the prediction of biological processes that the input proteins take part in.

### Supplementary Tables

**Table S1.** Input data type and featurization approaches in classical protein representations used in the CAFA2 participating protein function prediction methods.

| Input Data Type | Method | Utilized Feature |
|---|---|---|
| Protein Sequence | Cozzetto *et al.*[80] | Residue composition, Sequence length |
| | Lan *et al.*[81] | Identity based similarity score |
| | Hawkins *et al.*[82] | Identity based similarity score |
| | Cao and Cheng[83] | Identity based similarity score, E-value based probabilistic confidence score |
| | Piovesan *et al.*[84] | Identity-based similarity score |
| Disordered regions | Cozzetto *et al.*[80] | Number of disordered regions |
| | Oates *et al.*[85] | One-hot encoding of disordered regions |
| Protein-protein interaction | Lan *et al.*[81] | Similarity score based on binary interaction vectors. |
| | Youngs *et al.*[86] | Interaction weighting matrices |
| | Piovesan *et al.*[84]} | Binary interaction vectors |
| Gene expression | Lan *et al.*[81] | Expression based similarity score |
| Functional annotations | Sasidharan *et al.*[87] | Domain annotation list |
| | Cozzetto *et al.*[80] | Number of transmembrane residues, Number of low complexity regions, Number of glycosylated residues, Localisation existence, Signal Peptide length, Signal Peptide score |
| Literature texts | Van Landeghem *et al.*[88] | Named entity tags |
| Secondary structures | Cozzetto *et al.*[80] | Number of helices, sheets, coils |
| Physicochemical properties | Cozzetto *et al.*[80] | Molecular weight, Average hydrophobicity, Charge, Molar extinction coefficient, Isoelectric point, Aliphatic index |

**Table S2.** The number of proteins for each GO category in the dataset used in the protein function prediction benchmark.

| | | | Grouping GO terms in terms of the number of annotations | | |
|---|---|---|---|---|---|
| | | | **High** | **Middle** | **Low** |
| **Grouping GO terms considering the specificity of the term (i.e., location on the hierarchy of GO DAG)** | **Shallow** | **BP** | 3,386 | 896 | 27 |
| | | **MF** | 3,039 | 613 | 41 |
| | | **CC** | 7,186 | 413 | 55 |
| | **Normal** | **BP** | 3,288 | 452 | 28 |
| | | **MF** | 1,324 | 369 | 31 |
| | | **CC** | 6,478 | 562 | 47 |
| | **Specific** | **BP** | 2,025 | 623 | 42 |
| | | **MF** | 0 | 204 | 41 |
| | | **CC** | 0 | 413 | 54 |

**Table S3.** Identifiers of GO terms that are incorporated into each of the 25 different multi-task models constructed for the protein function prediction benchmark.

| | | | Grouping GO terms in terms of the number of annotations | | |
|---|---|---|---|---|---|
| | | | **High** | **Middle** | **Low** |
| **Grouping GO terms considering the specificity of the term (i.e. location on the hierarchy of GO DAG)** | **Shallow** | **BP** | GO:0007399<br>GO:0006259<br>GO:0007167<br>GO:0006886<br>GO:0051707 | GO:0043488<br>GO:0043312<br>GO:0038096<br>GO:0006637<br>GO:0051091 | GO:0032933<br>GO:0006348<br>GO:0042989<br>GO:0000054<br>GO:0039529 |
| | | **MF** | GO:0016818<br>GO:0022890<br>GO:0046872<br>GO:0004672<br>GO:0000981 | GO:0036459<br>GO:0046943<br>GO:0005524<br>GO:0005244<br>GO:0022835 | GO:0015187<br>GO:0008568<br>GO:0008508<br>GO:0005328<br>GO:0043138 |
| | | **CC** | GO:0005789<br>GO:1990234<br>GO:1903561<br>GO:0005740<br>GO:0070013 | GO:0008076<br>GO:0005762<br>GO:0005747<br>GO:0036464<br>GO:0044853 | GO:0000124<br>GO:0097165<br>GO:0008024<br>GO:0032590<br>GO:0033162 |
| | **Normal** | **BP** | GO:0070647<br>GO:0016071<br>GO:0015833<br>GO:0048699<br>GO:0019752 | GO:0071427<br>GO:0016573<br>GO:0006613<br>GO:0031146<br>GO:0051092 | GO:0098703<br>GO:0061740<br>GO:0014808<br>GO:0000056<br>GO:0050482 |
| | | **MF** | GO:0016462<br>GO:0044212 | GO:0003774<br>GO:0008227<br>GO:0004866<br>GO:0004714<br>GO:0004843 | GO:0016934<br>GO:0022848<br>GO:0005314<br>GO:0003689<br>GO:0005335 |
| | | **CC** | GO:0070062<br>GO:0000228<br>GO:0031981<br>GO:0015630<br>GO:0005768 | GO:0000502<br>GO:0005925<br>GO:0022627<br>GO:0030665<br>GO:0034705 | GO:0036020<br>GO:0070081<br>GO:0044754<br>GO:1990454<br>GO:0031089 |
| | **Specific** | **BP** | GO:0045944<br>GO:0001934<br>GO:1903507 | GO:1903169<br>GO:0050773<br>GO:0071805<br>GO:0031124<br>GO:0000209 | GO:0043984<br>GO:0034625<br>GO:0006231<br>GO:0046323<br>GO:0072539 |
| | | **MF** | - | GO:0061733<br>GO:0003777<br>GO:0004386 | GO:0004957<br>GO:0008511<br>GO:0004571<br>GO:0017049<br>GO:0097199 |
| | | **CC** | - | GO:0005766<br>GO:0016591<br>GO:0101002<br>GO:0008021 | GO:0019908<br>GO:0005767<br>GO:0032009<br>GO:0005736<br>GO:0099061 |

**Table S4.** Overall performance results in ontology-based protein function prediction benchmark for: **(a)** GO molecular function, **(b)** GO biological process, **(c)** GO cellular component.

**(a)**

| **MF** | **APAAC** | **k-sep-bigram** | **ProtVec** | **Gene2Vec** | **SeqVec** | **TCGA_Embedding** | **LearnedEmbeddingVec** | **Mut2Vec** | **TAPE-BERT_Pool** | **TAPE-BERT_Avg** | **UniRep** |
|---|---|---|---|---|---|---|---|---|---|---|---|
| **Recall** | 0.627 | 0.888 | 0.701 | 0.525 | **0.908** | 0.174 | 0.596 | 0.468 | 0.844 | 0.869 | 0.868 |
| **Precision** | 0.755 | 0.903 | 0.745 | 0.582 | **0.929** | 0.235 | 0.747 | 0.733 | 0.903 | 0.890 | 0.862 |
| **F1-score (weighted)** | 0.648 | 0.890 | 0.703 | 0.534 | **0.915** | 0.173 | 0.632 | 0.541 | 0.862 | 0.874 | 0.860 |
| **Accuracy** | 0.557 | 0.842 | 0.598 | 0.468 | **0.895** | 0.166 | 0.584 | 0.461 | 0.816 | 0.832 | 0.803 |
| **Hamming dist. (avg.)** | 0.145 | 0.056 | 0.142 | 0.191 | **0.039** | 0.239 | 0.133 | 0.155 | 0.062 | 0.058 | 0.065 |

**(b)**

| **BP** | **APAAC** | **k-sep-bigram** | **ProtVec** | **Gene2Vec** | **SeqVec** | **TCGA_Embedding** | **LearnedEmbeddingVec** | **Mut2Vec** | **TAPE-BERT_Pool** | **TAPE-BERT_Avg** | **UniRep** |
|---|---|---|---|---|---|---|---|---|---|---|---|
| **Recall** | 0.320 | 0.597 | 0.383 | 0.411 | **0.577** | 0.142 | 0.262 | 0.293 | 0.511 | 0.572 | 0.566 |
| **Precision** | 0.435 | 0.647 | 0.454 | 0.540 | **0.684** | 0.230 | 0.325 | 0.540 | 0.672 | 0.635 | 0.553 |
| **F1-score (weighted)** | 0.333 | 0.606 | 0.390 | 0.441 | **0.610** | 0.153 | 0.277 | 0.353 | 0.556 | 0.589 | 0.547 |
| **Accuracy** | 0.255 | 0.480 | 0.298 | 0.348 | **0.526** | 0.140 | 0.239 | 0.260 | 0.452 | 0.472 | 0.402 |
| **Hamming dist. (avg.)** | 0.219 | 0.173 | 0.228 | 0.202 | **0.146** | 0.220 | 0.200 | 0.186 | 0.173 | 0.177 | 0.208 |

**(c)**

| **CC** | **APAAC** | **k-sep-bigram** | **ProtVec** | **Gene2Vec** | **SeqVec** | **TCGA_Embedding** | **LearnedEmbeddingVec** | **Mut2Vec** | **TAPE-BERT_Pool** | **TAPE-BERT_Avg** | **UniRep** |
|---|---|---|---|---|---|---|---|---|---|---|---|
| **Recall** | 0.369 | 0.614 | 0.451 | 0.509 | **0.615** | 0.315 | 0.309 | 0.328 | 0.571 | 0.612 | 0.581 |
| **Precision** | 0.460 | 0.662 | 0.467 | 0.568 | **0.713** | 0.345 | 0.367 | 0.500 | 0.670 | 0.652 | 0.592 |
| **F1-score (weighted)** | 0.361 | 0.616 | 0.430 | 0.510 | **0.637** | 0.304 | 0.313 | 0.358 | 0.587 | 0.612 | 0.569 |
| **Accuracy** | 0.286 | 0.502 | 0.321 | 0.421 | **0.553** | 0.293 | 0.285 | 0.296 | 0.496 | 0.514 | 0.433 |
| **Hamming dist. (avg.)** | 0.228 | 0.163 | 0.228 | 0.179 | **0.140** | 0.182 | 0.181 | 0.174 | 0.156 | 0.157 | 0.190 |

**Table S5.** Average performance results (weighted F1-score) in terms of GO groups such as "low", "middle", "high", and "specific", "normal", "shallow", in ontology-based protein function prediction benchmark for: **(a)** GO molecular function, **(b)** GO biological process, **(c)** GO cellular component.

**(a)**

| MF | APAAC | k-sep-bigram | ProtVec | Gene2Vec | SeqVec | TCGA_Embedding | LearnedEmbeddingVec | Mut2Vec | TAPE_BERT_Pool | TAPE_BERT_Avg | UniRep | Mean of all methods |
|---|---|---|---|---|---|---|---|---|---|---|---|---|
| **Low** | 0.694 | **0.916** | 0.784 | 0.525 | **0.914** | 0.102 | 0.573 | 0.452 | 0.871 | 0.872 | 0.872 | 0.689 |
| **Middle** | 0.631 | 0.887 | 0.621 | 0.611 | **0.920** | 0.153 | 0.639 | 0.568 | 0.849 | 0.883 | 0.858 | 0.693 |
| **High** | 0.603 | 0.854 | 0.706 | 0.433 | **0.907** | 0.309 | 0.711 | 0.633 | 0.868 | 0.863 | 0.843 | 0.703 |
| **Specific** | 0.667 | 0.909 | 0.671 | 0.486 | **0.945** | 0.043 | 0.542 | 0.511 | 0.826 | 0.867 | 0.885 | 0.669 |
| **Normal** | 0.781 | **0.911** | 0.773 | 0.663 | **0.908** | 0.331 | 0.725 | 0.645 | 0.907 | 0.895 | 0.880 | 0.765 |
| **Shallow** | 0.502 | 0.855 | 0.655 | 0.437 | **0.901** | 0.101 | 0.599 | 0.456 | 0.841 | 0.857 | 0.822 | 0.639 |

**(b)**

| BP | APAAC | k-sep-bigram | ProtVec | Gene2Vec | SeqVec | TCGA_Embedding | LearnedEmbeddingVec | Mut2Vec | TAPE_BERT_Pool | TAPE_BERT_Avg | UniRep | Mean of all methods |
|---|---|---|---|---|---|---|---|---|---|---|---|---|
| **Low** | 0.452 | **0.548** | 0.413 | 0.416 | 0.455 | 0.063 | 0.210 | 0.138 | 0.480 | 0.524 | 0.456 | 0.378 |
| **Middle** | 0.346 | 0.689 | 0.472 | 0.527 | **0.739** | 0.282 | 0.345 | 0.436 | 0.652 | 0.673 | 0.650 | 0.528 |
| **High** | 0.201 | 0.582 | 0.287 | 0.381 | **0.637** | 0.115 | 0.275 | 0.484 | 0.536 | 0.571 | 0.535 | 0.419 |
| **Specific** | 0.466 | 0.681 | 0.536 | 0.501 | **0.732** | 0.182 | 0.490 | 0.445 | 0.655 | 0.689 | 0.639 | 0.547 |
| **Normal** | 0.219 | **0.566** | 0.245 | 0.427 | 0.532 | 0.125 | 0.124 | 0.287 | 0.505 | 0.520 | 0.485 | 0.367 |
| **Shallow** | 0.315 | **0.571** | 0.390 | 0.397 | **0.567** | 0.152 | 0.217 | 0.327 | 0.508 | 0.560 | 0.517 | 0.411 |

**(c)**

| CC | APAAC | k-sep-bigram | ProtVec | Gene2Vec | SeqVec | TCGA_Embedding | LearnedEmbeddingVec | Mut2Vec | TAPE_BERT_Pool | TAPE_BERT_Avg | UniRep | Mean of all methods |
|---|---|---|---|---|---|---|---|---|---|---|---|---|
| **Low** | 0.311 | **0.556** | 0.373 | 0.433 | **0.551** | 0.178 | 0.173 | 0.175 | 0.502 | 0.533 | 0.491 | 0.389 |
| **Middle** | 0.354 | 0.657 | 0.461 | 0.588 | **0.679** | 0.336 | 0.338 | 0.426 | 0.630 | 0.645 | 0.618 | 0.521 |
| **High** | 0.445 | 0.646 | 0.470 | 0.508 | **0.703** | 0.447 | 0.485 | 0.531 | 0.651 | 0.680 | 0.614 | 0.562 |
| **Specific** | 0.180 | **0.552** | 0.404 | 0.466 | 0.523 | 0.093 | 0.090 | 0.195 | 0.484 | 0.503 | 0.483 | 0.361 |
| **Normal** | 0.384 | 0.622 | 0.439 | 0.519 | **0.658** | 0.386 | 0.398 | 0.389 | 0.619 | 0.636 | 0.595 | 0.513 |
| **Shallow** | 0.458 | 0.654 | 0.438 | 0.529 | **0.692** | 0.364 | 0.375 | 0.436 | 0.624 | 0.660 | 0.601 | 0.530 |

**Table S6.** Sample statistics of the dataset used in drug target protein family prediction benchmark, per target protein family and representation model.

| Method | Enzymes | Membrane receptors | Transcription factors | Ion channels | Others | Total |
|---|---|---|---|---|---|---|
| Learned EmbeddingVec | 1,564 | 322 | 82 | 151 | 265 | 2,384 |
| SeqVec | 1,564 | 322 | 82 | 151 | 265 | 2,384 |
| Mut2Vec | 1,506 | 314 | 79 | 148 | 259 | 2,306 |
| Gene2Vec | 1,548 | 311 | 82 | 150 | 263 | 2,354 |
| TCGA_embedding | 1,556 | 322 | 82 | 151 | 262 | 2,373 |
| ProtVec | 1,564 | 322 | 82 | 151 | 265 | 2,384 |
| TAPE-BERT_avg | 1,544 | 321 | 81 | 138 | 251 | 2,335 |
| TAPE-BERT_pool | 1,544 | 321 | 81 | 138 | 251 | 2,335 |
| Unirep | 1,564 | 322 | 82 | 151 | 265 | 2,384 |
| APAAC | 1,564 | 322 | 82 | 151 | 265 | 2,384 |
| k-sep-bigrams | 1,559 | 322 | 82 | 150 | 264 | 2,377 |
